# Rational optimization of a transcription factor activation domain inhibitor

**DOI:** 10.1101/2022.08.18.504385

**Authors:** Shaon Basu, Paula Martínez-Cristóbal, Mireia Pesarrodona, Marta Frigolé-Vivas, Michael Lewis, Elzbieta Szulc, C. Adriana Bañuelos, Carolina Sánchez-Zarzalejo, Stasė Bielskutė, Jiaqi Zhu, Karina Pombo-García, Carla Garcia-Cabau, Cristina Batlle, Borja Mateos, Mateusz Biesaga, Albert Escobedo, Lídia Bardia, Xavier Verdaguer, Alessandro Ruffoni, Nasrin R. Mawji, Jun Wang, Teresa Tam, Isabelle Brun-Heath, Salvador Ventura, David Meierhofer, Jesús García, Paul Robustelli, Travis H. Stracker, Marianne D. Sadar, Antoni Riera, Denes Hnisz, Xavier Salvatella

**Author notes:** equally contributed (Shaon Basu, Paula Martínez-Cristóbal). equally contributed (Mireia Pesarrodona, Marta Frigolé-Vivas).

## Abstract

Transcription factors are among the most attractive therapeutic targets but are considered largely undruggable due to the intrinsically disordered nature of their activation domains. Here we show that the aromatic character of the activation domain of the androgen receptor, a therapeutic target for castration resistant prostate cancer, is key for its activity as a transcription factor by allowing it to partition into transcriptional condensates. Based on this knowledge we optimized the structure of a small molecule inhibitor, previously identified by phenotypic screening, that targets a specific transactivation unit within the domain that is partially folded and rich in aromatic residues. The optimized compounds had more affinity for their target, inhibited androgen receptor-dependent transcriptional programs, and had antitumorigenic effect in models of castration-resistant prostate cancer in cells and *in vivo*. These results establish a generalizable framework to target small molecules to the activation domains of oncogenic transcription factors and other disease-associated proteins with therapeutic intent.

## Introduction

DNA-binding transcription factors (TFs) are among the most frequently mutated or dysregulated genes in cancer and are among the most coveted targets in oncology^1–3^. For example, TP53, the most frequently mutated gene in cancer, and MYC, the most frequently overexpressed gene in cancer, encode TFs^3^. The rewiring of transcriptional programs is a hallmark of cancer, and oncogenic transcriptional programs of numerous tumor types exhibit exquisite dependence on small subsets of specific TFs^2,4^. Despite their appeal, TFs are considered largely “undruggable” because their protein regions essential for transcriptional activity are intrinsically disordered, rendering them challenging targets for structure-based ligand discovery^5,6^.

Nuclear hormone receptors, e.g. the androgen receptor (AR), are TFs that contain a structured ligand-binding domain (LBD), and anti-androgens targeting the LBD are a common first-line therapy for the treatment of AR-driven prostate cancer^7,8^. However, ∼20% of prostate cancer patients progress into an ultimately lethal stage known as castration-resistant prostate cancer (CRPC) associated with the emergence of constitutively active AR splice variants. Such CRPC-associated AR splice variants lack the LBD, and consist of only the DNA-binding domain and the intrinsically disordered activation domain (AD), rendering them insensitive to LBD-targeting anti-androgens^9–13^. Insights into how intrinsically disordered activation domains function could thus facilitate the development of therapeutic approaches for some of the most lethal cancers.

Recent studies suggest that IDRs in many cellular proteins mediate liquid-liquid phase separation *in vitro*, and partitioning of the proteins into biomolecular condensates in cells^14–16^. Virtually all human TFs, including AR, contain an IDR, and these regions of sequence were recently shown to contribute to the formation of TF condensates and to the partitioning of TFs into heterotypic condensates with transcriptional effectors such as the Mediator co-activator or RNA Polymerase II^17–20^. The molecular basis of TF condensate interactions has only been dissected for a small number of TFs, but in all cases mutations of amino acids in the IDRs that altered phase separation also altered transcriptional activity^17,21–23^. Based on these findings, we hypothesized that the understanding of the molecular basis of phase separation capacity encoded in a TF IDR could be exploited to develop small molecules that alter the activity of oncogenic TFs.

To investigate this hypothesis we chose to study the sequence and structural determinants of AR phase separation. We discovered that this phase transition is required for transactivation and is driven by interactions between aromatic residues, which are scattered in the sequence of the AD but especially frequent in a sub-domain of the AD rich in secondary structure known as transactivation unit 5 (Tau-5). Tau-5, which plays a key role in transactivation by the splice variants associated with CRPC, harbors the binding site of EPI-001^24^, a small molecular inhibitor of the AR AD discovered by phenotypic screening, a derivative of which is under clinical trials for CRPC. Based on our understanding of how this small molecule interacts with the active conformation of Tau-5 we introduced changes in its chemical structure that have led to a substantial improvement of its potency in cells and in a mouse model of CRPC.

## Results

### AR phase separation is driven by tyrosine residues in the activation domain

AR forms mesoscale nuclear “speckles” in hormone stimulated cells, but the biophysical properties of the speckles have been elusive because of their small size, and the hormone-dependent nuclear shuttling of the otherwise cytoplasmic receptor^25–27^. Using live cell and fixed imaging we confirmed that hormone-stimulated endogenous and transgenic AR form clusters (Supplementary Fig. 1). To identify the molecular basis of AR phase separation, we first tested nuclear cluster formation of AR mutants that lack various domains. The full-length AR contains an intrinsically disordered N-terminal activation domain (AD), a central DNA-binding domain (DBD) and a C-terminal LBD (Fig. 1A). In agreement with recent findings^20^, we found that in transiently transfected HEK293T cells, both the full-length AR and the AR-V7 splice variant, that contain the AD and DBD, displayed the capacity to form nuclear clusters, but the DBD alone did not (Fig. 1B). As expected, AR-V7 formed nuclear clusters even in the absence of the hormone (Supplementary Fig. 2A, B). Of note, the AR-V7 splice variant is a key driver of AR-driven CRPC that is resistant to LBD inhibitors^9,11,28^. Cells with higher expression of AR and AR-V7 displayed increased nuclear clustering (Supplementary Fig. 2A, B), consistent with the notion that cluster formation involves phase separation^29^ and is driven by the AR AD.

**Figure 1.**
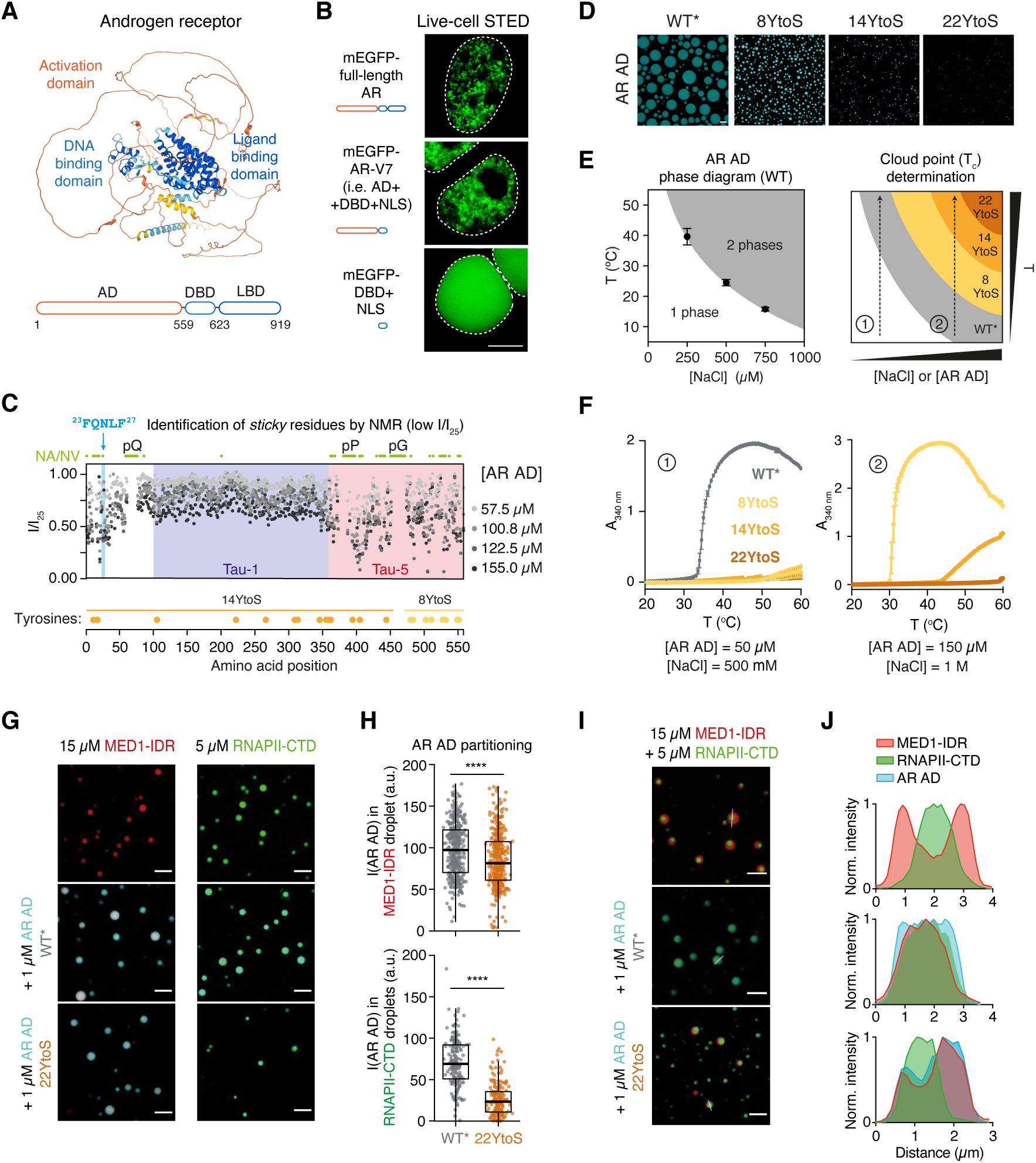
AR phase separation is driven by tyrosine residues in the activation domain. **A**) Structure of AR predicted with AlphaFold. The model is coloured by structure prediction confidence from high confidence (dark-blue) to low confidence (orange-yellow). The known AR domains are highlighted. **B**) Live-cell STED imaging of HEK293T cells transfected with the indicated AR constructs tagged with mEGFP. Cells were imaged after treatment with 10 nM DHT for four hours. Scale bar: 5 μm. Dashed line indicates the nuclear periphery. **C**) Intensity of the NMR resonances of the AR AD as a function of amino acid position, measured for the displayed AR AD concentrations. The position of Transactivation Unit 1 and 5 (Tau-1, Tau-5), and of the ^23^FQNLF^27^ motif are highlighted. Green circles indicate the positions of residues not assigned or not visible (NA/NV) in the NMR spectrum recorded at 25 μM, including residues in regions of low sequence complexity such as poly-glutamine (pQ), poly-proline (pP) and poly-glycine (pG) tracts. Yellow and orange circles represent the positions of tyrosine (Tyr) residues mutated to serine (Ser) in 8YtoS and 14YtoS, respectively; all residues Tyr were mutated to Ser in 22YtoS. **D**) Fluorescence microscopy images of 40 µM AR-AD *in vitro* droplets (WT* and Tyr to Ser mutants) at 1 M NaCl and room temperature. Scale bar: 10 μm. **E**) Schematic representation of the LCST phase diagram of the AR AD (WT) obtained by determining the cloud points of solutions of increasing NaCl concentration (left) and of how cloud point measurements under two different solution conditions (right), labeled as 1 and 2, allow ranking Tyr to Ser mutants in terms of their phase separate capacity. **F**) Determination of the cloud points of AR AD (WT* and Tyr to Ser mutants) under two different solution conditions, labeled as 1 and 2. **G**) Representative merged confocal images of 15 µM MED1-IDR (left column) and 5 µM RNAPII-CTD (right column) droplets obtained at 20 mM NaCl or 50 mM NaCl, respectively, and 10 % ficoll before and after addition of 1 µM AR AD (WT* or 22YtoS). Scale bar: 5 μm. **H**) Quantification of AR AD partitioning into MED1-IDR (top graph) and RNAPII-CTD droplets (bottom graph), by measuring AR AD fluorescence intensity in droplets. Boxes correspond to the mean and the quartiles of all droplets represented as coloured dots from three image replicates. **** p < 0.0001. **I**) Representative merged confocal images of MED1-IDR and RNAPII-CTD multiphasic droplets obtained in 125 mM NaCl and 10% ficoll with and without the addition of 1 µM AR AD (WT* or 22YtoS). Scale bar: 5 μm. j) Normalized intensity plot profile of droplet cross-sections from the images shown in panel **I**.

To identify the residues of the AR AD that drive phase separation, we used solution nuclear magnetic resonance (NMR). This technique provides residue-specific information in the absence of long-range protein order and is thus well suited to study intrinsically disordered proteins^30^. An analysis of the ^1^H-^15^N correlation spectrum of purified AR AD, that provides information on the structural and dynamical properties of the main chain NH groups, revealed that the intensity of the signals of many residues was low, especially at high protein concentration, suggesting that these residues are involved in transient intermolecular interactions^31,32^. We then analyzed the decrease in signal intensity as a function of position in the sequence and residue type, which revealed that the residues involved in such interactions are hydrophobic and, especially, aromatic (Fig. 1C, Supplementary Fig. 2C, D). “Sticky” interacting aromatic residues were particularly enriched around the previously characterized ^23^FQNLF^27^ motif^33^, and, especially, in Tau-5 (Supplementary Fig. 2C).

To directly test the contribution of aromatic residues to AR phase separation we measured how decreasing the aromatic character of the AR AD affects its cloud point (T_c_) *in vitro*^34,35^. We mutated tyrosine residues, the most abundant aromatic amino acid in the AR AD to serines, to generate three mutants: 8YtoS, in which the 8 tyrosines closest to the DBD were mutated, 14YtoS, in which the other 14 tyrosines were mutated, and 22YtoS in which all 22 tyrosines were mutated (Fig. 1C). To increase the stability of purified recombinant AR AD, we introduced an additional mutation (L26P) previously shown to increase protein solubility^36^. The AR AD containing the L26P mutation is referred to as WT* throughout the study. Fluorescently labeled WT* AR AD formed droplets in a concentration-dependent manner (Supplementary Fig. 2E). As expected for phase-separated droplets, the WT* AR AD droplets recovered near 100% of fluorescence after photobleaching with an average half-time of 2.15 seconds (Supplementary Fig. 2F). Mutation of tyrosines to serines led to a reduction in droplet formation (Fig. 1D). Cloud point measurements revealed that phase separation of the AR AD preferentially occurred at high temperature and ionic strength (Fig. 1E) in the so called “Lower Critical Solution Temperature” (LCST) regime, and therefore elevated T_c_ is indicative of a reduction in phase separation capacity. We found that none of the YtoS mutants phase-separated at temperatures lower than 60°C under conditions where T_c_ = 34°C for WT* AR AD (Fig. 1E, F). To better resolve the phase separation capacity of the various YtoS mutants, we increased both protein concentration and ionic strength. We observed that the T_c_ of 8YtoS and 14YtoS were 31°C and 48°C, respectively, while the 22YtoS mutant did not undergo phase separation (Fig. 1F).

Mutation of aromatic residues also compromised the partitioning of the AR AD into heterotypic condensates with known transcriptional effector partners. We incubated AR AD proteins with preassembled droplets formed by purified recombinant MED1 IDR, a frequently used *in vitro* model of Mediator condensates^17,18^, and droplets formed by purified recombinant RNAPII C-terminal domain (CTD), a frequently used *in vitro* model for RNAPII condensates^37^. WT* AR AD partitioned into both MED1 IDR and RNAPII CTD droplets, whereas the partitioning was reduced by the 22YtoS AR AD mutant (Fig. 1G, H). We further modeled heterotypic condensates by mixing MED1 IDR, RNAPII CTD and AR AD proteins. To our surprise, MED1 IDR and RNAPII CTD formed biphasic droplets where the RNAPII CTD was segregated from the MED1 IDR within the MED1 IDR droplets (Fig. 1I). The addition of 1 µM WT* AR AD caused the biphasic droplets to blend into a single phase with the three components homogeneously distributed (Fig. 1I-J). This phenomenon relied on the aromatic character of the AR AD as the addition of 1 µM 22YtoS led to preferential partitioning into the MED1-IDR liquid phase under the same experimental conditions (Fig. 1I, J and Supplementary Fig. 2G). These results collectively reveal that AR phase separation is driven by tyrosine residues within the AR AD.

### AR phase separation propensity is associated with nuclear translocation and transactivation

To test the relevance of phase separation for AR function in cells, we expressed eGFP-tagged wild type full-length AR and mutants that contain the 8YtoS, 14YtoS and 22YtoS substitutions in the AD in AR-negative PC3 cells. Contrary to what is the case for WT ^20^ none of the YtoS mutants formed condensates upon DHT treatment (Fig. 2A). In addition, although these mutations do not alter the NLS, they decreased the nuclear translocation rate of the AR: at t_DHT_ = 60 min, WT AR was nuclear, 8YtoS and 14YtoS were comparably distributed between the cytosol and nucleus and 22YtoS remained primarily cytosolic (Fig. 2A, B). Next we transfected cells with wild type and mutant eGFP-AR-V7 splice variants, which are constitutively nuclear, and measured nuclear cluster formation (Fig. 2C). We observed a decrease of the spatial variance of fluorescence intensity, i.e. granularity, in cells expressing the 8YtoS, 14YtoS and 22YtoS mutants indicating a reduction in the propensity to form clusters (Fig. 2C, D and Supplementary Fig. 3A).

**Figure 2.**
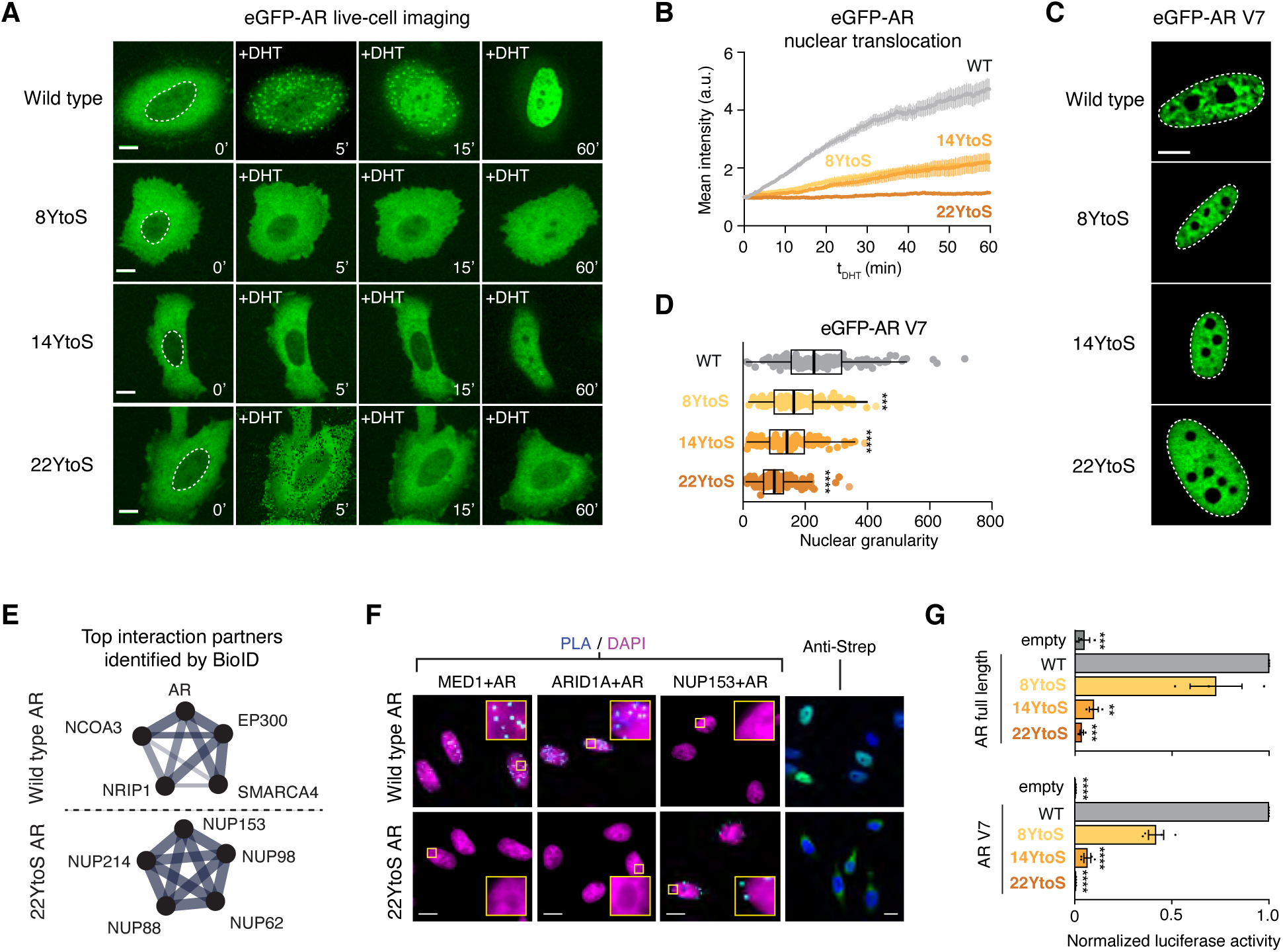
AR phase separation is associated with nuclear translocation and transactivation. **A**) Fluorescence images from live-cell time-lapse movies of PC3 cells expressing eGFP-AR or the indicated mutants. Scale bar: 10 μm. **B**) Quantification of eGFP-AR relative nuclear localization for the indicated cells in panel A as a function of time elapsed since the addition of 1 nM DHT (t_DHT_). Error bars represent s.d. of n ≥ 15 cells per time point. **C**) Representative images of live PC3 nuclei expressing eGFP-AR-V7 WT and Tyr to Ser mutants. **D**) Quantification of the nuclear granularity for the indicated cells in panel C, where each dot represents one nucleus and p values are from a Dunnett’s multiple comparison test against the WT (n ≥ 150 cells per condition). **E**) Selected gene ontology (GO) Molecular Function networks enriched in the Top 75 most abundant hits (BFDR ≤ 0.02, FC ≥ 3) for the indicated bait. *Androgen receptor binding* (WT) and *Structural constituent of the nuclear pore* (22YtoS) protein–protein interaction (PPI) networks are shown, line thickness corresponds to strength of published data supporting interactions, generated from STRING (string-db.org). Additional GO results are provided in Supplementary Fig. 3E and Supplementary Table 1. **F**) Proximity ligation assays (PLA) using the indicated antibodies are shown in cyan with DAPI staining in magenta in DHT treated PC3 cells. Streptavidin labeling is shown in green with DAPI in blue (far right panels) in DHT treated PC3 cells, scale bars 10 μm. **G**) Transcriptional activity of AR and Tyr to Ser mutants assessed with a luciferase reporter assay for AR (t_DHT_= 1 h, top) and AR-V7 (bottom) in HEK293 cells. Empty stands for empty vector and p values are from a Dunnett’s multiple comparison test against the WT (n = 3 in upper panel, n = 4 in lower panel).

To further probe the mechanistic basis of reduced translocation of phase separation-deficient AR mutants, we mapped the interactomes of WT and 22YtoS full length AR using BioID-mass spectrometry. The WT AR and 22YtoS proteins were fused to a FLAG tagged Mini-TurboID (MTID) enzyme and were introduced into PC3 cells using a lentiviral vector. Addition of biotin for 1 h led to increased protein labeling, demonstrating that the MTID enzyme was functional (Supplementary Fig. 3B). We carried out BioID-MS collecting samples before and after DHT treatment (t_DHT_ = 60 min): SAINTq analysis revealed that a large number of proteins were enriched following stimulation by DHT in cells expressing WT AR (Supplementary Fig. 3C, 3D). Enrichment analysis (STRING) identified categories primarily related to transcription, including a number of established AR interactors (Fig. 2E, Supplementary Fig. 3E, Supplementary Table 1). By contrast, the 22YtoS mutant identified fewer proteins, with little overlap to WT AR (Supplementary Figs. 3E, F): enrichment analysis of 22YtoS identified several categories related to nuclear transport with 5 nucleoporins identified amongst the top 75 most enriched proteins (Fig. 2E, Supplementary Fig. 3E, Supplementary Table S1). To validate these observations, we performed proximity ligation assays (PLA) for several of the top hits in WT and mutant AR, including the SWI/SNF component ARID1A, the Mediator component MED1, and NUP153. PLA signal was clearly evident for WT AR with MED1 and ARID1A, whereas no interaction was observed for 22YtoS (Fig. 2F); by contrast, 22YtoS showed clear interaction with NUP153 in the perinuclear space, that was absent for WT AR (Fig. 2F).

Finally, we measured the transcriptional activity of both AR and AR-V7 in transiently transfected cells in which we co-transfected a luciferase reporter gene driven by an AR-dependent promoter. We found that mutations of tyrosines in the activation domain of the AR led to a reduction of the transcriptional activity of both the full-length AR, and the AR-V7 splice variant: the degree of reduction in transcriptional activity depended on the number of tyrosine residues mutated (Fig. 2G). Taken together these results indicate that AR aromatic mutants that have reduced phase separation have lower nuclear translocation rate, increased association with nuclear pore, and reduced transcriptional activity.

### Short unstable helices enhance AR phase separation

Transcriptional activation involves interactions between short sequence motifs in activation domains (ADs) - also known as activation units - and members of the transcriptional machinery^38–42^. Some motifs are known to fold into ɑ-helices when interacting^43–46^. Therefore, we tested whether regions with ɑ-helical propensity in the AR AD contribute to its phase separation behavior. We annotated seven regions with helical propensity in the AR AD by NMR, which confirmed the helical propensity of the flanking region of the polyglutamine (pQ) tract starting at position 58, and of the ^179^LKDIL^183^ motif in the Tau-1 region (Fig. 3A), consistent with previous studies^47,48^. To map regions with helical propensity in the Tau-5 region, that have low peak intensity in the spectrum of full-length AR AD, we performed NMR on a Tau-5 fragment (referred to as Tau-5*), which revealed high helical propensity of the ^39^7WAAAAAQ^403^ motif (Fig. 3A). Previous X-ray crystallography work has shown that the ^23^FQNLF^27^motif forms an ɑ-helix when interacting with the AR LBD^33^. Our previous NMR experiments have also shown that the ^433^WHTLF^437^ motif in Tau-5 forms a helix when interacting with TFIIF, and identified two additional motifs ^232^DNAKELCKA^240^ and ^351^LDEAAAYQS^359^ with weak helical propensity^49^ (Fig. 3A).

**Figure 3.**
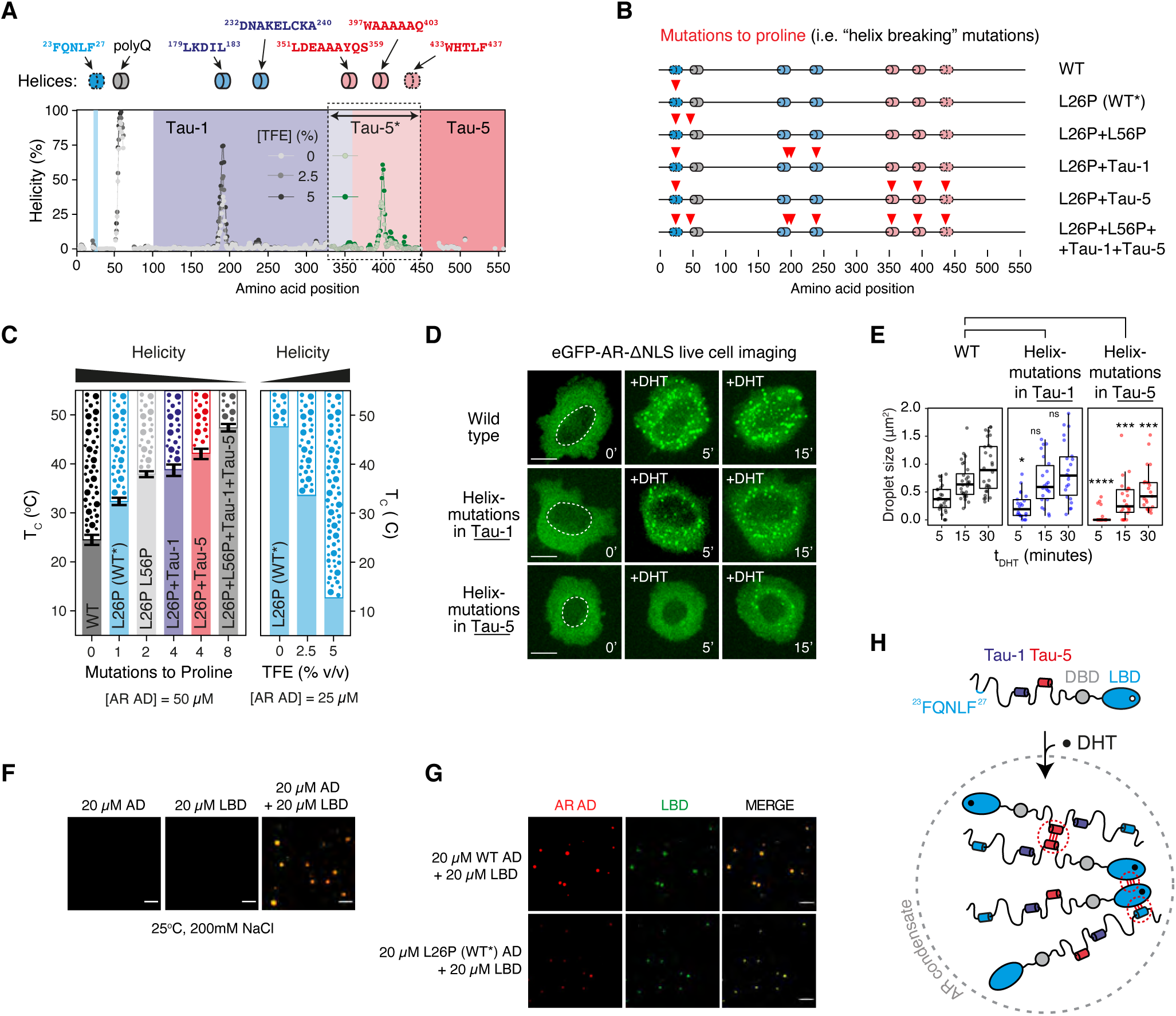
Short unstable helices enhance AR phase separation. **A**) Annotation of short helical motifs in the AR AD. The plots shows helical propensity of the WT* AD measured by NMR in the absence or presence of 2.5 or 5% TFE. The Tau-1 and Tau-5 regions are highlighted. Discontinuous contour denotes motifs that fold when bound to globular binding partners. Helicity values were derived from the H, N, C’ and Cɑ main chain chemical shifts measured by using solution NMR and using the δ2D software^71^. Values in green correspond to an equivalent experiment carried out with construct Tau-5* ^24^, which was used because the most informative resonances are invisible in AR AD due to their involvement in transient long-range interactions. **B**) Mutants generated to investigate the effect of reduced helical propensity on phase separation. Color code is the same as used in panel A. **C**) Cloud point measurements of AR AD purified proteins containing the indicated proline mutations, or in the presence of TFE. **D)** Representative live cell fluorescence microscopy images of DHT-treated PC-3 cells expressing the indicated eGFP-AR-ΔNLS mutants. Scale bars 10 μm. **E**) Quantification of droplet size formed by eGFP-AR-ΔNLS mutants in OC-3 cells as a function of t_DHT_. Each dot corresponds to the mean droplet size in a single cell (n > 20 cells). P values are from a Mann-Whitney’s U test. **F**) Fluorescence microscopy images of purified AR AD, LBD and an equimolar mixture of the two proteins *in vitro*. Images correspond to the merge of red (AR AD) and green (LBD) channels. Images obtained at 200 mM NaCl and 20 μM protein where ca 1% protein is labeled. **G**) Fluorescence microscopy images of an equimolar mixture of purified hormone-bound LBD with AR AD WT (top) or AR AD WT* (bottom) *in vitro*. Samples were prepared at 200 mM NaCl and 20 µM protein where ca 1% protein is labeled. Images correspond from left to right to AR AD (red), LBD (green channel) and the merge of them. **H**) Schematic model illustrating the key interactions that drive AR phase separation.

To investigate the contribution of helical propensity to AR phase separation, we introduced helix-breaker proline substitutions in the AR AD within or immediately adjacent to the annotated helices (Fig. 3B) and measured the cloud point (T_c_) of purified recombinant AR AD proteins (Fig. 3C). We found that the L26P mutation (WT*), which prevents helix formation by ^23^FQNLF^27^, increased the T_c_ by 8°C (Fig. 3C). Next, we studied three mutants, in the L26P background, designed to decrease helicity of the polyQ tract (L56P), Tau-1 (A186P, L192P and C238P) and Tau-5 (A356P, A398P and T435P). We observed that these mutations increased T_c_ to different extents: L56P increased it by 5°C, as did mutations in Tau-1, but mutations in Tau-5 had a larger effect, of *ca* 10 °C. (Fig. 3B, C). We also analyzed the effect of TFE, a co-solvent that induces helicity in disordered peptides and proteins^50^, on phase separation propensity. We found that it indeed increased the helical propensity of the most helical motifs in Tau-1 and Tau-5 (Fig. 3A) and strongly decreased the T_c_ of the AD: by 12°C at 2.5% TFE (v/v) and by 35°C at 5% (Fig. 3C). These results suggest that regions with helical propensity enhance AR AD phase separation *in vitro*.

We next investigated the effect of reduced helicity on AR phase separation in cells. For this, we developed an assay to characterize AR condensates in the absence of DNA binding by introducing a mutation in the NLS (eGFP-AR-ΔNLS) that prevents AR nuclear translocation (Supplementary Fig. 4A). Stimulation of PC3 prostate cancer cells expressing eGFP-AR-ΔNLS with DHT leads to the formation of large cytosolic AR condensates and facilitates quantifying their amounts and dimensions (Supplementary Fig. 4B, C). Live cell imaging revealed that the cytoplasmic AR condensates had spherical shape, and their number hardly changed with time but their size increased substantially (Supplementary Fig. 4D). In addition, the condensates were observed to undergo fusion events (Supplementary Fig. 4E), and recovered fluorescence intensity quickly after photobleaching (mobile fraction = 94 ± 8%, t_1/2_= 2.29 ± 1.17 s) both after 1h and 24h from DHT-stimulation (Supplementary Fig. 4F), these features are characteristics of phase separation^29^. Helix-breaking mutations in Tau-1 had a negligible effect on the formation and dynamics of cytosolic AR condensates, but mutations in Tau-5 significantly decreased both the number and size of condensates following short term (5-15 min) hormone exposure (Fig. 3B, D, E, Supplementary Fig. 4G). These results indicate that regions with helical propensity in the Tau-5 sub-domain enhance AR phase separation in cells.

The results described above suggest that aromatic residues and short unstable helices, particularly in the Tau-5 region, play important roles in AR phase separation, but do not explain why hormone binding is necessary to trigger AR phase separation. We hypothesized that the interaction between the AR LBD and AD enhances the phase separation capacity of the latter. To test this idea, we incubated the AD *in vitro,* at a concentration (20 µM) and solution conditions (25°C, 200 mM NaCl) that do not lead to phase separation, with 1 molar equivalent hormone-bound LBD. We observed the formation of heterotypic droplets containing both domains (Fig. 3F). WT* AR AD, containing the helix-breaking L26P mutation, formed smaller droplets in the presence of the LBD (Fig. 3G, Supplementary Fig. 4H). Consistent with the *in vitro* data, eGFP-AR-ΔNLS that lacks the ^23^FQNLF^27^ motif formed fewer and smaller condensates in PC3 cells (Supplementary Fig. 4I-K), in agreement with recent reports^20^. These results collectively suggest that at least three interactions play key roles in phase separation of the hormone-exposed androgen receptor: i) interaction between the LBD and AD that requires helix formation by the ^23^FQNLF^27^ motif, ii) aromatic residues especially around the short helical regions in the Tau-5 sub-domain of the AD, and iii) the previously described dimerization interface in the LBD^51^ (Fig. 3H).

### Rational design of small molecules with enhanced potency

EPI-001 is a small molecule inhibitor of the AR AD, identified by phenotypic screening, that interacts selectively with aromatic residues in secondary structure elements in Tau-5^24,52^. We therefore hypothesized that optimizing the distance and orientation of aromatic rings in EPI-001 and modulating the flexibility of the functional group connecting them, could increase the compound’s potency. We synthesized a series of compounds where the carbon atom between the aromatic rings of EPI-002, the (2R,19S) stereoisomer of EPI-001, was replaced by two carbon atoms separated by a single (compound 4aa), a double (2aa, *cis* and 3aa, *trans*) or a triple (1aa) bond (Fig. 4A, B). The potency of the compounds was tested in LNCaP cells transfected with a luciferase reporter driven by an AR–dependent promoter and enhancer^53–56^. Compounds 2aa and, especially, 1aa were the most potent inhibitors, substantially more potent than EPI-002 (Fig. 4A, Supplementary Fig. 5A).

**Figure 4.**
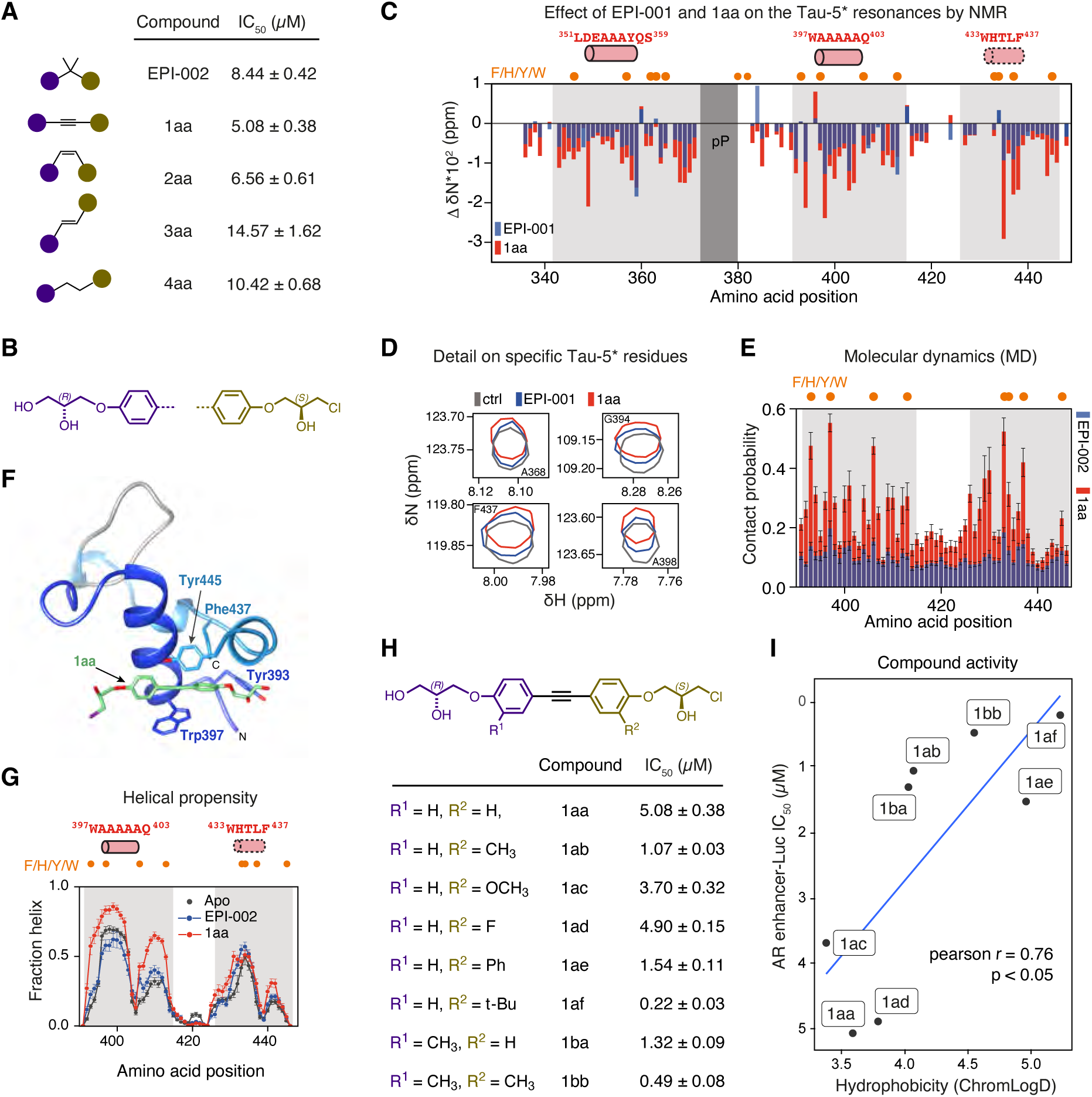
Optimization of the structure of EPI-001. **A,B**) Chemical structures of EPI-002 and compounds with modified linker between the two aromatic rings of EPI-002. (**A**) Schematic of the structures and the corresponding IC_50_ measured in androgen-induced PSA-luciferase assay. Purple and brown circles correspond to chemical groups depicted in (**B**), where R_1_ and R_2_ are hydrogens. **C**) ^15^N chemical shift changes in the NMR spectra of Tau-5* (60 μM) as a function of amino acid positions caused by addition of 1 molar equivalent of EPI-001 (blue) and 1aa (red). Orange circles indicate aromatic amino acids positions in the sequence of Tau-5*. R1-3^24^ and polyP regions are highlighted in light and dark grays, respectively. Samples contained 200 mM NaCl and 2 % DMSO-d_6_. **D**) Selected regions of Tau-5* ^1^H,^15^N BEST-TROSY spectra in the absence (gray) and presence of 1 mol equivalent of EPI-001 (blue) and 1aa (red), with an indication of partially folded helices. **E**) Per-residue contact probabilities observed in REST2 MD simulations between Tau-5 residues 391-446 and the compounds: EPI-002 (blue) and 1aa (red). Contacts are defined as occurring in frames where any non-hydrogen ligand atom is within 6.0 Å of a non-hydrogen protein atom. Orange circles on top of the panel represent the positions of aromatic residues. **F**) Illustrated MD Snapshot of the AR AD interacting with 1aa: helices are colored in dark and light blues colors, while the loop between them is gray. 1aa is shown in green color and the chlorine is colored in purple. **G**) Helical propensities of Tau5_R2_R3_ in its apo form (black) and in bound conformations obtained from simulations run in the presence of EPI-002 (blue) and 1aa (red) with an indication of the positions of helical motifs and aromatic residues in the sequence. The data was obtained from the 300 K REST2 MD simulations. **H**) Compounds developed from 1aa and their corresponding potency in the androgen-induced PSA-luciferase assay. **I**) Correlation between the activity of the compounds and their in the PSA-luciferase assay and their hydrophobicity.

To test whether the change in compound structure led to an optimized interaction with the AR AD, we analyzed the NMR spectrum of the Tau-5* protein in the presence of the compounds. The chemical shift perturbations caused by 1 molar equivalent of compound 1aa were larger than those induced by EPI-001, indicating enhanced interaction for 1aa (Fig. 4C, D). We also simulated the interaction of the compounds with residues 391 to 446 of the AR AD ^57^ and observed that its atoms contacted those of 1aa more frequently than those of EPI-002 (Fig. 4E-G, Supplementary Fig. 5B), leading to a more stable and structured complex (simulated K_D_ 1.4 ± 0.1 mM for 1aa vs K_D_ 5.2 ± 0.4 mM for EPI-002), in agreement with the NMR and gene reporter data. Given that the liquid droplets formed by the AR AD are stabilized by hydrophobic and aromatic interactions (Fig. 1) we also synthesized analogues of 1aa with substitutions in positions R_1_ and R_2_ (Fig. 4H, I, Supplementary Fig. 5C) aimed at modulating the hydrophobic and aromatic character of the compounds: introduction of a methyl (CH_3_) group at R_1_ (1ab) or R_2_ (1ba) increased potency from IC_50_ *∼*5 µM to IC_50_ *∼*1 µM whereas introduction of this group in both positions (1bb) further increased it to 0.5 µM in the luciferase reporter system. In line with these findings, the introduction of a *tert*-butyl (C(CH_3_)_3_) group at R_2_, bearing three methyl groups (1af), afforded IC_50_ to 0.22 µM. Substitution of hydrogen by fluorine (1ad) or a methoxy (CH_3_O) group (1ac) at position R_2_ hardly changed potency but introduction of an additional aromatic ring (1ae) instead increased potency to ∼1.5 µM in the reporter assay (Fig. 4H, I).

Next, we measured the potency of the compounds as inhibitors of AR-V7 by using the V7BS3-luciferase reporter that is specific for AR-V7^58^. As expected, 5 µM enzalutamide, that binds to the AR LBD, had no activity against AR-V7-induced V7BS3-luciferase activity, whereas 35 µM EPI-002 blocked luciferase activity, consistent with previous reports^56^ (Supplementary Fig. 5D). Importantly, 1ae was the most potent inhibitor of AR-V7 transcriptional activity, in a dose-response manner (Supplementary Fig. 5E), whereas 1ab and 1bb had no inhibitory effects (Supplementary Fig. 5F, G). The effect of 1ae on the transcriptional activity of AR-V7 was not due to non-selective inhibition of transcription or translation as determined by its negligible effect on the activity of the non-AR-driven reporter, AP-1 luciferase activity (Supplementary Fig. 5H). In line with these results, 1ae blocked the proliferation of both LNCaP and LNCaP95 cells, driven by full-length AR and AR-V7, respectively (Supplementary Fig. 5I), while enzalutamide blocked the proliferation of LNCaP cells only, consistent with its mechanism of action (Supplementary Fig. 5J).

To better understand the mechanisms by which small molecule inhibitors of the AR AD decrease transcriptional activity and thus prevent the proliferation of prostate cancer cells we performed a BioID-MS analysis in LNCaP cells stably expressing MTID-AR-WT treated with EPI-001 or 1ae (Fig. 5A and Supplementary Fig. 6A) and found that both inhibitors caused a general decrease in AR interactions (Fig. 5B, C). 1ae caused a stronger decrease in total interactions than EPI-001 and in addition showed a significant decrease in known AR interactors according to the BioGrid database (Fig. 5D). An enrichment analysis of the depleted AR-interactors revealed that the decrease was more marked for 1ae in all categories, in line with its higher potency (Fig. 5E, Table S2). Focusing on Mediator we found that, amongst all sub-units, MED1 was the most significantly reduced by 1ae (Fig. 5F, G and Supplementary Fig. 6B, C). Finally, we used PLA between AR and MED1 or ARID1A to validate these findings and found a visible reduction in the total number of foci per cell, with an increasing time of drug incubation with both inhibitors (Fig. 5H). Again, 1ae proved to be more potent than EPI-001 in reducing the strength of these interactions (Fig. 5I). Put together, these data show that the small molecule inhibitors decrease the extent to which AR interacts with the transcription machinery, thus leading to an inhibition of transcriptional activity.

**Figure 5.**
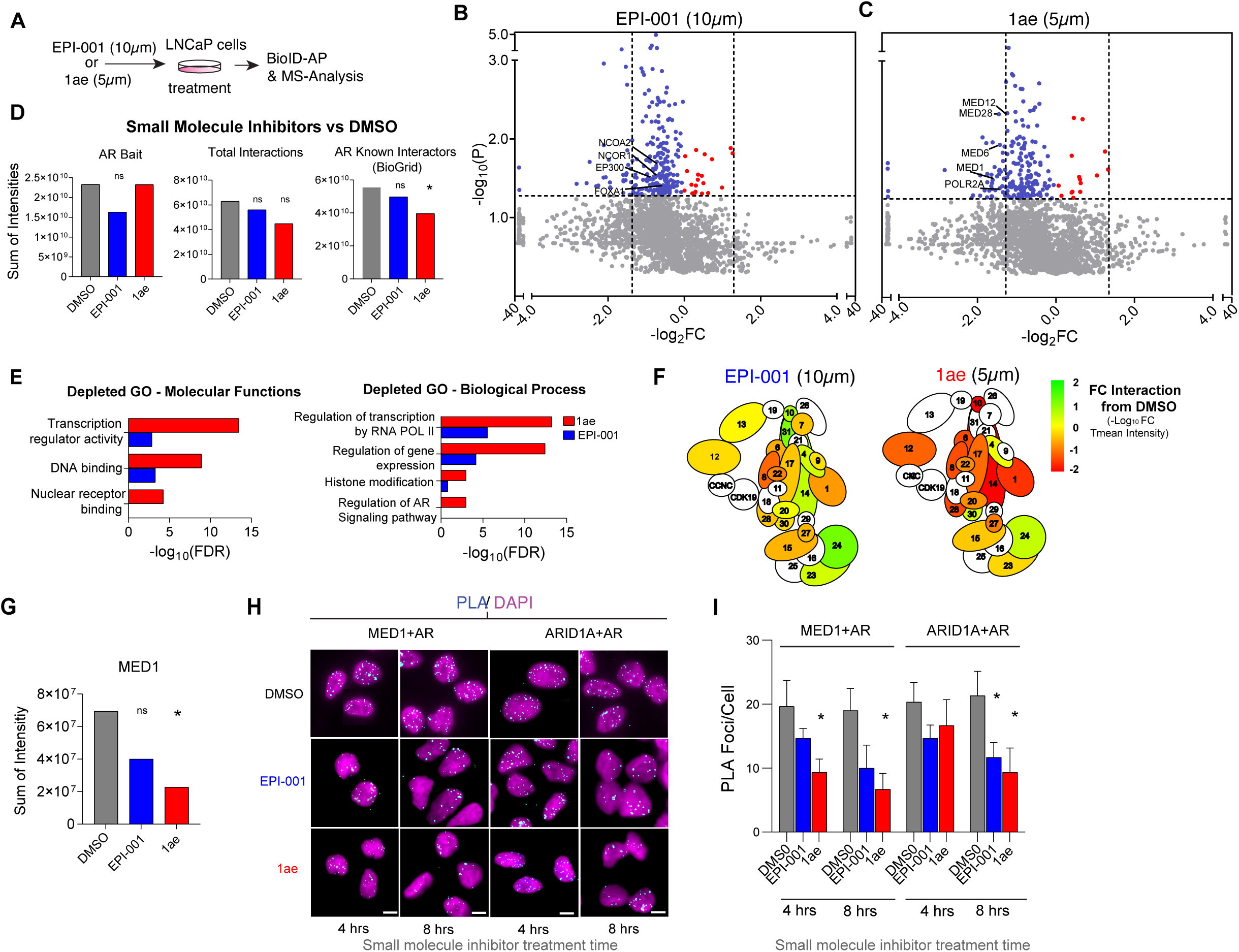
1ae decreases interactions between AR and the transcription machinery. **A)** Schematic of method for small molecule treatment of cells and BioID experiment. **B)** BioID-MS of LNCaP MTID-AR-WT cells treated with 1 hour EPI-001 10uM and then 2 hrs of DHT and Biotin. SAINTq intensity data of LogFC of intensity of interaction vs DMS0 treated. Red highlights reduced interactors vs DMSO, pvalue < 0.05 and dotted line Log_2_FC < -1.5. Blue highlights increase in interaction vs DMSO pvalue < 0.05 and dotted line Log_2_FC > 1.5. Annotated with proteins of interest. N=3. **C)** As in B, BioID-MS of LNCaP MTID-AR-WT cells but treated with 1 hour 1ae 5uM and then 2 hrs of DHT and Biotin. N=3. **D)** TMean SAINTq intensity of individual proteins (bait), total interactors (all) or collated AR known interactors sourced from BioGrid (https://thebiogrid.org/) were compared across LNCaP MTID-AR-WT with DMSO or treated cells with small molecule inhibitors. Statistical significance was determined by Mann-Whitney U test *P < 0.05. **E)** GO search terms of key Biological process and Molecular functions of SAINTq intensity data from B and C (BFDR < 0.02, Depleted =Log_2_FC < -1.5) of the LNCaP MTID-AR-WT treated cells with small molecule inhibitors vs DMSO. Full categories see Table S2. **F)** BioID-MS of LNCaP MTID-AR-WT interaction with Mediator complex with small molecule inhibitors vs DMSO. SAINTq data, colour indicates strength of interaction change from LogFC_10_ Tmean of intensity. **G)** TMean SAINTq intensity of individual proteins MED1 subunit was compared across LNCaP MTID-AR-WT with DMSO or treated cells with small molecule inhibitors. Statistical significance was determined by Student’s t-test against the control arm. *P < 0.05. **H)** Proximity ligation assay (PLA) using the indicated antibodies shown in cyan with DAPI staining in magenta in small molecules or DMSO and DHT treated LNCaP MTID-AR-WT cells at times indicated. Scale bars=10 μm**. I**) Quantitation of PLA foci shown in I per cell scored. TTest used * p value < 0.05

### Compound 1ae is a potent inhibitor of AR-dependent transcription and tumor growth

The potency and specificity of 1ae and EPI-001 on the AR-driven oncogenic transcriptional program was investigated further by RNA-Seq after treating LNCaP prostate adenocarcinoma cells with ∼IC_10_ and ∼IC_50_ doses of the compounds for 6 and 24 hours (Fig. 6A-C and Supplementary Fig. 7A, B). As expected, 6 hour treatment with IC_10_ concentrations had negligible effect on the gene expression profile of prostate cancer cells (Supplementary Fig. 7A-D). In contrast, 24 hours treatment with 25 µM EPI-001 led to the differential expression of 64 genes, and 24 hour treatment with 5 µM 1ae led to the differential expression of 231 genes, compared to DMSO-treated control cells (Fig. 6D). Gene set enrichment analysis (GSEA) revealed that downregulated genes were significantly enriched for known AR-targets, for both EPI-001 and 1ae (p_adj_< 0.01) (Supplementary Fig. 7C-E). Both EPI-001 and 1ae dysregulated the same subset (5/50) of pathways tested with GSEA (Fig. 6E and Supplementary Fig. 7E). The significantly dysregulated pathways included the AR response pathway and other pathways known to be hyperactive in CRPC ^59,60^. Of note, 5 µM 1ae treatment led to a more profound reduction in the expression of all downregulated and differentially expressed genes induced than 25 µM EPI-001 treatment (Fig. 6F and Supplementary Fig. 7D) and in addition decreased the levels of soluble AR as revealed by Western blot analysis (Fig. 6G and Supplementary Fig. 7F). These results indicate that 1ae inhibits AR-dependent targets in prostate cancer cells and is more potent in its transcriptional inhibitory effect than EPI-001.

**Figure 6.**
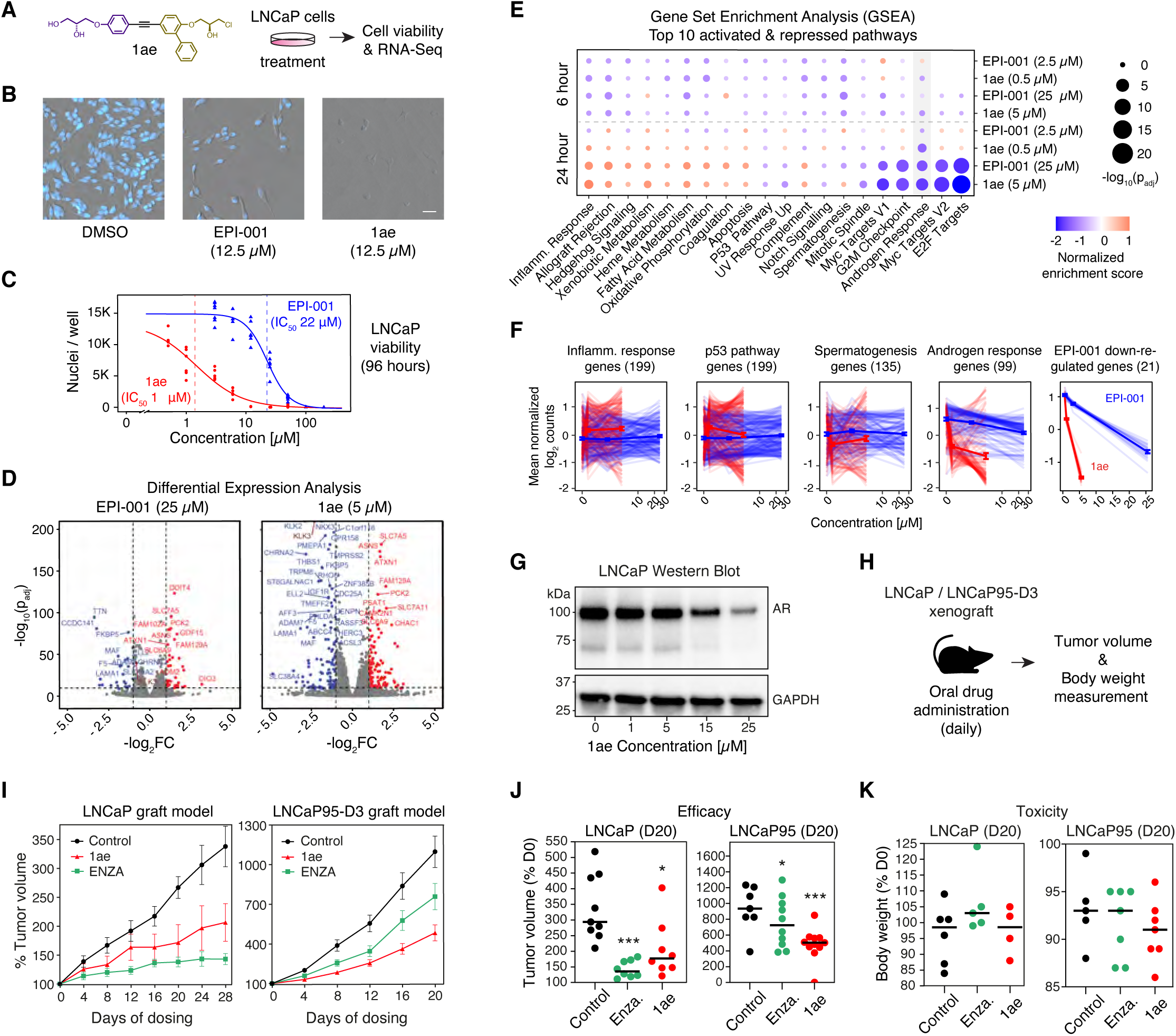
Compound 1ae is a potent inhibitor of AR-dependent transcription and tumor growth. **A**) Molecular structure of compound 1ae and schematic to investigate its effect on AR-dependent prostate carcinoma (LNCaP) cells. **B**) Representative images of LNCaP nuclei (counterstained with Hoescht) in indicated condition after 96 hours of treatment (scale bar : 50 μm). **C**) Dose response curve (log-logistic fit) of viable LNCaP nuclei treated with indicated compound as a function of compound concentration, with IC_50_s for EPI-001 or 1ae calculated from dose response curve (n = 6). **D**) Volcano plots of differentially expressed genes in LNCaP cells treated with EPI-001 and 1ae for 24 hours at the concentration near IC_50_ versus DMSO at 24 hours (Fold change cut offs: 2x, 0.5x). **E**) Gene set enrichment analysis of top 10 enriched and top 10 depleted msigdb hallmark signature pathways^72^ in LNCaP cells treated with EPI-001 or 1ae, at indicated time points and concentrations versus DMSO at the same time point. Circles scaled in size to significance value of normalized enrichment score (log of adjusted p-value) and color gradient scaled to normalized enrichment score of the indicated pathway analyzed with GSEA. Hallmark Androgen Response Pathway shaded in gray (N = 3). **F**) Line plots of log transformation of mean normalized counts of the indicated gene sets in LNCaP cells treated with EPI-001 or 1ae, as a function of compound concentration. Light lines represent individual genes, dark lines represent average of all genes, bars represent standard error (N = 3). **G**) Representative western blot of AR in LNCaP cells pretreated with 1ae at indicated concentration for 1 hour prior to activation with 1 nM DHT for 4 h: GAPDH was used as lysate loading control (bottom). **H**) Schematic of LNCaP and LNCaP95-D3 xenografts CRPC model. **I**) Tumor volume of mice LNCaP (left) and LNCaP95-D3 (right) xenografts during the course of the *in vivo* experiment. Values are presented as percentage relative to the tumor volume measured at the first day of drug dosing. **J**) Average tumor volume on day 20 of the xenograft experiments. Values are presented as percentage relative to the tumor volume measured at the first day of drug dosing. Statistical significance was determined by Student’s t-test against the control arm. *P < 0.05, ***P < 0.001. Error bars represent the SEM of n > 8 (LNCaP) or n > 7 (LNCaP95-D3) tumors per treatment group. **K**) Body weight of animals on day 20 of the xenograft experiments. Values are presented as percentages relative to body weight measured on the first day of drug dosing.

The *in vivo* efficacy of 1ae was tested on human CRPC xenografts in castrated murine hosts. For this purpose, LNCaP cells (driven by the full-length AR), and LNCaP95-D3 cells (expressing elevated levels of the AR-V7 splice variant)^61^ were xenografted into murine hosts that were castrated. 1ae was administered at a daily dose of 30 mg/kg body weight for 28 days (Fig. 6H). After 20-28 days of treatment, 1ae significantly reduced tumor volumes both in the LNCaP and LNCaP95-D3 xenograft model compared to the control animals (Fig. 6I, J). In the AR-V7 driven LNCaP95-D3 xenograft model of CRPC, 1ae outperformed enzalutamide, a second-generation antiandrogen that targets the AR LBD. No overt toxicity was observed for 1ae as determined by no substantial differences in body weight of the animals at the end of the experiment (Fig. 6K).These results suggest that 1ae has *in vivo* antitumor activity in CRPC xenograft models, and outperforms enzalutamide.

## Discussion

Our data provides insights into the molecular basis of phase separation encoded in the AR, that may be general for other nuclear hormone receptors and transcriptional regulators. AR condensates are stabilized by interactions between aromatic residues, similar to condensates formed by various prion-like proteins^32,62,63^. In the AR activation domain, these residues cluster in the ^23^FQNLF^27^ motif and, especially, within the Tau-5 region located at the C-terminus of the domain (Fig. 1C). An inter-domain interaction between the ^23^FQNLF^27^ motif and the LBD, stimulated by hormone binding to the LBD, also contributes to stabilizing the condensates because it provides additional valencies for phase separation^33,51,64^. The presence of partially folded helices in the AR AD further facilitates phase separation, suggesting that disordered proteins might gain structure in condensates^65,66^. Such structures may in turn increase the “druggability” of the target protein, a proposal consistent with the evidence for structure-activity relationship (SAR) of the AR AD-targeting compounds described here (Fig. 4).

The results presented here reveal unexpected insights into the link between phase separation and the molecular functions of transcriptional regulators. We found that reducing phase separation of the AR AD by reducing its aromatic character inhibited transcriptional activity, consistent with previous mutagenesis studies on a small number of TFs^17,21–23^. In addition, mutations that reduced AR phase separation also inhibited nuclear translocation of the receptor (Fig. 2A, B). Aromatic AR mutants, which still contain the intact nuclear localization signal (NLS), preferentially interacted with nucleoporins on the nuclear surface (Fig. 2E, F). We speculate that aromatic residues in the AR AD interact directly with aromatic residues in FG repeats of nucleoporins, without the mediation of nuclear import receptors and adaptor proteins^67^. This idea is supported by the observation that substituting surface residues by aromatic ones in a large globular protein increases the rate of nuclear translocation^68^, and suggests a potentially generic nuclear import mechanism, different from linear sequence information encoded in nuclear localization signals.

Finally, we provide evidence that targeting small molecules to the region of the AR that drives its phase separation has antitumorigenic effect in an *in vivo* CRPC model driven by an “undruggable” AR variant. Anti-androgens used as first line therapy against prostate cancer, such as enzalutamide, target the LBD, and inhibit activation by androgens^69^. A hallmark of CRPC is the emergence of AR splice variants that lack the LBD and are thus resistant to this class of drugs. Such isoforms consist of the DNA binding domain and the disordered activation domain of the receptor, suggesting that inhibition of the AR AD could inhibit prostate cell proliferation in CRPC. We took advantage of a previously described small molecule, EPI-001, derivatives of which are under clinical investigation^70^, clarified its mode of action, and substantially improved its potency by using insights into the molecular basis of AR phase separation.

In summary we establish the basis to further develop anti-CRPC drugs and in addition propose a generalizable framework to target with therapeutic intent the phase-separation capacity of intrinsically disordered regions in oncogenic transcription factors and other disease-associated proteins^6^.

## Acknowledgements

We thank Tanja Mittag (St Jude Children’s Research Hospital), Cristine Helsen and Frank Claessens (University of Leuven), Diego Presman and Gordon Hager (NCI-NIH), and the members of the Bulut-Karslioglu, Hnisz and Salvatella laboratories for helpful discussions. We thank Alf Honingmann (MPI-CBG) for help with image analysis, Rene Buschow (MPI-MG) for introduction to STED microscopy and Helene Kretzmer (MPI-MG) for advice on bioinformatics analyses. We acknowledge help and support from the advanced digital microscopy, high throughput protein expression and mass spectrometry, biostatistics/bioinformatics, and sequencing core facilities at MPI-MG and IRB. Finally we acknowledge help and support from the spanish ICTS Red de Laboratorios de RMN de biomoléculas (R-LRB). T.H.S. was supported by the Intramural Research Program of the National Institutes of Health. P.R. and J.Z. were supported by the National Institutes of Health under award R35GM142750. J.Z. was also supported by the China Scholarship Council. M.D.S acknowledges funding from the National Cancer Institute of the National Institutes of Health (R01CA105304). X.S. was supported by AGAUR (2017 SGR 324), MINECO (BIO2015-70092-R and PID2019-110198RB-I00), the Fundació La Caixa (CI20-00098), the AECC (INNO20010FRIG) and the European Research Council (CONCERT, contract number 648201). IRB Barcelona is the recipient of a Severo Ochoa Award of Excellence from MINECO (Government of Spain).

## Author contributions

Conceptualization: S.Ba., P.M-C, M.P., M.F-V., M.D.S., A.Ri.,D.H. and X.S; Methodology: S.Ba., P.M-C., M.P., M.F-V., E.S., S.Bi., J.Z., L.B., A.R. and D.M.; Investigation and formal analysis: S.Ba., P.M-C., M.P., M.F-V., E.S., M.L., C.A.B., C.S-Z., S.Bi., J.Z., C.G-C., C.B. and M.B.; Software: S.Ba., M.L, and C.A.B.; Supervision: B.M., A.E., X.V., N.R.W., J.W., T.T., I.B-H., S.V., J.G., P.R., T.H.S., M.D.S., A.Ri., D.H. and X.S; Writing - Original Draft: S.Ba., P.M-C., M.P., D.H. and X.S: Writing - Review and Editing: all authors. Funding acquisition: P.R., T.H.S., M.D.S., A.Ri., D.H. and X.S

## Competing interests

M.F.-V. is an employee of Dewpoint Therapeutics. M.P. is an employee of Nuage Therapeutics. D.H., M.B. and X.S. are founders of Nuage Therapeutics. D.H. and X.S are scientific advisors of Nuage Therapeutics.

## Methods

### Resource availability

#### Materials Availability

All unique reagents generated in this study are available from the Lead Contacts with a completed Materials Transfer Agreement.

#### Data & Code Availability

All custom code is available upon request.

RNA-sequencing data has been deposited in the NCBI GEO database with accession GSE206853. Reviewers can access data at:

URL: https://www.ncbi.nlm.nih.gov/geo/query/acc.cgi?acc=GSE206853

Token: czkfcwuejzetpev

### Experimental Model & Subject details

#### Cell culture

PC3 (ATCC; CRL-1435) and LNCaP clone FGC (ATCC; CRL-1740) cells were cultured in RPMI 1640 containing 4.5 g/l glucose (Glutamax, Gibco) supplemented with either 10% (v/v) charcoal stripped serum (CSS, Thermo A3382101) or 5% FBS (v/v), as specified in method details, and antibiotics. Induction of transcriptional activation by the androgen receptor in experiments using 5% FBS cultured LNCaP cells (Fig. 6A-F and Supplementary Fig. 1A,D and 7A-E) was verified using high resolution microscopy and qRT-PCR. HEK293T cells (ATCC; CRL-3216) and AR-eGFP Hela stable cells (gift from Pennuto lab) were maintained in DMEM containing 4.5 g/l glucose supplemented with 10% (v/v) charcoal stripped FBS and antibiotics. LNCaP95 was obtained from Dr. Stephen R. Plymate (University of Washington, Seattle, Washington, USA) and cultured in phenol red-free RPMI supplemented with 10%(v/v) charcoal stripped FBS (Gibco) and antibiotics. Cells were cultured in a humidified atmosphere containing 5% CO_2_ at 37 °C. Cells were negative for mycoplasma.

#### Human prostate cancer xenografts

All animal experiments conform to regulatory and ethical standards and were approved by the University of British Columbia Animal Care Committee (A18-0077). Prior to any surgery, metaCAM (1 mg/kg, 0.05 ml/10 g of body weight) was administered subcutaneously. Isoflurane was used as the anesthetic. Animals euthanized by CO_2_. Six to eight-weeks-old male mice (NOD-scid IL2Rgamma^null^) were maintained in the Animal Care Facility at the British Columbia Cancer Research Centre. Five million LNCaP cells were inoculated subcutaneously in a 1:1 volume of matrigel (Corning Discovery Labware, Corning, NY). Tumor volume was measured daily with the aid of digital calipers and calculated by the formula for an ovoid: length × width × height × 0.5236. When xenograft volumes were approximately 100 mm^3^, the mice were castrated with dosing starting weeks later. Animals were dosed daily by oral gavage with 30 mg/kg body weight of 1ae, 10 mg/kg body weight enzalutamide, or vehicle (5% DMSO/1.5% Tween-80/1% CMC).

### Method details

#### Cloning of constructs

##### GFP-AR FL, V7, and ΔNLS cloning strategy

pEGFPC1AR ΔNLS:

The NLS sequence (RKLKK, corresponding to amino acids 629-633 of AR) of the eGFP-AR fusion protein^73^ was removed from peGFP-C1-AR (Addgene #28235) using the Q5 site-directed mutagenesis kit and primer design tools (New England BioLabs) with the following primer pair:

ΔNLS forward primer:

CTTGGTAATCTGAAACTACAGGAG

ΔNLS reverse primer:

GGCTCCCAGAGTCATCCC

Any clones that were found to contain expansion or shrinkage of either the polyQ or poly G sites in AR were corrected by the exchange of the 1510 b.p. KpnI-KpnI fragment with that of the wild-type AR sequence from peGFP-C1-AR.

pEGFPC1A-V7:

The V7 variant of AR was generated from peGFP-C1-AR using the Q5 site-directed mutagenesis kit and primer design tools (New England BioLabs) with the following primer pair:

V7 forward primer:

GCACCTGAAGATGACCAGGCCCTGAGCCCGGAAGCTGAAGAAA

V7 reverse primer:

TTGCAGTTGCCCACCCTGAACTTCTCTCCCAGAGTCATCCCTGC

Any clones that were found to contain expansion or shrinkage of either the polyQ or poly G sites in AR were corrected by the exchange of the 1510b.p. KpnI-KpnI fragment with that of the wild-type AR sequence from peGFP-C1-AR.

mEGFP constructs:

Monomeric EGFP was subcloned into vectors containing human AR (Addgene #29235) and AR-V7 (Addgene #86856) using Gibson assembly to create mEGFP-AR-FL and mEGFP-AR-V7 (referred to as ‘AD+DBD+NLS’ in Fig. 1B, Supplementary Fig. 2A, B) mammalian expression vectors. AR-V7 contains a 16 aa constitutively active NLS containing exon that replaces the LBD exons in AR-FL^74^. The sequence downstream of the AR activation domain in AR-V7, containing the DBD and NLS, was subcloned into an mEGFP plasmid (Addgene #18696) using Gibson assembly to create the mEGFP-AR-V7-ΔAD (referred to as “DBD+NLS” in Fig. 1B, Supplementary Fig. 2A, B) expression vector.

AR-V7 ΔAD forward primer:

AGTTCGTGACCGCCGCCGGGATCACTCTCGGCATGGACGAGCTGTACAAGCTGATCTGTGGAGATGAAGC

AR-V7 ΔAD reverse primer:

CAATAAACAAGTTGGGCCATGGCGGCCAAGCCTCTACAAATGTGGTATGGC

##### AR tyrosine mutagenesis strategy

Production of YtoS mutants for mammalian expression:

The following sequences were optimized for expression in human cells, synthesized and cloned into the pUC57 plasmid (high copy AmpR) by GenScript Biotech, Netherlands, B.V. To enable simple excision from pUC57 and insertion into pEGFPC1AR derivative plasmids two HindIII sites were included as flanks on the fragments. After digestion with HindIII, the resulting 1722b.p. fragments were excised from TBE agarose gels, purified using the E.Z.N.A.® MicroElute Gel Extraction Kit (Omega Biotech) and ligated into HindIII-cut, CIP-treated and gel purified pEGFPC1AR, pEGFPC1AR ΔNLS or pEGFPC1A-V7 plasmids to produce the YtoS mutants.

22YtoS synthetic gene:

AAGCTTCTAATAGCGCCGTGGACGGCACCATGGAAGTGCAGCTGGGACTGGGCAGAGTGAGCCCCAGACCTCCCAGCAAAACATCCAGAGGAGCCTTCCAAAACCTGTTCCAGAGCGTGCGCGAAGTGATTCAGAACCCCGGCCCTAGACACCCTGAGGCTGCCAGCGCCGCCCCTCCCGGCGCCAGCCTGCTGTTGCTGCAGCAGCAGCAGCAGCAACAGCAGCAGCAGCAGCAGCAGCAGCAGCAGCAGCAGCAGCAGCAACAGGAGACAAGCCCCAGACAGCAGCAGCAGCAGCAGGGCGAGGATGGCAGCCCCCAGGCGCACCGGAGAGGCCCTACCGGCAGCCTCGTGCTGGACGAGGAACAGCAGCCTAGCCAGCCTCAATCCGCCCTTGAGTGCCACCCCGAAAGAGGCTGCGTGCCAGAACCAGGCGCCGCCGTGGCCGCCAGCAAGGGCCTGCCTCAGCAACTGCCTGCCCCTCCAGATGAGGACGACAGCGCCGCCCCTAGCACCCTGAGCCTGCTGGGCCCTACTTTTCCAGGCCTGAGCAGCTGCAGCGCTGATCTGAAGGACATCCTGTCTGAGGCTAGCACCATGCAGCTGCTGCAGCAACAGCAACAAGAGGCCGTTTCTGAGGGCTCGAGCAGCGGACGGGCCAGGGAAGCCAGCGGCGCTCCTACCAGCTCTAAGGACAATTCTCTGGGCGGCACAAGCACCATCAGCGATAACGCCAAGGAACTGTGTAAAGCCGTGAGCGTGTCTATGGGCCTGGGAGTGGAAGCCCTGGAACACCTGAGCCCCGGCGAGCAGCTGAGAGGCGACTGCATGAGTGCACCCCTGCTGGGCGTGCCCCCCGCTGTGCGGCCTACACCTTGTGCCCCTCTGGCCGAGTGCAAGGGGTCTCTGCTGGATGACAGCGCTGGCAAGAGCACCGAGGACACCGCCGAGAGCAGCCCCTTCAAGGGCGGCAGCACAAAGGGCCTGGAGGGAGAAAGCCTTGGCTGTTCTGGATCAGCCGCGGCCGGCTCCTCAGGCACCCTGGAACTGCCTAGCACACTGTCTCTGTCTAAATCCGGCGCCCTGGACGAGGCCGCTGCCTCTCAGTCTAGAGATAGCTCTAACTTCCCCCTGGCTCTCGCTGGCCCCCCCCCTCCTCCGCCACCTCCTCACCCACATGCCAGAATCAAGCTGGAAAACCCTCTGGATAGCGGCTCTGCCTGGGCCGCCGCCGCCGCCCAGTGCAGAAGCGGCGACCTGGCCTCTCTGCACGGCGCCGGCGCCGCTGGACCTGGCTCCGGCTCTCCAAGTGCTGCCGCCAGCAGCTCCTGGCACACCCTGTTCACCGCCGAAGAGGGCCAGCTGAGCGGACCTTGCGGCGGCGGAGGAGGGGGCGGCGGGGGAGGCGGCGGCGGCGGCGGCGGCGGCGGCGGAGGCGGAGGCGGCGAGGCTGGCGCTGTGGCCCCTAGCGGCAGCACCAGACCTCCTCAGGGCCTGGCCGGACAGGAGAGCGACTTCACAGCCCCTGATGTCTGGAGCCCCGGCGGAATGGTGAGCCGGGTGCCCTCCCCTAGCCCCACCTGTGTCAAGAGCGAGATGGGCCCTTGGATGGACAGCTCTAGCGGCCCCTCTGGCGACATGCGGCTGGAGACAGCCCGGGACCACGTGCTGCCTATCGACAGCTCTTTTCCACCTCAGAAGACCTGCCTGATCTGCGGAGACGAAGCTT

14YtoS synthetic gene:

AAGCTTCTAATAGCGCCGTCGACGGCACCATGGAAGTGCAGCTGGGCCTGGGAAGAGTGTCCCCTCGGCCCCCCTCTAAGACCAGCAGGGGCGCTTTTCAGAATCTGTTCCAGAGCGTGCGGGAGGTGATCCAGAACCCTGGCCCAAGACACCCTGAGGCTGCTTCCGCCGCCCCACCTGGTGCGAGCCTGCTGCTCCTTCAGCAGCAGCAGCAACAGCAGCAGCAACAGCAGCAGCAGCAACAGCAGCAACAGCAGCAACAGCAGGAGACCTCTCCTAGACAGCAGCAGCAGCAGCAAGGCGAAGATGGCAGCCCTCAGGCCCACCGGAGAGGCCCCACAGGCTCCCTGGTGCTGGATGAGGAACAGCAACCTAGCCAGCCACAGTCTGCGCTGGAATGCCACCCCGAGCGGGGATGTGTGCCCGAGCCTGGCGCCGCCGTCGCCGCCTCTAAAGGCCTGCCTCAGCAGCTGCCCGCCCCTCCAGACGAGGATGATTCTGCTGCCCCTAGCACACTGAGCCTGCTGGGCCCTACCTTTCCTGGCCTCAGCTCATGCAGCGCCGACCTGAAGGACATCCTGAGCGAGGCCTCCACAATGCAGCTGCTGCAGCAGCAGCAGCAGGAGGCTGTGTCTGAGGGCAGCAGTTCCGGCAGAGCCAGAGAGGCCAGCGGAGCCCCCACCAGCAGCAAGGACAACAGCCTGGGCGGAACCAGCACAATCTCTGATAACGCCAAGGAACTGTGCAAAGCCGTGTCCGTGAGCATGGGCCTGGGCGTGGAAGCCCTGGAACACCTGAGCCCTGGCGAGCAGCTGAGAGGCGACTGCATGAGCGCTCCTCTGCTTGGAGTTCCACCAGCCGTGCGGCCTACCCCTTGCGCCCCTCTGGCCGAGTGCAAGGGCTCCCTGCTGGACGACTCAGCCGGCAAGTCCACCGAAGATACCGCCGAGTCTTCCCCCTTCAAGGGCGGAAGCACAAAGGGCCTGGAGGGTGAGAGCCTGGGCTGTAGTGGCAGCGCCGCCGCCGGGAGCAGCGGCACCCTGGAACTACCTAGCACACTGTCTCTGAGCAAGAGCGGAGCGCTGGACGAGGCCGCCGCATCCCAGAGCAGAGATAGCAGCAACTTCCCCCTGGCCCTGGCCGGCCCTCCTCCTCCCCCTCCACCTCCACATCCTCACGCCCGCATCAAGCTGGAAAACCCCCTGGACTCGGGCTCTGCCTGGGCCGCCGCTGCCGCTCAATGTAGAAGCGGCGACCTGGCCAGCCTGCACGGCGCCGGCGCCGCTGGCCCTGGAAGCGGAAGCCCCAGCGCCGCCGCCAGCTCTAGTTGGCACACACTGTTCACCGCCGAGGAAGGCCAGCTGAGCGGCCCTTGTGGCGGCGGCGGCGGAGGCGGCGGCGGCGGCGGCGGGGGAGGAGGCGGCGGCGGCGGAGGCGGCGGCGGAGAGGCCGGCGCCGTGGCCCCTTACGGCTACACCAGACCCCCACAGGGCCTGGCTGGCCAGGAGAGCGACTTCACCGCCCCTGACGTGTGGTACCCCGGAGGCATGGTGTCCAGAGTGCCCTATCCTAGCCCTACATGCGTGAAGTCTGAAATGGGACCTTGGATGGACTCGTACAGCGGCCCTTACGGCGATATGCGGCTGGAAACCGCTAGAGACCACGTGCTGCCTATCGACTACTACTTCCCCCCTCAGAAAACCTGCCTGATTTGCGGCGACG AAGCTT

8YtoS synthetic gene:

AAGCTTCTAACAGCGCTGTGGACGGCACAATGGAAGTGCAACTGGGCCTGGGGCGCGTGTACCCCAGGCCTCCTTCCAAAACCTACAGAGGCGCCTTCCAGAACCTGTTTCAGAGCGTGAGAGAGGTGATTCAGAATCCTGGCCCTAGACATCCTGAGGCAGCGAGCGCCGCCCCTCCTGGCGCCTCTCTGCTGCTCCTGCAACAGCAACAGCAGCAGCAGCAGCAGCAGCAGCAGCAGCAGCAGCAGCAGCAGCAGCAGCAGCAGGAGACCAGCCCCAGACAGCAACAACAACAGCAGGGCGAGGATGGCAGCCCTCAGGCCCACAGACGGGGACCTACAGGCTACCTGGTGCTGGACGAGGAACAGCAGCCTAGCCAGCCCCAGTCTGCCCTGGAGTGCCACCCCGAGAGAGGCTGCGTGCCAGAGCCTGGCGCCGCCGTGGCCGCCTCTAAGGGCCTGCCCCAGCAGCTGCCTGCCCCCCCTGATGAGGACGACAGCGCTGCCCCTAGCACCCTGAGCCTGTTGGGACCTACCTTCCCTGGACTGTCTAGCTGCAGCGCAGATCTGAAGGACATCCTGAGCGAGGCCTCTACAATGCAGCTGCTGCAGCAGCAACAGCAGGAAGCCGTCAGCGAGGGAAGCTCTTCCGGCAGAGCCCGGGAGGCCAGCGGCGCCCCTACCAGCAGCAAGGACAATTACCTGGGAGGAACAAGCACCATCAGCGACAACGCCAAGGAACTGTGCAAGGCCGTGTCCGTTAGCATGGGCCTGGGCGTGGAAGCCCTGGAACACCTGAGCCCAGGCGAGCAGCTGAGAGGCGACTGCATGTACGCCCCTCTTCTGGGGGTGCCCCCGGCCGTGCGGCCTACCCCTTGCGCCCCTCTGGCCGAATGTAAAGGCTCTTTACTGGATGACAGCGCCGGAAAAAGCACGGAAGATACCGCCGAGTATAGCCCGTTCAAGGGCGGTTATACAAAGGGCCTGGAAGGCGAGAGCCTGGGCTGTAGCGGTAGCGCCGCTGCCGGCAGTAGCGGCACACTCGAACTGCCAAGCACCCTGAGCCTGTACAAGTCCGGCGCCCTGGATGAGGCCGCCGCCTACCAGAGCAGAGATTACTACAACTTCCCTCTGGCTCTGGCCGGACCTCCTCCTCCTCCTCCCCCCCCCCACCCCCACGCCAGAATCAAGCTGGAGAACCCCCTGGACTACGGCTCCGCCTGGGCCGCTGCCGCGGCCCAGTGCAGATACGGCGACCTGGCCTCCCTGCACGGCGCTGGCGCCGCCGGACCTGGAAGTGGCAGCCCATCTGCCGCCGCCAGCTCCAGCTGGCACACCCTGTTCACCGCTGAAGAAGGCCAGCTGTACGGCCCTTGTGGTGGTGGCGGCGGCGGCGGCGGAGGCGGCGGCGGCGGCGGCGGTGGCGGAGGCGGCGGCGGCGGCGGAGAGGCTGGCGCCGTCGCCCCTTCTGGATCTACCAGACCCCCTCAAGGCCTGGCTGGCCAGGAGTCCGACTTCACCGCCCCCGACGTGTGGTCCCCTGGAGGAATGGTGTCTCGGGTGCCCTCACCTTCTCCCACATGCGTGAAGAGCGAGATGGGCCCCTGGATGGACAGCAGCAGCGGCCCTTCTGGAGATATGCGGCTGGAGACAGCCCGGGACCACGTGCTGCCTATCGACTCTAGCTTTCCACCACAAAAGACCTGTCTGATCTGCGGCGATGAA GCTT

Production of YtoS mutants for bacterial expression:

pDEST17 plasmids for AR AD YtoS mutants bacterial recombinant production were synthesized by Thermo with the following ORF sequence (flanked with attB1 and attB2 sequences):

22YtoS synthetic gene:

GAGAACCTGTATTTTCAAGGTATGGAAGTTCAGTTAGGTCTGGGTCGTGTTAGTCCGCGTCCGCCTAGCAAAACCAGCCGTGGTGCATTTCAGAATCCGTTTCAGAGCGTTCGTGAAGTTATCCAGAATCCGGGTCCGCGTCATCCGGAAGCAGCAAGCGCAGCACCGCCTGGTGCAAGCCTGCTGCTGCTTCAGCAGCAACAACAGCAGCAACAGCAACAGCAGCAACAACAACAACAACAGCAGCAACAGCAAGAAACCAGTCCGCGTCAACAGCAACAGCAACAAGGTGAAGATGGTAGTCCGCAGGCACATCGTCGTGGTCCGACCGGTAGCCTGGTTCTGGATGAAGAACAGCAGCCGAGCCAGCCGCAGAGCGCACTGGAATGCCATCCGGAACGTGGTTGTGTGCCGGAACCGGGTGCAGCAGTTGCAGCAAGCAAAGGTCTGCCGCAGCAGCTGCCTGCACCTCCGGATGAAGATGATAGTGCAGCACCGAGCACACTGAGCCTGCTGGGTCCGACCTTTCCGGGTCTGAGCAGCTGTAGCGCAGATCTGAAAGATATTCTGAGCGAAGCAAGCACCATGCAGCTGTTACAACAGCAACAACAAGAAGCAGTTAGCGAAGGTAGCAGCAGCGGTCGTGCACGTGAAGCAAGTGGTGCACCGACCAGCAGCAAAGATAATAGCTTAGGTGGCACCAGCACCATTAGCGATAATGCAAAAGAACTGTGTAAAGCCGTTAGCGTTAGCATGGGTCTGGGTGTTGAAGCACTGGAACATCTGAGTCCGGGTGAACAGCTGCGTGGCGATTGTATGAGCGCTCCGCTGCTGGGTGTTCCGCCTGCAGTTCGTCCGACACCGTGTGCACCGCTGGCAGAATGTAAAGGTAGTCTGCTGGATGATAGCGCAGGTAAAAGCACCGAAGATACCGCAGAAAGCAGCCCGTTTAAAGGTGGTAGCACCAAAGGCCTGGAAGGTGAAAGCCTGGGTTGTAGCGGTAGCGCAGCAGCCGGTAGCAGCGGCACACTGGAACTGCCGAGTACACTGTCACTGAGCAAAAGCGGTGCCCTGGATGAGGCAGCCGCAAGCCAGAGCCGTGATAGCAGCAATTTTCCGCTGGCACTGGCAGGTCCTCCGCCACCTCCTCCGCCTCCGCATCCTCATGCACGTATTAAACTGGAAAATCCGCTGGATAGCGGTAGTGCATGGGCTGCAGCGGCAGCACAGTGTCGTAGCGGTGATCTGGCCAGCCTGCATGGCGCAGGCGCAGCAGGTCCTGGTAGCGGTTCACCGTCAGCCGCAGCAAGCTCAAGCTGGCATACCCTGTTTACAGCAGAAGAAGGTCAGCTGAGCGGTCCGTGTGGTGGTGGCGGTGGCGGAGGCGGTGGTGGCGGAGGTGGTGGTGGTGGTGGCGGTGGCGGTGGTGGTGGTGGTGAAGCCGGTGCAGTTGCACCGAGCGGTAGCACCCGTCCGCCTCAAGGTCTGGCAGGCCAAGAAAGCGATTTTACCGCACCGGATGTTTGGAGCCCTGGTGGTATGGTTAGCCGTGTTCCGAGTCCGTCACCGACCTGTGTTAAAAGCGAAATGGGTCCGTGGATGGATAGCAGCTCAGGTCCGAGCGGTGATATGCGTCTGGAAACCGCACGTGATCATGTTCTGCCGATTGATAGCAGTTTTCCGCCTCAGAAAACATAA

14YtoS synthetic gene:

GAGAACCTGTATTTTCAAGGTATGGAAGTTCAGTTAGGTCTGGGTCGTGTTAGTCCGCGTCCGCCTAGCAAAACCAGCCGTGGTGCATTTCAGAATCCGTTTCAGAGCGTTCGTGAAGTTATCCAGAATCCGGGTCCGCGTCATCCGGAAGCAGCAAGCGCAGCACCGCCTGGTGCAAGCCTGCTGCTGCTTCAGCAGCAACAACAGCAGCAACAGCAACAGCAGCAACAACAACAACAACAGCAGCAACAGCAAGAAACCAGTCCGCGTCAACAGCAACAGCAACAAGGTGAAGATGGTAGTCCGCAGGCACATCGTCGTGGTCCGACCGGTAGCCTGGTTCTGGATGAAGAACAGCAGCCGAGCCAGCCGCAGAGCGCACTGGAATGCCATCCGGAACGTGGTTGTGTGCCGGAACCGGGTGCAGCAGTTGCAGCAAGCAAAGGTCTGCCGCAGCAGCTGCCTGCACCTCCGGATGAAGATGATAGTGCAGCACCGAGCACACTGAGCCTGCTGGGTCCGACCTTTCCGGGTCTGAGCAGCTGTAGCGCAGATCTGAAAGATATTCTGAGCGAAGCAAGCACCATGCAGCTGTTACAACAGCAACAACAAGAAGCAGTTAGCGAAGGTAGCAGCAGCGGTCGTGCACGTGAAGCAAGTGGTGCACCGACCAGCAGCAAAGATAATAGCTTAGGTGGCACCAGCACCATTAGCGATAATGCAAAAGAACTGTGTAAAGCCGTTAGCGTTAGCATGGGTCTGGGTGTTGAAGCACTGGAACATCTGAGTCCGGGTGAACAGCTGCGTGGCGATTGTATGAGCGCTCCGCTGCTGGGTGTTCCGCCTGCAGTTCGTCCGACACCGTGTGCACCGCTGGCAGAATGTAAAGGTAGTCTGCTGGATGATAGCGCAGGTAAAAGCACCGAAGATACCGCAGAAAGCAGCCCGTTTAAAGGTGGTAGCACCAAAGGCCTGGAAGGTGAAAGCCTGGGTTGTAGCGGTAGCGCAGCAGCCGGTAGCAGCGGCACACTGGAACTGCCGAGTACACTGTCACTGAGCAAAAGCGGTGCCCTGGATGAGGCAGCCGCAAGCCAGAGCCGTGATAGCAGCAATTTTCCGCTGGCACTGGCAGGTCCTCCGCCACCTCCTCCGCCTCCGCATCCTCATGCACGTATTAAACTGGAAAATCCGCTGGATAGCGGTAGTGCATGGGCTGCAGCGGCAGCACAGTGTCGTAGCGGTGATCTGGCCAGCCTGCATGGCGCAGGCGCAGCAGGTCCTGGTAGCGGTTCACCGTCAGCCGCAGCAAGCTCAAGCTGGCATACCCTGTTTACAGCAGAAGAAGGTCAGCTGAGCGGTCCGTGTGGTGGTGGCGGTGGCGGAGGCGGTGGTGGCGGAGGTGGTGGTGGTGGTGGCGGTGGCGGTGGTGGTGGTGGTGAAGCCGGTGCAGTTGCACCGTATGGTTATACCCGTCCGCCTCAAGGTCTGGCAGGCCAAGAAAGCGATTTTACCGCACCGGATGTTTGGTATCCTGGTGGTATGGTTAGCCGTGTTCCGTATCCGAGTCCGACCTGTGTTAAAAGCGAAATGGGTCCGTGGATGGATAGCTATAGTGGTCCGTATGGTGATATGCGTCTGGAAACCGCACGTGATCATGTTCTGCCGATTGATTATTACTTTCCGCCTCAGAAAACGTAA

8YtoS synthetic gene:

GAGAACCTGTATTTTCAAGGTATGGAAGTTCAGTTAGGTCTGGGTCGTGTTTATCCGCGTCCGCCTAGCAAAACCTATCGTGGTGCATTTCAGAATCCGTTTCAGAGCGTTCGTGAAGTTATCCAGAATCCGGGTCCGCGTCATCCGGAAGCAGCAAGCGCAGCACCGCCTGGTGCAAGCCTGCTGCTGCTTCAGCAGCAACAACAGCAGCAACAGCAACAGCAGCAACAACAACAACAACAGCAGCAACAGCAAGAAACCAGTCCGCGTCAACAGCAACAGCAACAAGGTGAAGATGGTAGTCCGCAGGCACATCGTCGTGGTCCGACCGGTTATCTGGTTCTGGATGAAGAACAGCAGCCGAGCCAGCCGCAGAGCGCACTGGAATGCCATCCGGAACGTGGTTGTGTGCCGGAACCGGGTGCAGCAGTTGCAGCAAGCAAAGGTCTGCCGCAGCAGCTGCCTGCACCTCCGGATGAAGATGATAGTGCAGCACCGAGCACACTGAGCCTGCTGGGTCCGACCTTTCCGGGTCTGAGCAGCTGTAGCGCAGATCTGAAAGATATTCTGAGCGAAGCAAGCACCATGCAGCTGTTACAACAGCAACAACAAGAAGCAGTTAGCGAAGGTAGCAGCAGCGGTCGTGCACGTGAAGCAAGTGGTGCACCGACCAGCAGCAAAGATAACTATTTAGGTGGCACCAGCACCATTAGCGATAATGCAAAAGAACTGTGTAAAGCCGTTAGCGTTAGCATGGGTCTGGGTGTTGAAGCACTGGAACATCTGAGTCCGGGTGAACAGCTGCGTGGCGATTGTATGTATGCTCCGCTGCTGGGTGTTCCGCCTGCAGTTCGTCCGACACCGTGTGCACCGCTGGCAGAATGTAAAGGTAGTCTGCTGGATGATAGCGCAGGTAAAAGCACCGAAGATACCGCAGAATATTCACCGTTTAAAGGTGGTTATACCAAAGGCCTGGAAGGTGAAAGCCTGGGTTGTAGCGGTAGCGCAGCAGCCGGTAGCAGCGGCACACTGGAACTGCCGAGTACACTGTCACTGTATAAAAGCGGTGCCCTGGATGAGGCAGCAGCATATCAGAGCCGTGATTATTACAATTTTCCGCTGGCACTGGCAGGTCCGCCTCCGCCTCCACCACCGCCTCATCCGCATGCACGTATTAAACTGGAAAATCCGCTGGATTATGGTAGCGCATGGGCAGCCGCAGCCGCACAGTGTCGTTATGGTGATCTGGCCAGCCTGCATGGCGCTGGTGCAGCCGGTCCTGGTAGCGGTTCACCGAGTGCAGCCGCAAGCAGCTCATGGCATACCCTGTTTACAGCCGAAGAGGGTCAGCTGTATGGTCCGTGTGGTGGCGGAGGTGGCGGTGGTGGTGGTGGCGGAGGCGGTGGCGGAGGTGGTGGCGGTGGCGGAGGTGGCGGTGAAGCTGGTGCAGTTGCACCGAGCGGTAGCACCCGTCCGCCTCAAGGTCTGGCAGGCCAAGAAAGCGATTTTACCGCACCGGATGTTTGGAGCCCTGGTGGTATGGTTAGCCGTGTTCCGAGTCCGTCACCGACCTGTGTTAAAAGCGAAATGGGTCCGTGGATGGATAGCAGCTCAGGTCCGAGCGGTGATATGCGTCTGGAAACCGCACGTGATCATGTTCTGCCGATTGATAGCAGTTTTCCGCCTCAGAAAACATAA

##### AR helix breaking mutagenesis strategy

pDONR221-AR-AD-WT:

The DNA sequence corresponding to the 1-558 amino acid fragment of AR-AD was synthesized and encoded in a pDONR221 vector by Thermo using the following ORF sequence (flanked with attB1 and attB2 sequences):

GAGAACCTGTATTTTCAGGGTATGGAAGTTCAGCTGGGTCTGGGTCGTGTTTATCCGCGTCCGCCTAGCAAAACCTATCGTGGTGCATTTCAGAACCTGTTTCAGAGCGTTCGTGAAGTTATTCAGAATCCGGGTCCGCGTCATCCGGAAGCAGCAAGCGCAGCACCGCCTGGTGCAAGCTTACTGCTGCTGCAACAGCAACAGCAGCAGCAACAACAGCAGCAACAGCAACAACAACAACAACAGCAGCAGCAAGAAACCAGTCCGCGTCAACAGCAACAGCAACAGGGTGAAGATGGTAGTCCGCAGGCACATCGTCGTGGTCCGACCGGTTATCTGGTTCTGGATGAAGAACAGCAGCCGAGCCAGCCGCAGAGCGCACTGGAATGCCATCCGGAACGTGGTTGTGTGCCGGAACCGGGTGCAGCAGTTGCAGCAAGCAAAGGTCTGCCGCAGCAGCTGCCTGCACCTCCGGATGAAGATGATAGTGCAGCACCGAGCACCCTGAGCCTGCTGGGTCCGACCTTTCCGGGTCTGAGCAGCTGTAGCGCAGATCTGAAAGATATTCTGAGCGAAGCAAGCACCATGCAGCTGCTGCAACAACAGCAACAAGAAGCAGTTAGCGAAGGTAGCAGCAGCGGTCGTGCACGTGAAGCAAGTGGTGCACCGACCAGCAGCAAAGATAACTATCTGGGTGGCACCAGCACCATTAGCGATAATGCAAAAGAACTGTGTAAAGCCGTTAGCGTTAGCATGGGCCTGGGTGTTGAAGCACTGGAACATCTGAGTCCGGGTGAACAGCTGCGTGGTGATTGTATGTATGCTCCGCTGCTGGGTGTTCCGCCTGCAGTTCGTCCGACCCCGTGTGCACCGCTGGCAGAATGTAAAGGTAGTCTGCTGGATGATAGCGCAGGTAAAAGCACCGAAGATACCGCAGAATATTCACCGTTTAAAGGTGGTTATACCAAAGGCCTGGAAGGTGAAAGCCTGGGTTGTAGCGGTAGCGCAGCAGCCGGTAGCAGCGGCACACTGGAACTGCCGAGTACCCTGTCACTGTATAAAAGCGGTGCCCTGGATGAGGCAGCAGCATATCAGAGCCGTGATTATTACAATTTTCCGCTGGCACTGGCAGGTCCGCCTCCGCCTCCACCACCGCCTCATCCGCATGCACGTATTAAACTGGAAAATCCGCTGGATTATGGTAGCGCATGGGCAGCCGCAGCCGCACAGTGTCGTTATGGTGATCTGGCAAGCCTGCATGGCGCTGGTGCAGCCGGTCCGGGTAGCGGTTCACCTAGTGCAGCCGCAAGCAGCTCATGGCATACCCTGTTTACAGCCGAAGAGGGTCAGCTGTATGGTCCGTGTGGTGGGGGTGGCGGAGGGGGAGGGGGTGGGGGAGGTGGGGGTGGTGGCGGTGGGGGTGGGGGTGGTGGTGGCGAAGCTGGTGCAGTTGCACCGTATGGTTATACCCGTCCGCCTCAGGGTCTGGCAGGCCAAGAAAGCGATTTTACCGCACCGGATGTTTGGTATCCGGGTGGTATGGTTAGCCGTGTTCCGTATCCGAGCCCGACCTGTGTTAAAAGCGAAATGGGTCCGTGGATGGATAGCTATAGCGGTCCGTATGGTGATATGCGTCTGGAAACCGCACGTGATCATGTTCTGCCGATTGATTACTATTTCCCTCCGCAGAAAACCTAATAA

pDEST17-AR-AD-WT:

pDONR221-AR-AD-WT was subcloned into a pDEST17 vector by using LP clonase reaction (Thermo).

pDEST17-AR-AD-WT*:

The L26P mutation was introduced into a wild-type AR sequence (pDONR221-AR-AD-WT) using a Quickchange™ protocol with Pfu Turbo polymerase (Agilent) and the following primer pairs:

L26P forward primer:

TGGTGCATTTCAGAACCCGTTTCAGAGCGTTCGTG

L26P reverse primer:

CACGAACGCTCTGAAACGGGTTCTGAAATGCACCA

The resulting plasmid incorporating L26P mutation (pDONR221-AR-AD-WT*) was subcloned into a pDEST17 vector by using LP clonase reaction (Thermo).

pDEST17-AR-AD-L56P*:

L56P mutation was introduced into the pDONR221-AR-AD-WT* (bearing L26P mutation and previously described) using a Quickchange™ protocol with Pfu Turbo polymerase (Agilent) and the following primer pairs to generate pDONR221-AR-AD-L56P*:

L56P forward primer:

GCAAGCTTACTGCTGCCGCAACAGCAACAGCAG

L56P reverse primer:

CTGCTGTTGCTGTTGCGGCAGCAGTAAGCTTGC

The resulting plasmid incorporating L26P and L56P mutations (pDONR221-AR-AD-L56P*) was subcloned into a pDEST17 vector by using LP clonase reaction (Thermo).

pDEST17-AR-AD-Tau-1*:

The A186P, L192P, C238P mutations were introduced step-wise into pDONR221-AR-AD-WT* (bearing L26P mutation and previously described) using a Quickchange™ protocol with Pfu Turbo polymerase (Agilent) and the following primer pairs to generate

pDONR221-AR-AD-Tau-1*:

A186P, L192P forward primer:

GATATTCTGAGCGAACCAAGCACCATGCAGCTGCCGCAACAACAGCAACA

A186P, L192P reverse primer:

TTGCTGTTGTTGCGGCAGCTGCATGGTGCTTGGTTCGCTCAGAATATCTT

C238P forward primer:

TAGCGATAATGCAAAAGAACTGCCGAAAGCCGTTAGCGTTAGCATGG

C238P reverse primer:

CCATGCTAACGCTAACGGCTTTCGGCAGTTCTTTTGCATTATCGCTA

The resulting plasmid incorporating L26P, A186P, L192P and C238P mutations (pDONR221-AR-AD-Tau-1*) was subcloned into a pDEST17 vector by using LP clonase reaction (Thermo).

pDEST17-AR-AD-Tau-5*:

The A356P, A398P and T435P mutations were introduced step-wise into pDONR221-AR-AD-WT* (bearing L26P mutation and previously described) using a Quickchange™ protocol with Pfu Turbo polymerase (Agilent) and the following primer pairs to generate pDONR221-AR-AD-Tau-5*:

A356P forward primer:

CCTGGATGAGGCAGCACCGTATCAGAGCCGTGATT

A356P reverse primer:

AATCACGGCTCTGATACGGTGCTGCCTCATCCAGG

A398P forward primer:

GGTAGCGCATGGCCGGCCGCAGCCGCA

A398P reverse primer:

TGCGGCTGCGGCCGGCCATGCGCTACC

T435P forward primer:

CAAGCAGCTCATGGCATCCGCTGTTTACAGCCGAAG

T435P reverse primer:

CTTCGGCTGTAAACAGCGGATGCCATGAGCTGCTTG

The resulting plasmid incorporating L26P, A356P, A398P and T435P mutations (pDONR221-AR-AD-Tau-5*) was subcloned into a pDEST17 vector by using LP clonase reaction (Thermo).

pDEST17-AR-AD-L56P+Tau-1+Tau-5*:

The L56P, A186P, L192P, C238P mutations were introduced step-wise into pDONR221-AR-AD-TAU-5* (bearing L26P, A186P, L192P and C238P mutation and previously described) using a Quickchange™ protocol with Pfu Turbo polymerase (Agilent) and the following primer pairs to generate pDONR221-AR-AD-L56P+Tau-1+Tau-5*:

L56P forward primer:

GCAAGCTTACTGCTGCCGCAACAGCAACAGCAG

L56P reverse primer:

CTGCTGTTGCTGTTGCGGCAGCAGTAAGCTTGC

A186P, L192P forward primer:

GATATTCTGAGCGAACCAAGCACCATGCAGCTGCCGCAACAACAGCAACA

A186P, L192P reverse primer:

TTGCTGTTGTTGCGGCAGCTGCATGGTGCTTGGTTCGCTCAGAATATCTT

C238P forward primer:

TAGCGATAATGCAAAAGAACTGCCGAAAGCCGTTAGCGTTAGCATGG

C238P reverse primer:

CCATGCTAACGCTAACGGCTTTCGGCAGTTCTTTTGCATTATCGCTA

The resulting plasmid incorporating L26P, L56P, A186P, L192P, C238P, A356P, A398P and T435P mutations (pDONR221-AR-AD-L56P+Tau-1+Tau-5*) was subcloned into a pDEST17 vector by using LP clonase reaction (Thermo).

eGFP-AR-ΔNLS-Δ21-35:

A 507b.p. fragment containing Δ21-35 was amplified from pCMV5-FLAG-AR deltaFQNLF ^36^ using KOD polymerase (Merck Millipore), the supplied buffer#2 and the following primers: ARΔ21-35 forward primer: AATTCTGCAGTCGACGGTACCATGGAAGTGCAGTTAGGGCTGGGAAGGGTC

ARΔ21-35 reverse primer:

GGAGCAGCTGCTTAAGCCGGGGAAAGTGG

The resulting fragment was purified using AmPure XT (Beckman) before InFusion (Takara Bio) into SalI and AflII-cut and gel purified pEGFPC1AR ΔNLS plasmid.

eGFP-AR-ΔNLS-Tau-1:

The A186P, L192P, C238P mutations were introduced step-wise into a wild-type AR sequence encoded in pDONR221-AR-AD-WT using a Quickchange™ protocol with Pfu Turbo polymerase (Agilent) and the following primer pairs:

A186P, L192P forward primer:

GATATTCTGAGCGAACCAAGCACCATGCAGCTGCCGCAACAACAGCAACA

A186P, L192P reverse primer:

TTGCTGTTGTTGCGGCAGCTGCATGGTGCTTGGTTCGCTCAGAATATCTT

C238P forward primer:

TAGCGATAATGCAAAAGAACTGCCGAAAGCCGTTAGCGTTAGCATGG

C238P reverse primer:

CCATGCTAACGCTAACGGCTTTCGGCAGTTCTTTTGCATTATCGCTA

A 755bp fragment was amplified from the resulting clone incorporating A186P, L192P, C238P mutations (pDONR221-AR-AD-TAU1) using KOD polymerase (Takara Bio) and the following primer pair.

Tau-1 forward primer:

CTTTCCCCGGCTTAAGCAGCTGTAGCGCAGATCTGAAAGATATTCTGAGCG

Tau-1 reverse primer:

GCGGCTGAGGGTGACCCGCTACCCGGACCGGCTGCAC

The resulting fragment was digested with DpnI to remove template, and purified using AmPure XT (Beckman) before InFusion into AflII-BstEII-cut and gel-purified pEGFPC1-ARΔNLS plasmid.

eGFP-AR-ΔNLS-Tau-5:

The A356P, A398P and T435P mutations were introduced step-wise into a wild-type AR sequence (pDONR221-AR-AD-WT) using a Quickchange™ protocol with Pfu Turbo polymerase (Agilent) and the following primer pairs:

A356P forward primer:

CCTGGATGAGGCAGCACCGTATCAGAGCCGTGATT

A356P reverse primer:

AATCACGGCTCTGATACGGTGCTGCCTCATCCAGG

A398P forward primer:

GGTAGCGCATGGCCGGCCGCAGCCGCA

A398P reverse primer:

TGCGGCTGCGGCCGGCCATGCGCTACC

T435P forward primer:

CAAGCAGCTCATGGCATCCGCTGTTTACAGCCGAAG

T435P reverse primer:

CTTCGGCTGTAAACAGCGGATGCCATGAGCTGCTTG

A 1544bp fragment was then amplified from the resulting plasmid incorporating A356P, A398P and T435P mutations (pDONR221-AR-AD-TAU-5) using KOD polymerase (Takara Bio) and the following primer pair.

Tau-5 forward primer:

ATTCTGCAGTCGACGGTACCATGGAAGTTCAGCTGGGTCTGGGTCGTG

Tau-5 reverse primer:

CACCATGCCGCCAGGGTACCAAACATCCGGTGCGGTAAAATCGCTTTCTTGGC

The resulting fragment was digested with DpnI to remove template, and purified using AmPure XT (Beckman) before InFusion into KpnI-cut and gel-purified pEGFPC1-ARΔNLS plasmid.

##### BioID plasmid generation strategy

Constructs to express FLAG-MTID or its fusions to AR WT and 22YtoS fusion proteins were synthesized by Genscript and either cloned into pcDNA3.1(-) and subsequently cloned into pLenti-CMV-MCS-GFP-SV-puro using XbaI and BamHI to replace GFP or cloned directly into pLenti-CMV-MCS-GFP-SV-puro by Genscript using the same sites. Sequences were codon optimized for mammalian expression and verified by sequencing.

pLenti-CMV-MCS-GFP-SV-puro was a gift from Paul Odgren (Addgene plasmid # 73582) FLAG-MTID synthetic gene:

TCTAGAGCGCTGCCACCATGGATTACAAGGATGACGACGATAAGATCCGCATCCCGCTGCT GAACGCTAAACAGATTCTGGGACAGCTGGACGGCGGGAGCGTGGCAGTCCTGCCTGTGGT CGACTCCACCAATCAGTACCTGCTGGATCGAATCGGCGAGCTGAAGAGTGGGGATGCTTG CATTGCAGAATATCAGCAGGCAGGGAGAGGAAGCAGAGGGAGGAAATGGTTCTCTCCTTTT GGAGCTAACCTGTACCTGAGTATGTTTTGGCGCCTGAAGCGGGGACCAGCAGCAATCGGC CTGGGCCCGGTCATCGGAATTGTCATGGCAGAAGCGCTGCGAAAGCTGGGAGCAGACAAG GTGCGAGTCAAATGGCCCAATGACCTGTATCTGCAGGATAGAAAGCTGGCAGGCATCCTGG TGGAGCTGGCCGGAATAACAGGCGATGCTGCACAGATCGTCATTGGCGCCGGGATTAACG TGGCTATGAGGCGCGTGGAGGAAAGCGTGGTCAATCAGGGCTGGATCACACTGCAGGAAG CAGGGATTAACCTGGACAGGAATACTCTGGCCGCTATGCTGATCCGAGAGCTGCGGGCAG CCCTGGAACTGTTCGAGCAGGAAGGCCTGGCTCCATATCTGTCACGGTGGGAGAAGCTGG ATAACTTCATCAATAGACCCGTGAAGCTGATCATTGGGGACAAAGAGATTTTCGGGATTAGC CGGGGGATTGATAAACAGGGAGCCCTGCTGCTGGAACAGGACGGAGTTATCAAACCCTGG ATGGGCGGAGAAATCAGTCTGCGGTCTGCCGAAAAGTGAGGATCC

FLAG-MTID-AR-WT synthetic gene:

TCTAGAGCGCTGCCACCATGGACTACAAGGACGATGACGATAAGATCAGAATCCCCCTGCT GAACGCCAAGCAGATCCTGGGACAGCTGGATGGAGGCTCTGTGGCCGTGCTGCCAGTGG TGGACAGCACCAATCAGTACCTGCTGGATAGGATCGGCGAGCTGAAGAGCGGCGACGCCT GCATCGCCGAGTATCAGCAGGCAGGAAGGGGCTCTCGGGGAAGAAAGTGGTTCAGCCCAT TTGGCGCCAACCTGTACCTGTCCATGTTCTGGCGGCTGAAGAGAGGACCAGCAGCAATCG GACTGGGACCTGTGATCGGCATCGTGATGGCAGAGGCCCTGAGGAAGCTGGGAGCAGAC AAGGTGAGAGTGAAGTGGCCCAATGACCTGTATCTGCAGGATAGGAAGCTGGCAGGCATC CTGGTGGAGCTGGCAGGAATCACCGGCGATGCAGCACAGATCGTGATCGGAGCAGGCATC AACGTGGCAATGAGGAGAGTGGAGGAGAGCGTGGTGAATCAGGGATGGATCACCCTGCAG GAGGCAGGCATCAACCTGGATCGCAATACACTGGCAGCAATGCTGATCAGGGAGCTGAGG GCCGCCCTGGAGCTGTTTGAGCAGGAGGGCCTGGCCCCATACCTGTCTAGGTGGGAGAA GCTGGACAACTTCATCAATCGCCCCGTGAAGCTGATCATCGGCGATAAGGAGATCTTTGGC ATCTCCAGAGGCATCGACAAGCAGGGCGCCCTGCTGCTGGAGCAGGATGGCGTGATCAAG CCTTGGATGGGAGGAGAGATCAGCCTGAGGTCCGCCGAGAAGGAGGTGCAGCTGGGACT GGGACGGGTGTACCCAAGACCACCTAGCAAGACCTATCGCGGCGCCTTCCAGAACCTGTT TCAGTCCGTGCGGGAAGTGATCCAGAATCCAGGCCCCAGACACCCAGAGGCAGCATCCGC CGCACCACCAGGAGCATCTCTGTTATTACTGCAACAACAGCAACAACAGCAACAGCAACAG CAGCAACAACAGCAGCAACAGCAACAACAACAGCAGGAGACATCTCCTAGGCAGCAGCAG CAGCAGCAGGGAGAGGACGGCAGCCCACAGGCACACAGGAGGGGACCCACCGGCTACC TGGTGCTGGATGAGGAGCAGCAGCCATCCCAGCCACAGTCTGCCCTGGAGTGCCACCCAG AGAGAGGCTGCGTGCCTGAGCCAGGAGCAGCAGTGGCAGCCAGCAAGGGCCTGCCCCA GCAGCTGCCTGCACCTCCAGACGAGGACGATTCCGCCGCACCATCTACCCTGAGCCTGCT GGGCCCTACATTCCCAGGACTGAGCTCCTGCTCCGCCGACCTGAAGGATATCCTGTCCGA GGCCTCTACAATGCAGCTGCTGCAGCAGCAGCAGCAGGAGGCCGTGTCTGAGGGCTCTAG CTCCGGAAGGGCAAGGGAGGCAAGCGGAGCACCCACCTCTAGCAAGGACAACTACCTGG GCGGCACCAGCACAATCTCCGATAATGCCAAGGAGCTGTGCAAGGCCGTGTCCGTGTCTAT GGGACTGGGAGTGGAGGCCCTGGAGCACCTGAGCCCAGGAGAGCAGCTGAGGGGCGAC TGTATGTATGCCCCTCTGCTGGGAGTGCCACCTGCCGTGCGCCCAACACCTTGCGCACCA CTGGCAGAGTGTAAGGGCTCCCTGCTGGACGATAGCGCCGGCAAGTCCACCGAGGATACA GCCGAGTACTCCCCTTTCAAGGGCGGCTATACCAAGGGCCTGGAGGGCGAGTCTCTGGGA TGTAGCGGCTCCGCCGCAGCAGGCTCCTCTGGCACCCTGGAGCTGCCATCTACACTGAGC CTGTACAAGTCCGGCGCCCTGGACGAGGCAGCAGCCTATCAGTCTAGGGATTACTATAACT TTCCCCTGGCCCTGGCAGGACCTCCTCCACCACCACCTCCACCACACCCACACGCACGGA TCAAGCTGGAGAATCCTCTGGACTACGGCTCTGCCTGGGCAGCAGCAGCAGCACAGTGCA GATATGGCGATCTGGCAAGCCTGCACGGAGCAGGAGCAGCAGGCCCAGGCTCTGGCAGC CCCTCCGCCGCCGCCAGCTCCTCTTGGCACACCCTGTTCACAGCCGAGGAGGGCCAGCT GTACGGCCCCTGTGGGGGCGGCGGGGGCGGCGGCGGCGGAGGCGGCGGAGGAGGCGG AGGGGGCGGAGGAGGCGGCGGCGGCGAGGCAGGAGCAGTGGCACCTTACGGATATACCA GGCCTCCACAGGGACTGGCAGGACAGGAGAGCGACTTTACAGCCCCTGACGTGTGGTACC CAGGCGGCATGGTGAGCAGAGTGCCATATCCCTCCCCTACCTGCGTGAAGTCTGAGATGG GCCCTTGGATGGACTCTTACAGCGGCCCATATGGCGATATGAGGCTGGAGACCGCAAGGG ACCACGTGCTGCCCATCGATTACTATTTCCCCCCTCAGAAGACATGCCTGATCTGTGGCGAC GAGGCAAGCGGATGCCACTACGGCGCCCTGACCTGCGGCTCCTGTAAGGTGTTCTTTAAG CGGGCCGCCGAGGGCAAGCAGAAGTATCTGTGCGCCTCCAGAAACGACTGTACAATCGAT AAGTTTCGGAGAAAGAATTGCCCTTCTTGTCGGCTGAGAAAGTGTTACGAGGCAGGAATGA CCCTGGGAGCAAGGAAGCTGAAGAAGCTGGGCAACCTGAAGCTGCAGGAGGAGGGAGAG GCAAGCTCCACCACATCCCCAACCGAGGAGACCACACAGAAGCTGACAGTGTCTCACATC GAGGGCTATGAGTGCCAGCCTATCTTCCTGAATGTGCTGGAGGCAATCGAGCCAGGAGTG GTGTGCGCAGGCCACGACAACAATCAGCCTGATAGCTTTGCCGCCCTGCTGTCTAGCCTGA ACGAGCTGGGAGAGAGGCAGCTGGTGCACGTGGTGAAGTGGGCCAAGGCCCTGCCAGG CTTCAGAAATCTGCACGTGGACGATCAGATGGCCGTGATCCAGTACTCCTGGATGGGCCTG ATGGTGTTCGCCATGGGCTGGAGGAGCTTTACAAACGTGAACAGCCGGATGCTGTATTTCG CCCCTGACCTGGTGTTTAACGAGTACCGGATGCACAAGAGCCGGATGTATAGCCAGTGCGT GAGGATGCGCCACCTGAGCCAGGAGTTCGGCTGGCTGCAGATCACCCCTCAGGAGTTCCT GTGCATGAAGGCCCTGCTGCTGTTTTCCATCATCCCAGTGGACGGCCTGAAGAACCAGAAG TTCTTTGATGAGCTGAGGATGAATTACATCAAGGAGCTGGACAGGATCATCGCCTGCAAGC GCAAGAACCCCACCTCCTGTTCTAGGCGCTTTTATCAGCTGACAAAGCTGCTGGATAGCGT GCAGCCTATCGCAAGGGAGCTGCACCAGTTCACATTTGACCTGCTGATCAAGTCCCACATG GTGTCTGTGGATTTCCCCGAGATGATGGCCGAGATCATCAGCGTGCAGGTGCCAAAGATCC TGTCCGGCAAGGTGAAGCCCATCTACTTTCACACCCAGTGAGGATCC

FLAG-MTID-AR-WT-Y22toS synthetic gene:

TCTAGAGCGCTGCCACCATGGACTACAAGGACGATGACGATAAGATCAGAATCCCCCTGCTGAACGCCAAGCAGATCCTGGGACAGCTGGATGGAGGCTCTGTGGCCGTGCTGCCAGTGGTGGACAGCACCAATCAGTACCTGCTGGATAGGATCGGCGAGCTGAAGAGCGGCGACGCCTGCATCGCCGAGTATCAGCAGGCAGGAAGGGGCTCTCGGGGAAGAAAGTGGTTCAGCCCATTTGGCGCCAACCTGTACCTGTCCATGTTCTGGCGGCTGAAGAGAGGACCAGCAGCAATCGGACTGGGACCTGTGATCGGCATCGTGATGGCAGAGGCCCTGAGGAAGCTGGGAGCAGACAAGGTGAGAGTGAAGTGGCCCAATGACCTGTATCTGCAGGATAGGAAGCTGGCAGGCATCCTGGTGGAGCTGGCAGGAATCACCGGCGATGCAGCACAGATCGTGATCGGAGCAGGCATCAACGTGGCAATGAGGAGAGTGGAGGAGAGCGTGGTGAATCAGGGATGGATCACCCTGCAGGAGGCAGGCATCAACCTGGATCGCAATACACTGGCAGCAATGCTGATCAGGGAGCTGAGGGCCGCCCTGGAGCTGTTTGAGCAGGAGGGCCTGGCCCCATACCTGTCTAGGTGGGAGAAGCTGGACAACTTCATCAATCGCCCCGTGAAGCTGATCATCGGCGATAAGGAGATCTTTGGCATCTCCAGAGGCATCGACAAGCAGGGCGCCCTGCTGCTGGAGCAGGATGGCGTGATCAAGCCTTGGATGGGAGGAGAGATCAGCCTGAGGTCCGCCGAGAAGGAGGTGCAGCTGGGACTGGGACGGGTGTCCCCAAGACCACCTAGCAAGACCTCTCGCGGCGCCTTCCAGAACCTGTTTCAGTCCGTGCGGGAAGTGATCCAGAATCCAGGCCCCAGACACCCAGAGGCAGCATCCGCCGCACCACCAGGAGCATCTCTGTTATTACTGCAACAACAGCAACAACAGCAACAGCAACAGCAGCAACAACAGCAGCAACAGCAACAACAACAGCAGGAGACATCTCCTAGGCAGCAGCAGCAGCAGCAGGGAGAGGACGGCAGCCCACAGGCACACAGGAGGGGACCCACCGGCTCCCTGGTGCTGGATGAGGAGCAGCAGCCATCCCAGCCACAGTCTGCCCTGGAGTGCCACCCAGAGAGAGGCTGCGTGCCTGAGCCAGGAGCAGCAGTGGCAGCCAGCAAGGGCCTGCCCCAGCAGCTGCCTGCACCTCCAGACGAGGACGATTCCGCCGCACCATCTACCCTGAGCCTGCTGGGCCCTACATTCCCAGGACTGAGCTCCTGCTCCGCCGACCTGAAGGATATCCTGTCCGAGGCCTCTACAATGCAGCTGCTGCAGCAGCAGCAGCAGGAGGCCGTGTCTGAGGGCTCTAGCTCCGGAAGGGCAAGGGAGGCAAGCGGAGCACCCACCTCTAGCAAGGACAACTCCCTGGGCGGCACCAGCACAATCTCCGATAATGCCAAGGAGCTGTGCAAGGCCGTGTCCGTGTCTATGGGACTGGGAGTGGAGGCCCTGGAGCACCTGAGCCCAGGAGAGCAGCTGAGGGGCGACTGTATGTCTGCCCCTCTGCTGGGAGTGCCACCTGCCGTGCGCCCAACACCTTGCGCACCACTGGCAGAGTGTAAGGGCTCCCTGCTGGACGATAGCGCCGGCAAGTCCACCGAGGATACAGCCGAGTCCTCCCCTTTCAAGGGCGGCTCTACCAAGGGCCTGGAGGGCGAGTCTCTGGGATGTAGCGGCTCCGCCGCAGCAGGCTCCTCTGGCACCCTGGAGCTGCCATCTACACTGAGCCTGTCCAAGTCCGGCGCCCTGGACGAGGCAGCAGCCTCTCAGTCTAGGGATTCCTCTAACTTTCCCCTGGCCCTGGCAGGACCTCCTCCACCACCACCTCCACCACACCCACACGCACGGATCAAGCTGGAGAATCCTCTGGACTCCGGCTCTGCCTGGGCAGCAGCAGCAGCACAGTGCAGATCTGGCGATCTGGCAAGCCTGCACGGAGCAGGAGCAGCAGGCCCAGGCTCTGGCAGCCCCTCCGCCGCCGCCAGCTCCTCTTGGCACACCCTGTTCACAGCCGAGGAGGGCCAGCTGTCCGGCCCCTGTGGGGGCGGCGGGGGCGGCGGCGGCGGAGGCGGCGGAGGAGGCGGAGGGGGCGGAGGAGGCGGCGGCGGCGAGGCAGGAGCAGTGGCACCTTCCGGATCTACCAGGCCTCCACAGGGACTGGCAGGACAGGAGAGCGACTTTACAGCCCCTGACGTGTGGTCCCCAGGCGGCATGGTGAGCAGAGTGCCATCTCCCTCCCCTACCTGCGTGAAGTCTGAGATGGGCCCTTGGATGGACTCTTCCAGCGGCCCATCTGGCGATATGAGGCTGGAGACCGCAAGGGACCACGTGCTGCCCATCGATTCCTCTTTCCCCCCTCAGAAGACATGCCTGATCTGTGGCGACGAGGCAAGCGGATGCCACTACGGCGCCCTGACCTGCGGCTCCTGTAAGGTGTTCTTTAAGCGGGCCGCCGAGGGCAAGCAGAAGTATCTGTGCGCCTCCAGAAACGACTGTACAATCGATAAGTTTCGGAGAAAGAATTGCCCTTCTTGTCGGCTGAGAAAGTGTTACGAGGCAGGAATGACCCTGGGAGCAAGGAAGCTGAAGAAGCTGGGCAACCTGAAGCTGCAGGAGGAGGGAGAGGCAAGCTCCACCACATCCCCAACCGAGGAGACCACACAGAAGCTGACAGTGTCTCACATCGAGGGCTATGAGTGCCAGCCTATCTTCCTGAATGTGCTGGAGGCAATCGAGCCAGGAGTGGTGTGCGCAGGCCACGACAACAATCAGCCTGATAGCTTTGCCGCCCTGCTGTCTAGCCTGAACGAGCTGGGAGAGAGGCAGCTGGTGCACGTGGTGAAGTGGGCCAAGGCCCTGCCAGGCTTCAGAAATCTGCACGTGGACGATCAGATGGCCGTGATCCAGTACTCCTGGATGGGCCTGATGGTGTTCGCCATGGGCTGGAGGAGCTTTACAAACGTGAACAGCCGGATGCTGTATTTCGCCCCTGACCTGGTGTTTAACGAGTACCGGATGCACAAGAGCCGGATGTATAGCCAGTGCGTGAGGATGCGCCACCTGAGCCAGGAGTTCGGCTGGCTGCAGATCACCCCTCAGGAGTTCCTGTGCATGAAGGCCCTGCTGCTGTTTTCCATCATCCCAGTGGACGGCCTGAAGAACCAGAAGTTCTTTGATGAGCTGAGGATGAATTACATCAAGGAGCTGGACAGGATCATCGCCTGCAAGCGCAAGAACCCCACCTCCTGTTCTAGGCGCTTTTATCAGCTGACAAAGCTGCTGGATAGCGTGCAGCCTATCGCAAGGGAGCTGCACCAGTTCACATTTGACCTGCTGATCAAGTCCCACATGGTGTCTGTGGATTTCCCCGAGATGATGGCCGAGATCATCAGCGTGCAGGTGCCAAAGATCCTGTCCGGCAAGGTGAAGCCCATCTACTTTCACACCCAGTGAGGATCC

#### Expression and purification of AR constructs

AR AD (1-558 aa) WT and mutants were recombinantly produced in *E. coli* Rosetta (DE3) cells transformed with pDEST17 plasmid encoding His-AR-AD, as described previously^75^. Briefly, cell cultures at OD_600_ 0.5 were induced with 0.1 mM IPTG at 22°C overnight. Cells were lysed in PBS buffer and centrifuged. The pellet was solubilized overnight in Tris buffer (20 mM Tris, 500 mM NaCl, 5 mM Imidazole, pH 8) containing 8 M Urea, 500 mM NaCl at pH 8. Protein was captured on Nickel columns (His Trap HP, GE Healthcare) and eluted with 500 mM Imidazole. After urea removal by dialysis, His-tag was cleaved by TEV protease protein at 4 °C overnight. Urea at 8 M was added to cleaved protein prior to reverse-nickel affinity chromatography to separate noncleaved protein and His-tag. Protein in the flowthrough was concentrated, filtered and stored at −80°C. To prevent protein aggregation or instability, an additional purification step was conducted running the sample on a Superdex 200 16/600 column pre-equilibrated with AR AD buffer (20 mM NaP, 1 mM TCEP pH 7.4). Tau-5* (330-448 aa) was expressed and purified as previously described^24^ and an equivalent protocol was used to express and purify fragment AR AD (441-558).

AR-LBD (663-919) containing an N-terminal His-tag and encoded in pET15b plasmid (Addgene #89083) was expressed in Rosetta (DE3) cells with 1 mM IPTG at 16 °C overnight. Cells were resuspended in Ni-Wash buffer (25 mM HEPES, 500 mM NaCl, 10% glycerol, 1 mM DTT, 10 μM DHT, 1% Tween-20, 20 mM Imidazole at pH 7.4), lysed and centrifuged. Soluble protein was captured by IMAC and eluted with 500 mM Imidazole. During an overnight dialysis, His-tag was cleaved by thrombin (GE Healthcare) and NaCl concentration was reduced to 100 mM. Cleaved protein was captured by cation exchange (GE Healthcare) and eluted with 1 M NaCl gradient. LBD was injected in a Superdex 200 16/600 column pre-equilibrated with 25 mM HEPES, 250 mM NaCl, 10% glycerol, 1 mM TCEP, 10 μM DHT, 1 mM EDTA, 0.5% Tween-20 at pH 7.2. MED1 IDR (948-1573), encoded in a peTEC plasmid, was kindly gifted by Prof. Thomas Graf. A 3C cleavage site was introduced by Q5 directed mutagenesis (NEB) between mCherry and MED1 sequence providing peTEC-His-mcherry-3C-MED1-IDR plasmid. Protein was expressed in B834 (DE3) cells at 16 °C overnight with 0.1 mM IPTG. Upon cell lysis in Tris buffer but 100 mM NaCl, soluble cell fraction was injected in a HisTrap HP column and protein was eluted with 500 mM Imidazole. Eluted protein was concentrated and separated by cation exchange chromatography. Fractions collected were cleaved by 3C protease and MED1-IDR was separated from other protein fragments by SEC chromatography with Superdex 200 16/600 column pre-equilibrated with 20 mM NaP, 100 mM NaCl, 1 mM TCEP at pH 7.4.

RNAPII CTD (1592-1970) was produced in *E. coli* B834(DE3) cells transformed with pDEST17 plasmid encoding H6-3C-RNAPII-CTD. Protein was expressed at 25 °C overnight with 0.1 mM IPTG and extracted from insoluble cell fraction. Pellet was resuspended in Tris Buffer with 8 M Urea and loaded on HisTrap HP column. Captured protein was dialyzed against 50 mM Tris-HCl, 50 mM NaCl, 1 M NaCl at pH 8 and cleaved by 3C protease overnight at 4 °C. RNAPII-CTD was injected in a Superdex 200 16/600 column pre-equilibrated with 20 mM NaP, 150 mM NaCl, 5 % glycerol, 1 mM TCEP at pH 7.4.

AR-LBD, MED1-IDR and RNAPII-CTD fractions from SEC were concentrated, filtered and stored at −80 °C for its further use.

#### Turbidity measurements

Protein samples were prepared in AR AD buffer (20 mM NaP, 1 mM TCEP pH 7.4) at the protein and NaCl concentration indicated on ice. Samples were centrifuged at 21130 rcf for 20 min at 4 °C and supernatant was transferred to a quartz cuvette. Phase separation cloud points of protein solutions were monitored by their absorbance at 340 nm as a function of temperature on a Cary 100 Multicell UV-vis spectrophotometer equipped with Peltier temperature controller at a rate of heat of 1 °C/min. The T_c_ were obtained as the maximum of the first order derivative of the obtained curves from 3 independent samples.

#### Protein labeling

For in vitro condensation experiments, proteins were labeled with fluorescent dye instead of tagged with fluorescent protein to avoid nonspecific interaction in heterotypic condensates. AR AD and MED1-IDR were fluorescently labeled with Dylight 405 and Alexa Fluor 647 respectively unless otherwise indicated in the figure captions. LBD and RNAPII-CTD were labeled with Oregon Green 488. In all cases, sulfhydryl-reactive dyes were used reacting to the naturally occurring cysteines of the protein except for RNAPII-CTD in which an N-terminal Cys was added. Protein was labeled according to the manufacturer’s instructions for sulfhydryl-reactive dyes(Thermo). Briefly, protein and dye were mixed at 1:20 ratio in each protein storage buffer adjusted at pH 7.5 overnight at 4 °C. 1 mM DTT was added to stop reaction and protein was separated from free dye with a pre-equilibrated PD-10 column. Protein was concentrated, filtered and concentration and conjugation efficiency were analyzed following the manufacturer’s instructions for sulfhydryl-reactive dyes (Thermo).

#### Fluorescence microscopy of *in vitro* protein condensation

Each protein solution was previously prepared by adding *ca*. 1 % of equivalent labeled protein and stored on ice. Samples were prepared by mixing proteins at the indicated protein concentration with AR AD buffer (20 mM NaP, 1 mM TCEP pH 7.4) in low binding PCR tubes at RT. Once all proteins were mixed the phase separation trigger was added; NaCl for AR samples or Ficoll 70 for transcriptional component samples. Samples were homogenized and 1.5 μl was transferred into sealed chambers containing samples composed by a slide and a PEGylated coverslip sandwiching 3M 300 LSE high-temperature double-sided tape (0.34 mm). Coverslips were PEGylated according to the published protocol^76^. Images were taken using Zeiss LSM 780 Confocal Microscope with a Plan-ApoChromat 63x/1.4 Oil objective lens.

#### NMR experiments

##### Assignment strategy

All NMR experiments were recorded at 5°C (278K) on either a Bruker 800 MHz (DRX or Avance NEO) or a Bruker Avance III 600 MHz spectrometer, both equipped with TCI cryoprobes. A 300 μM ^15^N,^13^C double labeled AR AD (441-558) sample in NMR buffer (20 mM sodium phosphate (pH 7.4), 1 mM TCEP, 0.05 %(w:w) NaN_3_) was used for backbone resonance assignment. The following series of 3D triple resonance experiments were acquired: HNCO, HN(CA)CO, HNCA, HN(CO)CA, CBCANH, and CBCA(CO)NH. Chemical shifts were deposited in BMRB (ID: 51476).

The assignment of AR AD (1-558) was guided by those of smaller AR fragments first studied here (residues 441-558) or previously reported (residues 1-151 (BMRB ID: 25607) and 142-448 (BMRB ID: 51479). In addition, 3D HNCO and HNCA experiments were acquired for two ^15^N,^13^C double labeled AR AD (1-558) samples (25 μM and 100 μM) dissolved in NMR buffer. For the 100 μM sample additional 3D HN(CA)CO and HN(CO)CACB experiments were also recorded. 3D experiments were acquired with 25% non-uniform sampling (NUS). Chemical shifts were deposited in BMRB (ID: 51480).

Backbone resonances of AR WT* were almost identical to those of AR AD (1-558), with only local differences in residues around the mutated position (L26), which were assigned using non-uniform sampled 3D BEST-TROSY HNCO and HNCA experiments^77^ recorded on a 50 μM ^15^N,^13^C double labeled WT* AR AD sample dissolved in NMR buffer.

##### Site-specific and residue-type identification of oligomerization

The oligomerization of AR AD was monitored by recording the induced intensity changes on the two-dimensional ^1^H,^15^N correlation spectrum by adding increasing amounts of unlabeled sample on a 25 μM ^15^N-labeled AR AD to reach a total concentration of 57.5, 100.8, 122.5 and 155 μM, respectively. Spectra was recorded using 128 scans per increment (experimental time of 21h per spectrum) to ensure the proper quantification of intensities in the regions with weaker signals.

##### Helicity studies upon TFE addition

The effect of TFE on 50 μM WT* AR AD an Tau-5* secondary structures were monitored by a series of ^1^H,^15^N correlation spectra and non-uniform sampled 3D BEST-TROSY, HNCO and HNCA experiments recorded in the presence of increasing TFE amounts (0, 2.5 and 5%).

##### Binding studies to AR-drugs

EPI-001 and 1aa binding to Tau-5* was studied by comparing ^15^N chemical shifts in 2D ^1^H,^15^N CP-HISQC^78^ spectra at 37°C (310K) of 60 μM Tau-5* in the absence and presence of 60 μM compounds (ratio 1:1). Samples contained NMR buffer (above) pH 6.6 with 200 mM NaCl and 2% DMSO-d6. The CP-HISQC pulse sequence and pH 6.6 were chosen to reduce water exchange of labile amide protons at 37°C (310K).

##### Data processing

Data processing was carried out with qMDD^79^ for non-uniform sampled data and NMRPipe^80^ for all uniformly collected experiments. Data analysis was performed with CcpNmr Analysis^81^. Helix populations were extracted using the δ2D software^71^.

#### Live-cell microscopy

PC3 cells were seeded in collagen I-coated µ-slide 4-well Glass Bottom plates (Ibidi 80426) at 60% confluency 24 hours before transfection. Then, 170 ng of expression vectors encoding androgen receptor (AR) tagged with eGFP (eGFP-AR) or mutants were transfected per well using polyethylenimine (PEI) (Polysciences) at a ratio of 1 µg DNA to 3 µl PEI. Four hours after transfection, media was changed to RPMI supplemented with 10% charcoal stripped FBS and cells were cultured for 16 hours before imaging. Transiently transfected PC3 cells expressing eGFP-AR were imaged in 3D for one minute every 15 seconds to acquire a baseline readout of AR expression. Cells were then immediately treated with 1 nM of DHT and imaged consecutively for 1 h (t_DHT_=1 h) every 15-sec time interval acquisition. Time lapse imaging was performed in an Andor Revolution Spinning Disk Confocal with an Olympus IX81 microscope and a Yokogawa CSU-XI scanner unit. Images were acquired with an Olympus PlanApo N 60x/1.42 Oil objective lens. A stable temperature (37°C) was maintained during imaging in a CO2 and temperature regulated incubation chamber (EMBL, Heidelberg, Germany). eGFP was excited with a 488 nm laser and Z-stack images were acquired every 0.45 μm step size. Time lapse images were compiled, processed and analyzed with Fiji image processing software (ImageJ)^82^. Intensity thresholds were set manually and uniformly to minimize background noise.

FLAG-MTID-AR-WT and PC3 FLAG-MTID-AR-WT-Y22toS cell lines were seeded in 24 well culture plates, on 12 mm sterilized coverslips. The next day ±50 μM biotin and 1nM DHT were added for 2 hours. The culture medium was removed and the cells were washed with PBS. Next, cells were fixed for 15 min with 4% paraformaldehyde. After fixation, cells were washed with PBS and then permeabilized with 0.1% Triton X-100 for 10 min. Coverslips were then washed and blocked with blocking buffer (3% BSA 0.1%Tween/PBS) for 1 hour at 37°C. Coverslips were incubated with primary antibody (1:100) - Anti-Androgen Receptor (Abcam), overnight. The next day coverslips were washed with PBS and secondary antibodies were added (1:500); Anti-Streptavidin antibody or Alexa Fluor™ 488 conjugate or Alexa Fluor 488 Goat anti-Rabbit IgG (H+L). Fluorescence images were acquired using a Leica TCS SP8 confocal microscope. Images were taken with 63x oil objectives, standard LAS-AF software.

HEK293T cells in DMEM 10% FBS were seeded at a density of 40,000 cells / well on glass bottom chambered coverslips (Ibidi 80827). 16 hours later, wells were refreshed with 280 µL seeding media and transfected with 50 nanograms of mEGFP expression plasmids shown in Fig. S2A using LipoD293 transfection reagent (SignaGen SL100668) according to the manufacturer’s protocol. 48 hours later, wells were refreshed with media spiked with 10 nM DHT or equivalent DMSO control (v/v). 4 hours after treatment, coverslips were imaged on a Zeiss LSM 880 Confocal Microscope with a Plan-ApoChromat 63x/1.4 Oil DIC objective lens in a CO_2_ incubation chamber set to 37°C. Images were acquired across two biological replicates.

#### STED microscopy

##### Sample preparation

HEK293T and HeLa eGFP-AR cells in DMEM 10% FBS were seeded at a density of 40,000 cells / well on glass bottom chambered coverslips (Ibidi 80827). 16 hours later, wells containing HEK293T cells were refreshed with 280 µl seeding media and transfected with 50 nanograms of mEGFP expression plasmids shown in Fig. 1B using LipoD293 transfection reagent (SignaGen SL100668) according to the manufacturer’s protocol. 48 hours later, wells were refreshed with media spiked with 10 nM DHT. Samples were imaged after 4 hours of DHT treatment.

LNCaP cells (Clone FGC, ATCC CRL-1740) were seeded in RPMI-1640 5% FBS onto PLL coated 18 mm #1.5 thickness glass coverslips (Sigma P4707, Roth LH23.1) at a density of 100,000 cells / coverslip in a 24 well plate. 16 hours later the media was refreshed and cells were incubated further for another 24 hours. For fixation, wells were washed with PBS, then fixed with 1 mL of 4% PFA in PBS for 20 minutes at room temperature. After a second wash in PBS, cells were permeabilized with 0.5% Triton-X, PBS (v/v) (Sigma 93443) and then stained with 1:50 AR primary antibody (AR 441, scbt 7305) and 1:200 STAR 635P secondary antibody (Abberior, ST635P-1001). Nuclear translocation of AR signal was validated by staining LNCaP cells grown in RPMI-1640 5% CSS (Gibco, A3382101) with the same protocol. DNA was counterstained with 1:2000 Spy555-DNA (spirochrome, SC201) and samples mounted onto glass slides with vectashield (Biozol, VEC-H-1900-10).

##### Live-Cell STED

HEK293T and HeLa cells were imaged on a Leica Stellaris STED DMI 8 microscope equipped with an okolab incubation chamber set to 37°C and 5% CO_2_ constant. EGFP imaging was performed using a 473 nm stimulation wavelength laser at 20% power and a 592 nm depletion laser at 20% power. Images were taken using a HC PL APO CS2 63x/1.40 oil objective with a final resolution of 23 nanometers / pixel.

##### STED FLIM

Fixed and stained LNCaP cells were imaged on a Leica Stellaris STED DMI 8 microscope. Abberior STAR 635P immuno-fluorescence imaging was performed using a 633 nm stimulation wavelength laser at 5% power and a 776 nm depletion laser at 5% power. Images were taken using a HC PL APO CS2 63x/1.40 oil objective with a final resolution of 48 nanometers / pixel. FLIM cutoffs and τ-STED deconvolution strengths were determined automatically using Leica LAS-X software v 2.5.6 to filter background photons with low fluorescence lifetimes. (Supplementary Fig. 1D).

#### FRAP assay in live cells

PC3 cells were transfected and prepared for microscopy in identical conditions to those of live cell imaging experiments. Before performing Fluorescence Recovery after Photobleaching (FRAP), cells were treated with 1 nM DHT. FRAP data for each condition were acquired over the course of approximately 1 hour after treatment, combining results for each condition as no trend was observed between FRAP data acquired at the beginning versus the end of the hour. FRAP measurements were performed on an Andor Revolution Spinning Disk Confocal microscope with a FRAPPA Photobleaching module and a iXon EMCCD Andor DU-897 camera. Images were taken at 100x/1.40 Oil U Plan S-Apo objective lens. Pre-bleaching and fluorescence recovery images of the eGFP-AR were acquired at the same 488 nm laser power with an exposure time of 100 msec. Bleaching was done in a 10×10 pixel square ROI on top of a droplet for 5 time repeats at maximum intensity 488 nm laser power at 40 usec dwell time bleaching. Twenty pre-bleached images and 200 post-bleached images were taken in total every 180 msec time interval acquisition. Post-bleached images were acquired immediately after the bleaching. Mean gray intensity measurements were quantified in three different Regions of Interest (ROIs) in each FRAP experiment: A bleached region, a background region outside the cells and a region spanning the whole cell were drawn to allow to normalize the gray values. Fiji software (ImageJ) was used to measure it in each ROI by using plot Z-axis profile function to extract the intensity data. Exported csv tables were normalized and fitted in EasyFrap software^83^ in order to extract kinetic parameters such as T-half and mobile fraction. Double normalization was used to correct for fluorescence bleaching during imaging and for intensity level differences.

#### Luciferase reporter assay in HEK293T

HEK293T cells were co-transfected with an ARE-luc construct containing a luciferase reporter gene under the control of three androgen response elements (ARE) (kindly provided by Maria Pennuto’s lab), along with an empty vector, an AR-expression vector (pEGFP-C1-AR or AR V7) or different mutants in the presence or absence of DHT. HEK293T cells were maintained in DMEM with 10% charcoal stripped FBS during the assay. Transfections were carried out using PEI and cells were treated with vehicle or 1 nM DHT 24 h after transfection. Cell extracts were prepared 48 h after transfection when eGFP-AR mutants are mostly nuclear and assayed for luciferase activity using the Promega luciferase detection kit. Luciferase activities were normalized to co-transfected β-galactosidase activity^84^.

#### Luciferase reporter assays in LNCaP

PSA(6.1kb)-Luciferase, 5xAP1-luciferase, V7BS_3_-luciferase and AR-V7 plasmids and transfections of cells have been previously described ^53,56,58,85^. PSA(6.1kb)-luciferase reporter plasmid (0.25 μg/well) was transiently transfected into LNCaP cells that were seeded in 24-well plates. Twenty-four hours after transfection, cells were pre-treated with compounds for 1 hour prior to the addition of 1 nM R1881 and incubation for an additional 24 hours. For the V7BS3-luciferase reporter, an expression vector encoding AR-V7 (0.5 μg/well) and a filler plasmid (pGL4.26, 0.45 μg/well) were transiently co-transfected with V7BS3-luciferase reporter plasmid (0.25ug/well) into LNCaP cells in 24-wells plates. After 24-hours, the cells were treated with the indicated compounds for an additional 1 hour. Transfections were completed under serum-free conditions using Fugene HD (Promega, Madison, Wisconsin). Luciferase activity was measured for 10 seconds using the Luciferase Assay System (Promega, Madison, WI) and normalized to total protein concentration determined by the Bradford assay. Validation of consistent levels of expression of AR-V7 protein was completed by Western blot analyses.

#### Proliferation assays

LNCaP cells (Clone FGC, ATCC CRL-1740) in RPMI 1640 5% FBS were seeded at a density of 4000 cells / well into 96 well flat bottom plates (Greiner, 655075) pre-coated with poly-L-lysine (Sigma P4707). 16 hours later triplicate wells were refreshed with 100 µL of seeding media spiked with 7x serial dilutions of EPI-001 from 200 µM (Selleckchem Lot #S795502), 7 x serial dilutions of 1ae from 50 µM, or DMSO control, at 0.5% DMSO (v/v) constant. 96 hours later, wells were washed with 200 µL PBS and then fixed with 100 µL of 4% PFA in PBS for 20 minutes at room temperature. Post fixation, LNCaP nuclei in each well were counterstained using 100 µL of Hoescht 33342 (abcam ab228551) diluted to 1:4000 in PBS for 20 minutes at room temperature. After a final wash in PBS, plates were imaged using a Celldiscoverer 7 microscope equipped with a 20x air objective. 25 tile regions (5 x 5 tiles) were imaged for each technical replicate well (triplicate wells for each dose and compound). Data was acquired across 2 biological replicates performed in separate weeks.

To compare the antiproliferative effects of 1ae and enzalutamide in LNCaP and LNCaP95 cells, LNCaP cells (5,000 cells/well) were plated in 96-wells plates in their respective media plus 1.5% dextran-coated charcoal (DCC) stripped serum. LNCaP cells were pretreated with the compounds for 1 hour before treating with 0.1 nM R1881 for an additional 3 days. Proliferation and viability were measured using Alamar blue cell viability assay following the manufacturer’s protocol (ThermoFisher Scientific, Carlsbad, California). LNCaP95 cells (6,000 cells/well) were seeded in 96-well plates in RPMI plus 1.5% DCC for 48hrs before the addition of compounds and incubation for an additional 48hrs. BrdU incorporation was measured using BrdU Elisa kit (Roche Diagnostic, Manheim, Germany).

#### Quantitative real time polymerase chain reaction (qRT-PCR)

LNCaP cells (Clone FGC, ATCC CRL-1740) in RPMI 1640 5% FBS were seeded at a density of 300,000 cells / well in 6 well plates. 16 hours later wells were refreshed with seeding media spiked with either EPI-001 or 1ae at doses roughly corresponding to IC_50_ and IC_10_ values calculated from proliferation assays, indicated in Supplementary Fig. 7A, and 0.5% v/v DMSO control.t. After 6 and 24 hours media was removed and cells were harvested using 300 µL of TRIzol reagent (Thermo 15596026) for each well. RNA was then extracted using a Zymo DirectZol Micro kit (Zymo R2062) according to the manufacturer’s protocol. cDNA was synthesized using 1 µg of RNA, random hexamer primers, and the RevertAid First Strand cDNA Synthesis kit (Thermo K1622). cDNA harvested from LNCaP cells treated at each compound, dosage, and time point were then probed for transcript targets on 384 well plates using the SYBR Green master mix (Thermo A25777), and a QuantStudio 7 real time qPCR machine. For calculation of fold change (2^-ΔΔCt^ method), Ct values from target regions were normalized to Ct values from control regions from the treatment sample, and then normalized to the DMSO sample. Data was acquired across 3 biological replicates performed on separate days. The following target primer sequences were used to probe transcript levels.

FKBP5 forward primer 1:

GCAACAGTAGAAATCCACCTG

FKBP5 reverse primer 1:

CTCCAGAGCTTTGTCAATTCC

FKBP5 forward primer 2:

AGGAGGGAAGAGTCCCAGTG

FKBP5 reverse primer 2:

TGGGAAGCTACTGGTTTTGC

PSA (*KLK3*) forward primer 1:

TGTGTGCTGGACGCTGGA

PSA (*KLK3*) reverse primer 1:

CACTGCCCCATGACGTGAT

PSA (*KLK3*) forward primer 2:

AGGCCTTCCCTGTACACCAA

PSA (*KLK3*) reverse primer 2:

GTCTTGGCCTGGTCATTTCC

β-Glucuronidase forward primer:

CTCATTTGGAATTTTGCCGATT

β-Glucuronidase reverse primer:

CCGAGTGAAGATCCCCTTTTTA

#### RNA-Sequencing data generation

LNCaP cells (Clone FGC, ATCC CRL-1740) in RPMI 1640 5% FBS were seeded at a density of 300,000 cells / well in 6 well plates. 16 hours later, wells were refreshed with seeding media spiked with either EPI-001 or 1ae at the doses indicated in Fig. 6C and 0.5% v/v DMSO control. After 6 and 24 hours, media was removed and cells were harvested using 300 µL of TRIzol reagent (Thermo 15596026) for each well. RNA was then extracted using a Zymo DirectZol Micro kit (Zymo R2062) according to the manufacturer’s protocol. Total RNA-seq libraries were then prepared using 1 µg of RNA from each sample and the KAPA RNA HyperPrep Kit with RiboErase (Roche KR1351) according to the manufacturer’s protocol with 10 amplification cycles. Libraries were sequenced on a NovaSeq 6000 with paired-end reads of 100 base pairs, with a read depth of 50 million fragments / library. Three libraries from three corresponding biological replicates were prepared for each treatment (time, dosage, and compound).

#### Western Blot

Cells were washed and harvested in PBS 1x, lysed in RIPA buffer 1x (Thermo, 88900) containing phosphatase and protease inhibitors (Roche). Lysates were centrifuged at 15,000 g to separate soluble and pellet fractions. Total protein was quantified using BCA assay (Pierce Biotechnology). Proteins were resolved by 7.5 or 15% SDS-PAGE, transferred to PVDF membranes and blocked with 5% non-fat milk in TBST for 1 hour at room temperature with shaking. The membranes were incubated with the following antibodies: anti-GAPDH (#39-8600, 1:2000),anti-androgen receptor (ab108341, 1:1000) and anti-Streptavidin (#926-32230, 1:1000). Anti-mouse HRP-conjugated (G-21040, 1:10000) and anti-rabbit HRP-conjugated (65-6120, 1:10000) secondary antibodies from Invitrogen.

#### Lentiviral production for FLAG-BioID-AR Cell Lines

FLAG-MTID, FLAG-AR-WT-MTID or FLAG-22YtoS-MTID were subcloned from pcDNA3.1(-) (Genscript) into pLenti-CMV-MCS-GFP-SV-puro (addgene #73582) by replacing GFP using XbaI-BamHI digestion. Vectors were co-transfected with lentiviral packaging plasmid vectors REV (Cat# 12253), RRE (Cat# 12251) and VSV-G (Cat# 8454) into 293T cells with PEI (Sigma-Aldrich). Two days after transfection, virus-containing medium was collected and filtered through 0.45-µm a low-protein-binding filtration cartridge. The virus containing media was directly used to infect LNCaP/PC3 cells in the presence of polybrene (8 µg/mL) for 48 hours, before 1 μg/ml puromycin was introduced for 72 hours to select for stable cell lines. Cell lines were maintained in conditions previously described for Prostate Cancer cells. pMDLg/pRRE was a gift from Didier Trono (Addgene plasmid # 12251; http://n2t.net/addgene:12251; RRID:Addgene_12251). pCMV-VSV-G was a gift from Bob Weinberg (Addgene plasmid # 8454; http://n2t.net/addgene:8454; RRID:Addgene_8454). pRSV-Rev was a gift from Didier Trono (Addgene plasmid # 12253 ; http://n2t.net/addgene:12253; RRID:Addgene_12253).

#### BioID-MS

Prior to BioID experiments MTID containing stable cell lines were generated by lentiviral infection and Puromycin selection. They were subsequently grown in RPMI 1640 Medium modified w/L-Glutamine w/o Phenol Red or Biotin (US Biological life sciences, R9002-01) with 10% (v/v) charcoal stripped FBS for 48 hours. Cells were seeded and the next day ±50 μM biotin (IBA GmbH; 2-1016-002) and 1 nM DHT were added for 2 hours. For small molecule inhibitors cells were pre-treated with 1 hour of either EPI-001 or 1ae, then additionally with 2 hours of DHT + Biotin. For mass spectrometry, cells were harvested with trypsinization, washing two times in PBS and snap freezing on dry ice. Cell pellets were lysed in modified RIPA buffer (1% TX-100, 50 mM Tris-HCl, pH 7.5, 150 mM NaCl, 1 mM EDTA, 1 mM EGTA, 0.1% SDS, 0.5% sodium deoxycholate and protease inhibitors) on ice, treated with 250 U benzonase (Millipore) and biotinylated proteins were isolated using streptavidin-sepharose beads (GE Healthcare). Proteins were washed in ammonium bicarbonate and digested with trypsin. Mass spectrometry was performed in the IRB Barcelona Mass Spectrometry and Proteomics facility as described previously^86^. Data was analyzed using SAINTq^87^.

#### Proximity Ligation Assay

Protein—protein interactions were studied using a Duolink In Situ Orange Starter Kit Mouse/Rabbit (Sigma, DUO92102) following the manufacturer’s protocol. Briefly, transduced Prostate cancer cells were seeded in coverslips and cultured overnight. The next day ± 50 μM biotin and 1 nM DHT were added for 2 hours or pre-treated with small molecule inhibitors initially. Slides were washed with cold 1 × PBS and fixed in 4% paraformaldehyde for 15 min, washed in PBS and permeabilized using 0.1% Triton X-100 for 10 min and washed then blocked with blocking buffer (3% BSA 0.1% Tween/PBS) for 1 hour at 37°C. The coverslips were blocked with Duolink Blocking Solution in a pre-heated humidified chamber for 30 min at 37°C. Primary antibodies were added and incubated overnight at 4°C. Then coverslips were washed with 1×Wash Buffer A and subsequently incubated with the two PLA probes (1:5 diluted in antibody diluents) for 1 h, then the Ligation-Ligase solution for 30 min, and the Amplification-Polymerase solution for 100 min in a pre-heated humidified chamber at 37°C. Before imaging, slides were washed with 1 × Wash Buffer B and mounted with a cover slip using Duolink In Situ Mounting Medium with DAPI. Fluorescence images were acquired using a Leica TCS SP8 confocal microscope. Images were taken with 100x oil objectives, standard LAS-AF software.

### Quantification and Statistical Analysis

#### Statistical Analysis

Pairwise comparisons shown in Fig. 1H, 2D, 2G, 3E, 5D, 5G, 5I, 6J, 6K and Supplementary Figs. 4D, G, H, J, K, 5D, 6C were performed with Student’s t-test or Mann-Whitney U test, as indicated in the figure legends, in base R or python. Differences were considered significant when adjusted p-values were less than 0.0001 (****), 0.001 (***), 0.01 (**), or 0.05 (*).

#### AR ΔNLS image analysis in live-cells

A custom-made macro in Fiji software was developed to quantify the total number and the size of AR condensates into the cytoplasm as a function of time (Fig. 3E and Supplementary Figs. 4D, G, J, K). This macro also quantifies the total area of the cytosol to normalize the results.

The macro creates z intensity projections of the 3D stacks acquired. A manual step of drawing a ROI was integrated into the macro to select the nuclei to be removed and only to keep the cytoplasm area for the detection and quantification of the AR condensates. After filtering and thresholding, the cytosol area was segmented and quantified. Then a mathematical operation was done between the resulting mask of the cytosol, now without the nuclei, and the z maximum intensity projection data to detect and quantify the total number and the area of AR condensates in the cytosol. The quantification is done in 3 different time points after DHT exposure.

#### AR nuclear translocation rate analysis

A custom-made macro in Fiji software was developed to quantify the mean gray intensity value in the nuclei area along the time (Fig. 2B). The macro creates a z sum projection of the 3D stacks of the timelapse to be quantitative in the results. A stackreg plugin is used in the macro to register and correct the xy movement of the cells along the time; a manual step of drawing the nuclei area and the cytoplasm area is done, to extract automatically the mean gray values of these ROIs along the time.

#### Luciferase reporter assay in HEK293T

For the transcriptional activity assay, reported in Fig. 2G, a general linear model was used to compare differences in log transformed ARE-Luc vs β-galactosidase ratio between groups of interest using biological replicates as covariates. For clarity of representation, ARE-Luc vs β-galactosidase ratios are shown in the original scale.

#### Analysis of FRAP data for cell experiments

Mean intensities of bleached areas were corrected both for bleaching due to imaging over time and background noise. The corresponding calculations were performed with the EasyFrap by calculating the fluorescence intensity over time I(t). Obtained values were further normalized to the initial fluorescence by dividing I(t) by the mean gray value of the initial pre-bleaching acquisition images.

#### Granularity analysis

Image analysis was assisted by a custom ImageJ macro written at ADMCF. An individual segmentation mask was obtained for each nucleus (excluding the nucleoli) by simple median filtering, background subtraction and local thresholding. Nuclei exhibiting an insufficient or too strong level of expression were excluded manually and the standard deviation of the intensity was estimated inside the remaining nuclei in the original images. For the granularity analysis, reported in Fig. 2D, the standard deviations were compared across groups by linear regression. The relationship between standard deviation and mean intensity was also compared across groups, and reported in Supplementary Fig. 3A, by fitting a linear model with the standard deviation as response variable and taking the mean intensity, the group, the interaction between the group and the mean intensity and the biological replicate as explanatory variables. The slope between mean intensity and standard deviation was compared for every experimental group against the control through the interaction term of the linear model. Dunnett’s multiple comparisons correction was applied for comparing the linear effects of several experimental groups with a common control. Images of HEK239T cells transfected with mEGFP plasmids described in Supplementary Fig. 2A were analyzed using ZEN Blue version 3.2. Image fields were segmented for nuclear regions using automatic thresholding (Otsu thresholding) on the mEGFP channel, and the resulting objects were analyzed for mean intensity and standard deviation of pixels. As above, nuclear clustering (or granularity) was assayed as the standard deviation of pixels, and nuclear GFP concentration as the mean intensity of pixels in the corresponding nuclear object. Measurements were exported for data wrangling in R to create the plots shown in Supplementary Fig. 2B. 8 - 10 image fields were used to assay nuclei from each condition (transfection and treatment).

#### LNCaP dose response curves

Raw LNCaP nuclei counts from proliferation experiments, assayed as objects detected by automatic otsu thresholding on Hoechst signal from image fields from each well (aggregate of 25 tile regions), were used to construct dose response curves for EPI-001 and 1ae (Fig. 6C). Segmentation was performed using ZEN Blue version 3.2 on image data acquired across 2 biological replicates. Nuclear counts from each well were exported and processed using the DRC package in R^88^ to create dose response curves shown in Fig. 6C. Data was modeled with a three-parameter log-logistic function (lower limit 0) and the resulting fit was used to calculate IC_50_ and IC_10_ values for EPI-001 and 1ae (Fig. 6C).

#### In vitro droplet image analysis

For *in vitro* droplet analysis of AR AD in multiple component images in Fig. 1, droplets were identified applying a threshold (3, 255) to the channel sum image using FIJI. AR-AD intensity within the identified droplets larger than 0.1 μm across 3 image fields was extracted and plotted in Fig. 1H. Graph from Fig. 1J was obtained by normalizing each channel’s intensity from the plot profile of a section of a representative droplet using FIJI. Droplets from Fig. 3G were identified applying a threshold (3, 255) to the channel sum image using FIJI. Droplet size and density (number per field area) across 3 image fields was plotted in Supplementary Fig. 4H.

#### τ-STED image analysis

Composites acquired in τ-STED mode (Supplementary Fig. 1B) were exported as .tiff image fields using Leica LASX version 2.5.6 and analyzed using a custom FIJI pipeline (Supplementary Fig. 1C) available at https://github.com/BasuShaon/AR/tree/master/STED. In brief, the DNA counterstain was first used to identify and threshold nuclear objects. Clusters within nuclear objects were then detected using the rolling ball algorithm, with the size of the rolling ball set to 8 x the limit of detection (48 nanometers), according to standard protocol^89^. This enabled detection of nuclear AR clusters for cells imaged with the same τ-deconvolution strength. Nuclear AR clusters were pooled from 7 LNCaP nuclei, and analyzed for mean intensity and size as indicated in Supplementary Fig. 1B.

#### Chrom logD determination

Chrom logD values were experimentally determined as a measure of hydrophobicity of the 1aa family of compounds. The experimental evaluation was subcontracted to Fidelta, Ltd. Values of chrom logD were calculated from the equation:

Chrom logD = 0,0857*CHI - 2,

In which CHI is a chromatographic hydrophobicity index. CHI values were determined from gradient retention times at pH = 7.4. Chromatograms were measured by the Agilent 1100 HPLC instrument, using a Luna C18 analytical column from Phenomenex. 50 mM ammonium acetate (in H_2_O) and ACN were used as mobile phases, containing 0.1 % TFA. A method was optimized to 5 minute run with linear gradient from 0 to 100 % of ACN in the first 3 min.

#### Molecular dynamics (MD) simulation

A molecular dynamics simulation of the AR Tau-5 _R2_R3_ region (residues L391-G446, capped with ACE and NH2 groups) in the presence of 1aa was performed as a described previously^57^ and compared to previously reported simulation results of Tau-5_R2_R3_ in the presence of EPI-002 (Zhu et al). Briefly, an explicit solvent simulation was performed in a cubic box with a length of 7.5 nm and neutralized with a salt concentration of 20 mM NaCl by 8 Na^+^ ions and 5 Cl^−^ ions. The AR Tau-5_R2−R3_ protein was parameterized using the a99SB-*disp* force field; water molecules were parameterized with the a99SB-disp water model^90^. 1aa was parameterized using the GAFF ^91^ for ligand forcefield parameters. The replica exchange with solute tempering (REST2) algorithm^92^ was utilized to enhance conformational sampling. 16 replicas were run in parallel using a temperature ladder ranging from 300-500 K, with all protein atoms selected as the solute region. Tau-5_R2−R3_ with 1aa was simulated for 5.2 μs per replica respectively, for a total simulation time of 83.2 μs. Convergence of simulated properties was assessed by a comparison of the conformational sampling of each simulated replica as previously reported^57^, and statistical errors were calculated using a blocking analysis following Flyvbjerg and Peterson^93^. We define an intermolecular contact between a ligand and a protein residue as occurring in any frame where at least one heavy (non-hydrogen) atom of that residue is found within 6.0Å of a ligand heavy atom. To calculate a simulated K_D_ value for each compound, we define the bound population (P_b_) of each ligand as the fraction of frames with at least one intermolecular contact between a ligand and Tau-5_R2_R3_.

#### RNA-sequencing data pre-processing

Paired end sequencing reads were first quality checked using FASTQC and then aligned to the *Homo sapien* genome build hg19 using STAR aligner v2.7.5a^94^ with standard settings. 1^st^ and 4^th^ columns in ReadsPerGene.out.tab STAR output files (GeneIDs and reverse strand reads) were used to build raw count matrices for each sample library.

#### Differential expression analysis

Differential expression analysis between treatment conditions was conducted using the DESeq2 R/bioconductor package, a statistical tool that uses shrinkage estimates to compute fold changes^95^. First, raw count matrices from sample libraries were merged into a single object using the ‘DESeqDataSetFromHTSeqCount’ function with the design set to the treatment condition (time, compound, and dosage). The merged count matrix was then fit to the DESeq statistical model using the ‘DESeq’ function. The fit and merged matrix was then reduced using a variance stabilizing transformation ‘vst’ to visualize principal components 1 & 2 (Supplementary Fig. 7B). The fold change values in gene expression and corresponding significance scores were then extracted using the ‘results’ function by querying a contrast between any two conditions (Table S3). |Log_2_FC| > 1 and p-value < 1e-10 cutoffs were used to call differentially expressed genes in a given contrast (Fig. 6D).

#### Gene set enrichment analysis

Gene set enrichment analysis was performed using R/bioconductor packages fgsea and DOSE ^96,97^. Ranked gene lists were first constructed using log_2_FC values for genes in any given DESeq2 contrast by sorting log_2_FC values in descending order and filtering out duplicate entries. Ranked lists were then analyzed for the enrichment of 50 hallmark gene sets (collection H) obtained from the molecular signature database msigDB maintained by the Broad Institute using the ‘plotEnrichment’ and ‘plotfgseaRes’ functions in fgsea and the ‘GSEA’ function in DOSE (nperm = 10,000, p-value cutoff < 0.05).

Besides the commonly used gene set enrichment plot for a queried gene pathway (Supplementary Figs. 7C,E) we also represent enrichment scores for the top 10 negatively and top 10 positively enriched pathways as a dotplot with gradient scaling to the normalized enrichment score (red = positive NES, blue = negative NES) and size proportional to the statistical significance (p_adj_) of the calculated enrichment (Fig. 6E).

#### Mean expression value of genes in hallmark gene sets

Line plots for mean expression values of genes were adapted from^98^. In brief, reads from the merged count matrix were normalized according to the following equation log_2_(normalized DESeq counts + 1) to create a log_2_ normalized count matrix (Table S4). Normalized counts for each gene in the matrix were then z-score scaled using the ‘scale_rows’ function from the pheatmap R package. Code integrated with DESeq2 available at https://github.com/BasuShaon/AR/tree/master/RNAseqLoven. Values of the genes from the below gene sets were then plotted as indicated in Fig. 6F and Supplementary Fig. 7D as a function of compound concentration of EPI-001 and 1ae.

##### msigDB hallmark pathway set H

http://www.gsea-msigdb.org/gsea/msigdb/genesets.jsp?collection=H

*EPI-001 negative DEGs (24 hours of 25 µM EPI-001 vs 24 hours of DMSO, LNCaP)*

*KLK3, ADAM7, TBX15, FKBP5, PGC, LAMA1, ELL2, CHRNA2, STEAP4, DSC1, UGT2B28, TNS3, BMPR1B, SLC38A4, EAF2, TTN, SLC15A2, CCDC141, HPGD, TMEM100, MAF, F5, TRGC1*

## Supplementary Figure Legends

**Supplementary Figure 1.**
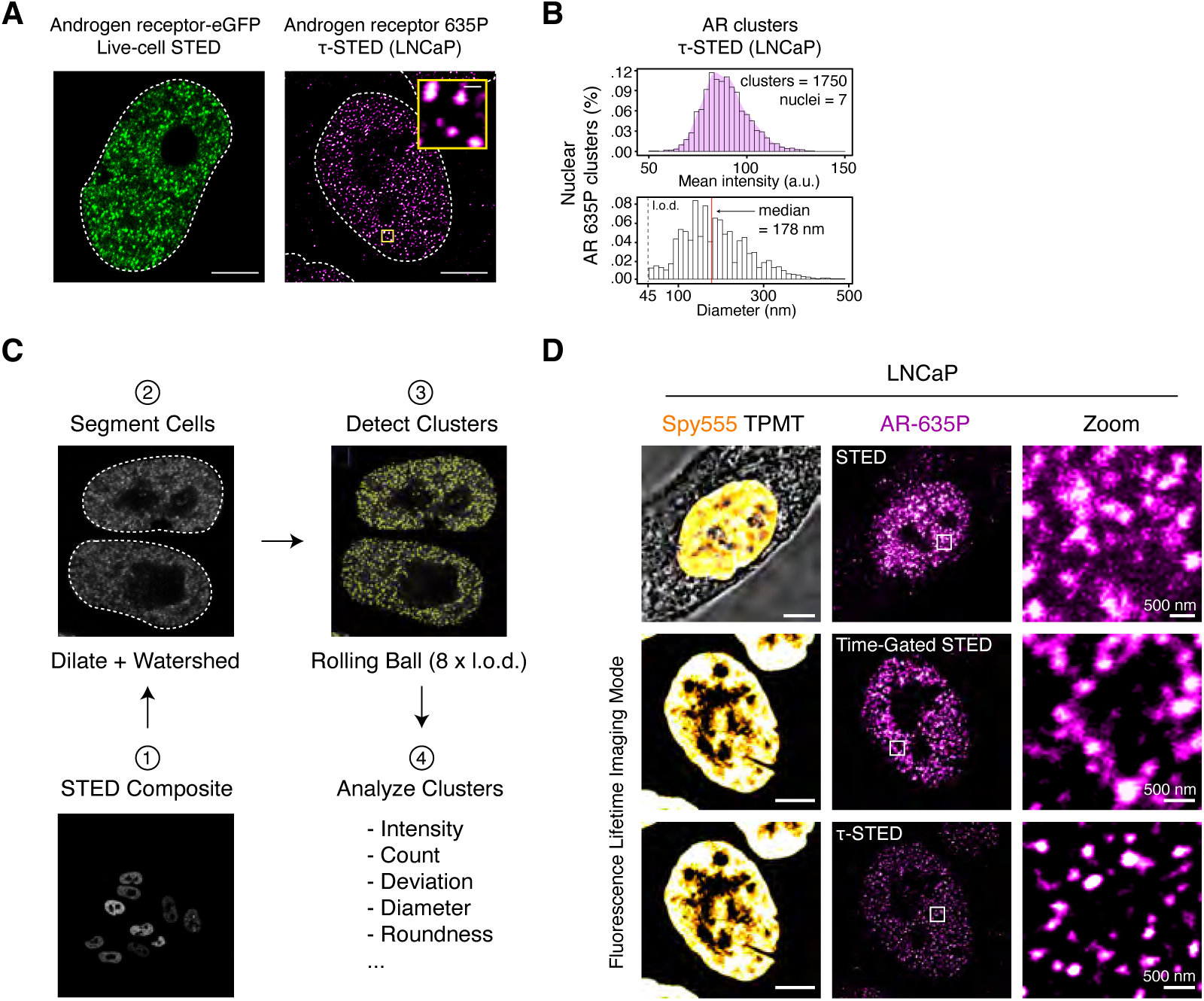
Characterization of AR condensates in cells using high resolution microscopy. **A)** (left) Live-cell stimulated emission depletion (STED) imaging of a HeLa cell nucleus expressing AR-eGFP, treated with 1 nM DHT for 4 hours. (right) τ-STED imaging of endogenous AR in fixed human prostate adenocarcinoma (LNCaP) cells. Large scale bars: 5 μm. Scale bar in τ-STED inset: 300 nm. Dashed line indicates the nuclear periphery. B) (top) Quantification of τ-STED intensity signal and (bottom) diameter of endogenous AR clusters in LNCaP cells (1750 AR clusters detected across 7 LNCaP nuclei imaged with same fluorescence time gating). L.o.d indicates the limit of detection. Density_max_ diameter (bin with highest density of AR clusters in the distribution of all detected AR clusters): 123 nm, median diameter: 178 nm. **C)** Quantification pipeline used to analyze STED image composites, showing segmentation of cells and detection of clusters using rolling ball background subtraction adjusted to 8 x the resolving capacity of the image (48 nanometers / pixel for TauSTED imaging of LNCaP cells). **D)** STED (top row) and FLIM STED images showing AR clusters in LNCaP nuclei before and after τ**-**STED deconvolution (middle and bottom row). Left column shows LNCaP nuclear counterstain using Spy555-DNA stain. Scale bar: 5 μm. Right panels show zoom-ins corresponding to intra-nuclear regions indicated by white boxes on panels in the central column. Scale bar: 500 nm.

**Supplementary Figure 2.**
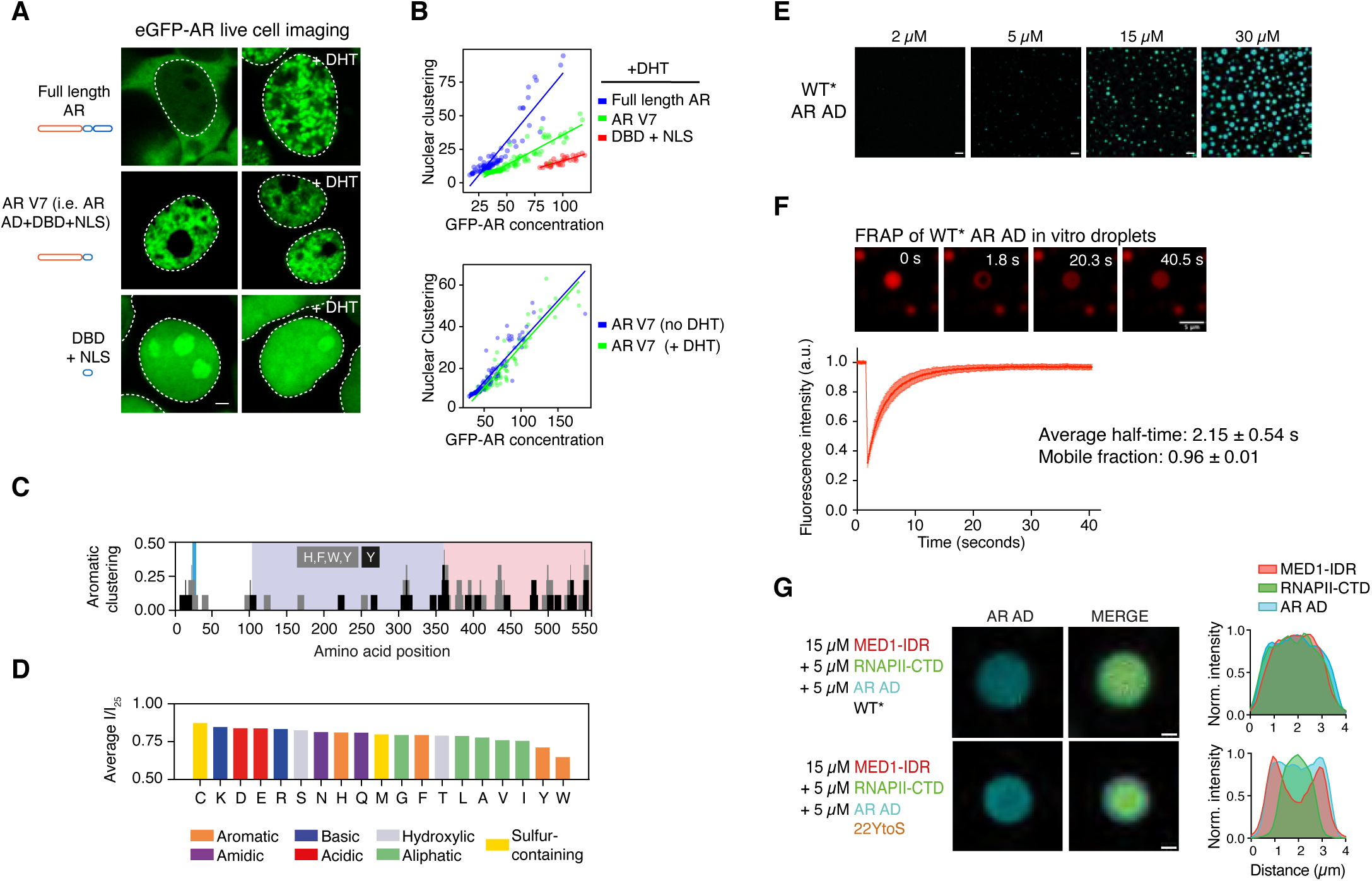
Tyrosine residues in AR-AD are the main drivers of AR phase separation. **A)** Live-cell confocal imaging of the indicated mEGFP construct transfected into HEK293T after treatment with vehicle or 10 nM DHT for four hours. Scale bar: 3 µm. Dashed lines indicate nuclear periphery. **B)** Quantification of confocal data in Supplementary Fig. 2A. Y-axis indicates the standard deviation, and x-axis indicates the mean intensity of pixels in the corresponding nucleus. Each dot represents measurements from an individual cell, and lines represent standard regression fits to the corresponding data spread (N = 2). **C)** Distribution of aromatic (Histidine, Phenylalanine, Tryptophan, Tyrosine) and Tyrosine residues along the AR AD sequence, clustered using a 9 amino acid window, where the shaded areas correspond to those represented in Fig. 1C. **D)** Average intensity ratio of the NMR resonances of the AR AD at the tested protein concentrations (57.5, 100.8, 122.5 and 155.0 μM) relative to their intensity at 25 μM grouped by amino acid type. **E)** Fluorescence microscopy images of in vitro AR AD (WT*) concentration-dependent condensation obtained in AR AD buffer (20 mM NaP, 1 mM TCEP pH 7.4) with 150 mM NaCl and 10% ficoll, where *ca* 1 % of AR-AD molecules were labeled with the dye Dylight 405. Scale bar: 10 µm. **F)** AR AD WT* liquid character *in vitro* by FRAP. Top panel: confocal microscopy images of WT* AR AD droplets labeled with Alexa-647, in 150 mM NaCl and 10 % ficoll before and after photobleaching in FRAP experiment. Scale bar 5 µm. Lower panel: average relative fluorescence intensity curve of WT* AR AD droplets as a function of time following photobleaching. Error bars represent s.d. of n=10 droplets. **G)** (Left) microcopy images of in vitro droplets formed by the indicated proteins. The signal of the AR AD channel and merged channel are shown. AR AD proteins were used in five-fold higher concentrations than in Fig. 1I. Scale bar: 1 µm. (Right) the representative droplet’s cross section intensity profile.

**Supplementary Figure 3.**
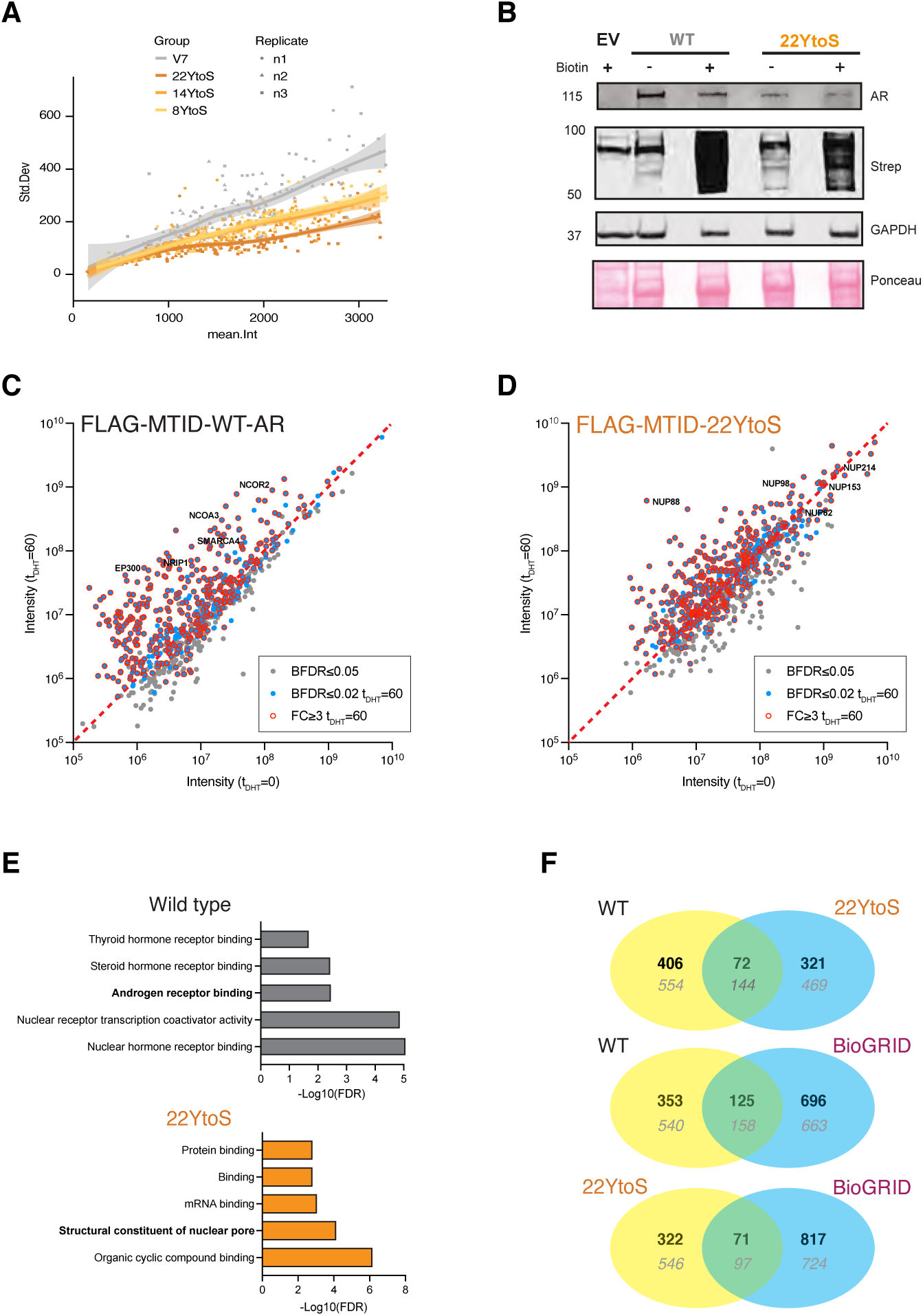
Tyrosine mutants decrease granularity, nuclear translocation and alter the proximal interactome of AR. **A)** Quantification of the nuclear granularity (standard deviation) as a function of the mean intensity of nuclei expressing the indicated AR-V7 constructs. **B)** Western blot showing expression of FLAG-MTID-AR or FLAG-MTID-Y22toS proteins in PC3 cells with antibodies for AR. Biotin-dependent labeling is shown with Streptavidin antibodies (Strep) and GAPDH and Ponceau staining are shown as loading controls. EV indicates the empty vector expressing FLAG-MTID. **C)** Scatter plot of the protein intensities at t_DHT_ = 0 and 60 for PC3 cells expressing FLAG-MTID-WT-AR following SAINTq analysis with control samples. Proteins with a BFDR ≤ 0.05 in either sample are shown (gray circle) and those with a BFDR ≤ 0.02 (blue circles) and/or FC ≥ 3 (red outline) in t_DHT_ = 60 are noted. Proteins in the networks shown in Fig. 2E are labeled. **D)** Same as in panel “C” for the FLAG-MTID-22YtoS samples. **E)** Enriched gene ontology molecular function (GO-MF) categories in the FLAG-MTID-WT-AR t_DHT_ = 60 samples (upper panel) and FLAG-MTID-22YtoS (lower panel). The top 75 most abundant proteins identified with a cutoff of BFDR ≤ 0.02 and FC ≥ 3 were analyzed using STRING and GO categories exported. The -log_10_(FDR) for selected GO-MF categories are graphed and the categories depicted in the networks in Fig. 2E are in bold. Full datasets and GO analysis results are provided in Table S1. **F)** Venn diagrams depicting overlapping proteins identified in the WT and 22YtoS samples (top), the WT and AR interactions reported in BioGRID (middle) and Y22toS and AR interactions reported in BioGRID (bottom). Numbers of proteins identified in each sample (t_DHT_ = 0 and 60) with a BFDR ≤ 0.02 and a FC ≥ 3 are indicated in bold and numbers of proteins identified with a BFDR ≤ 0.05 are indicated in gray. All proteins identified are provided in Table S1.

**Supplementary Figure 4.**
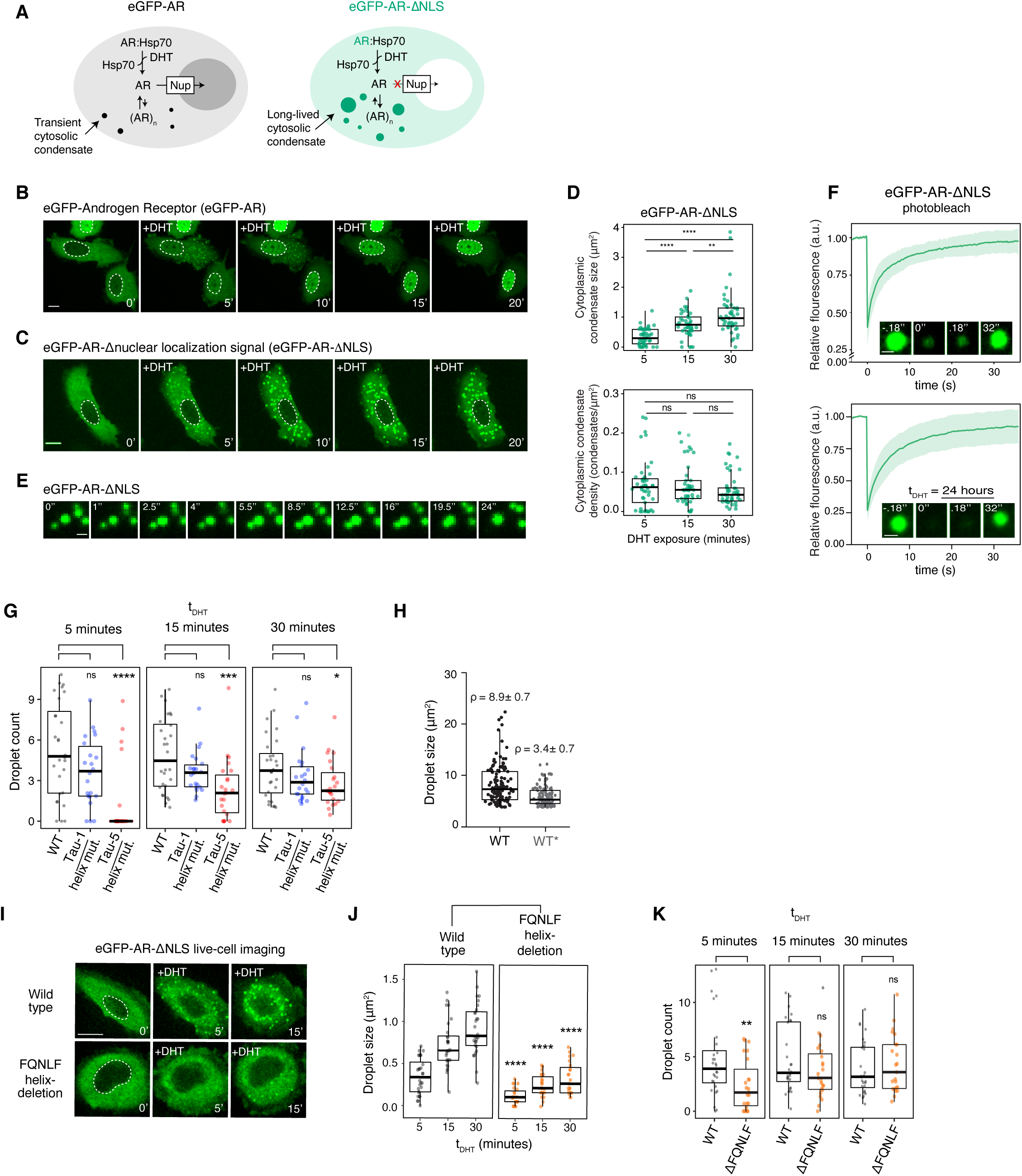
Transactivating units and motifs with helical propensity in AR AD contribute to condensation of AR *in vitro* and in cells. **A**) Schematic model describing the nuclear translocation pathway of eGFP-AR and cytoplasmic retention of eGFP-AR-ΔNLS upon exposure to ligand (DHT). **B,C)** Time-lapse fluorescence microscopy of eGFP-AR (**A**) and eGFP-AR-ΔNLS (**B**) condensates upon treatment with 1 nM dihydrotestosterone (DHT) in transiently transfected PC3 cells. Scale bar: 10 µm. Dashed line indicates the nuclear periphery. **D)** Distributions of average condensate size and density. Each dot corresponds to the mean values measured in an individual cell (n = 45 cells). P-values are from Mann-Whitney U tests. n.s.: not significant. **E)** Snapshots at the indicated time points highlighting a fusion event of eGFP-AR-ΔNLS condensates in the cytoplasm of a PC3 cell. Scale bar: 1 µm. **F)** Fluorescence recovery after photobleaching (FRAP) analysis of cytoplasmic eGFP-AR-ΔNLS condensates in PC3 cells 1 hour and 24 hours after addition of 1 nM DHT (t_DHT_ ≈ 1h). Average relative fluorescence intensity curve of the eGFP-AR-ΔNLS cytoplasmic condensates as a function of time is shown. Error bars represent s.d. of n = 34 condensates per time point. Within the box, representative images of condensates before and after photobleaching are shown. Scale bar: 1 µm. **G)** Effect of the mutations introduced in Tau-1 and Tau-5 on the density of the cytosolic condensates formed by eGFP-AR-ΔNLS as a function of t_DHT_ in PC-3 cells. Each dot corresponds to a cell (n > 20 cells). P values are from a Mann-Whitney U test. **H)** Quantification of droplet size and number from Fig. 3G. Droplets size are plotted with a p value < 0.0001 and the mean number of droplets per image field is indicated as density (droplets / 10^3^ um^2^). **I)** Effect of deleting the region of sequence of the AD containing the ^23^FQNLQ^27^ motif on the cytosolic condensates formed by eGFP-AR-ΔNLS upon addition of DHT. Scale bar:10 μm. Dashed line indicates nuclear periphery. **J, K)** Effect of deleting the region of sequence of the AD containing the ^23^FQNLQ^27^ motif on the average size (**D**) and density (**E**) of the cytosolic condenstates formed by eGFP-AR-ΔNLS as a function of t_DHT_. Each dot corresponds to a cell (n > 20 cells). P-values are from Mann-Whitney U tests.

**Supplementary Figure 5.**
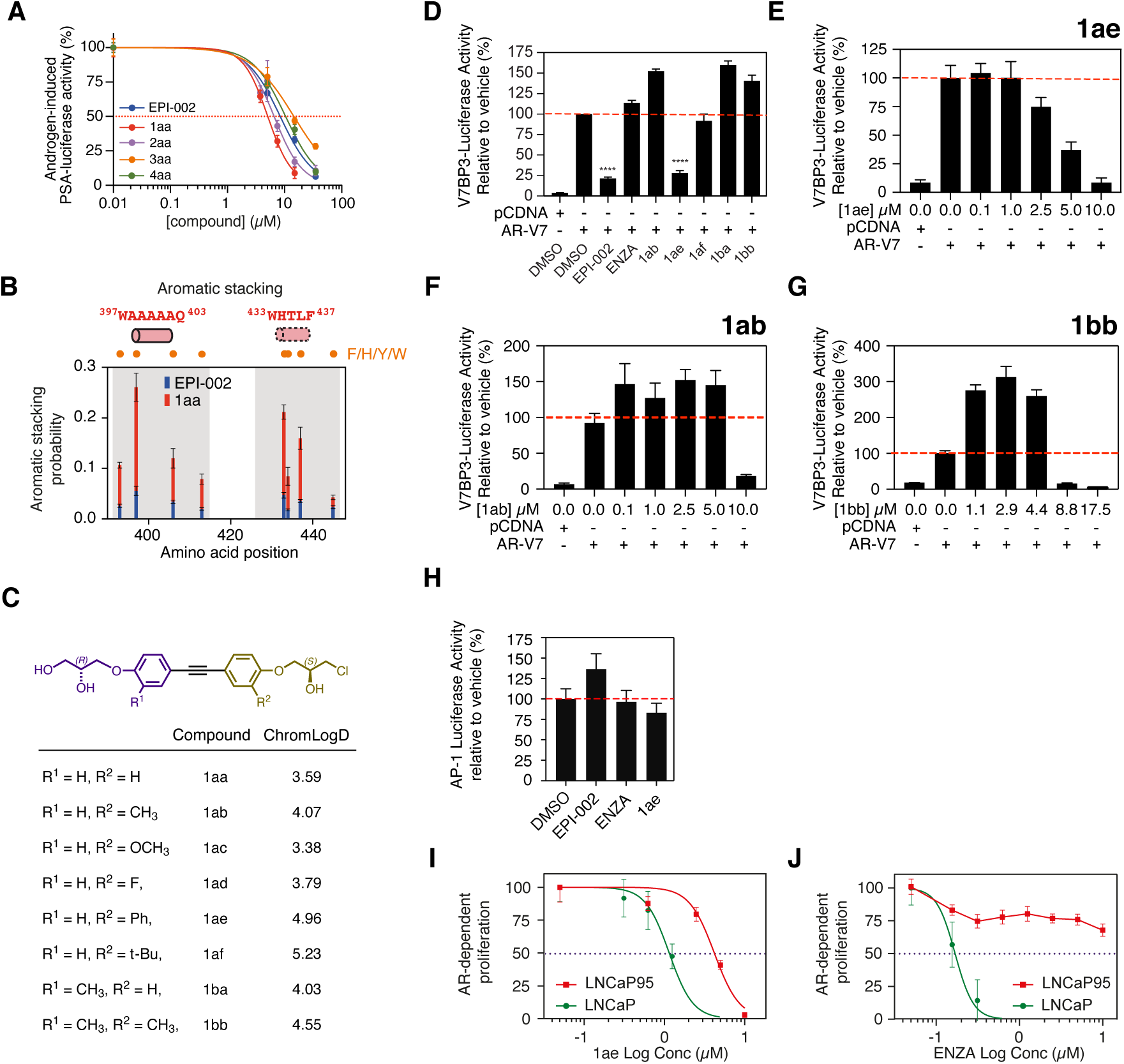
Characterisation of small molecules with enhanced potency. **A)** Populations of aromatic stacking contacts between aromatic side chains of Tau5_R2_R3_ and aromatic rings of EPI-002 (blue) and 1aa (red) with an indication of the positions of helical motifs and aromatic residues. The data was obtained from the 300 K REST2 MD simulations. **B)** Inhibition of the androgen-induced full-length AR transcriptional activity by compounds shown in Fig. 4A (N = 3). **C)** ChromLogD values of compounds developed from 1aa scaffold reporting their hydrophobicity (N=3). **D)** Comparison of EPI-002 (35 µM) and enzalutamide (ENZA, 5 µM) with the most potent compounds (5 µM) to block AR-V7 transcriptional activity (N = 3). **E, F, G)** Dose-dependent inhibition of AR-V7 transcriptional activity with 1ae (**G**), but not for 1ab (**H**) and 1bb (**I**). LNCaP cells that ectopically expressed AR-V7 were co-transfected with a V7BS3-luciferase reporter gene construct and incubated with the indicated concentrations of the compounds (N = 3). **H)** Activity of AP-1-luciferase reporter after incubating LNCaP cells with EPI-002 (35 µM), ENZA (5 µM), and 1ae (5 µM) or vehicle (DMSO) for 24 h (N = 3). **I)** 1ae blocked the proliferation of both LNCaP cells in response to androgen and AR-V-driven proliferation of LNCaP95 cells (N = 3). **J)** Enzalutamide (ENZA) blocked androgen-induced proliferation driven by full-length AR in LNCaP cells but had poor potency against AR-V-driven proliferation of LNCaP95 (LN95) cells (N = 3).

**Supplementary Figure 6.**
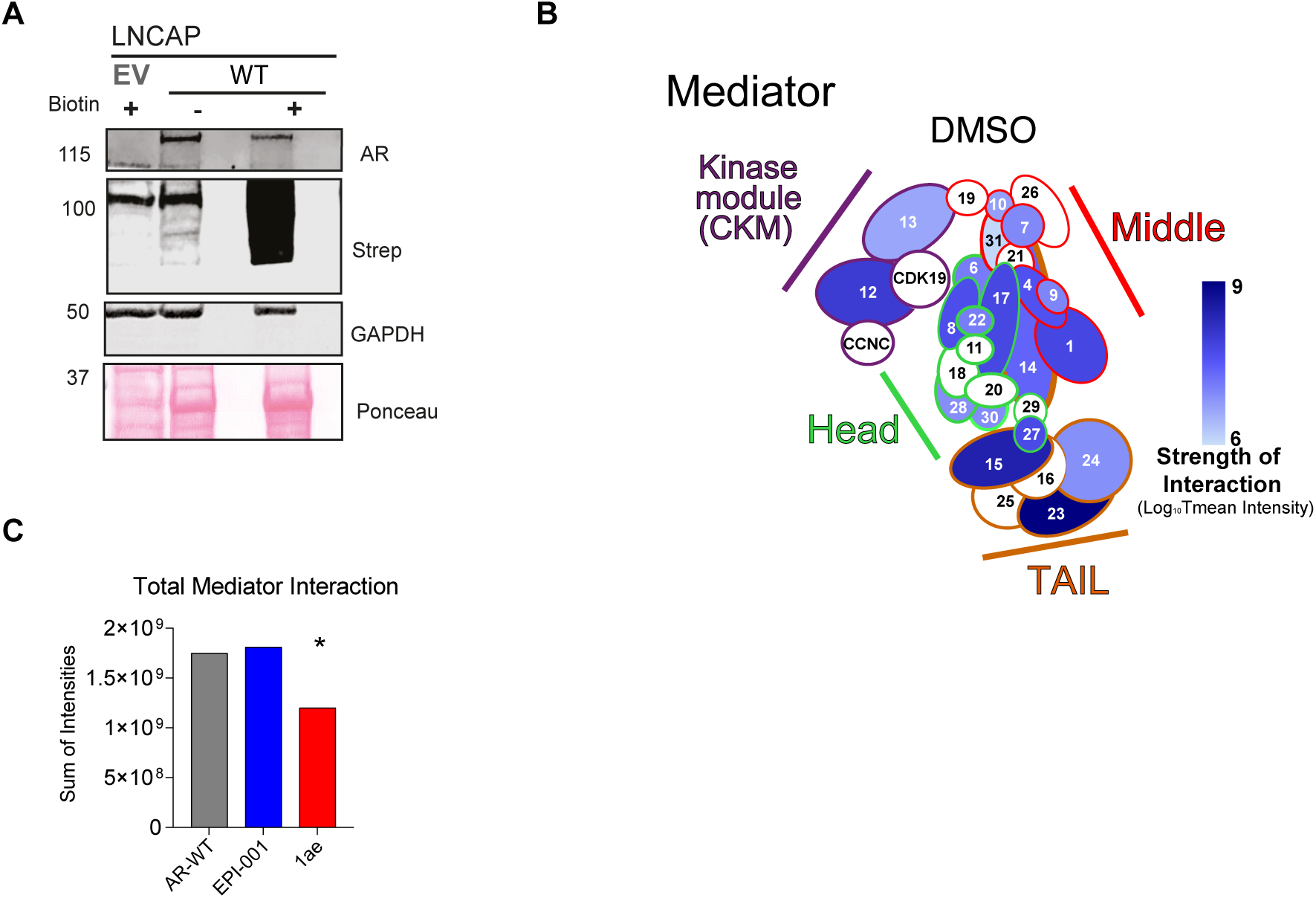
Small molecule inhibitors alter AR proteomic interactions with Mediator. **A)** Western blot showing expression of FLAG-MTID-AR or FLAG-MTID-Y22toS proteins in LNCaP cells with antibodies for AR. Biotin-dependent labeling is shown with Streptavidin antibodies (Strep) and GAPDH and Ponceau staining are shown as loading controls. EV indicates the empty vector expressing FLAG-MTID. **B)** BioID MS of LNCaP MTID-AR-WT interaction with Mediator complex. Colour indicates strength of interaction from FLAG, LogFC_10_ Tmean of intensity. **C)** TMean SAINTq intensity of total mediator interactions were compared across LNCaP MTID-AR-WT with DMSO or treated cells with small molecule inhibitors. Statistical significance was determined by Mann-Whitney U test *P < 0.05.

**Supplemental Figure 7.**
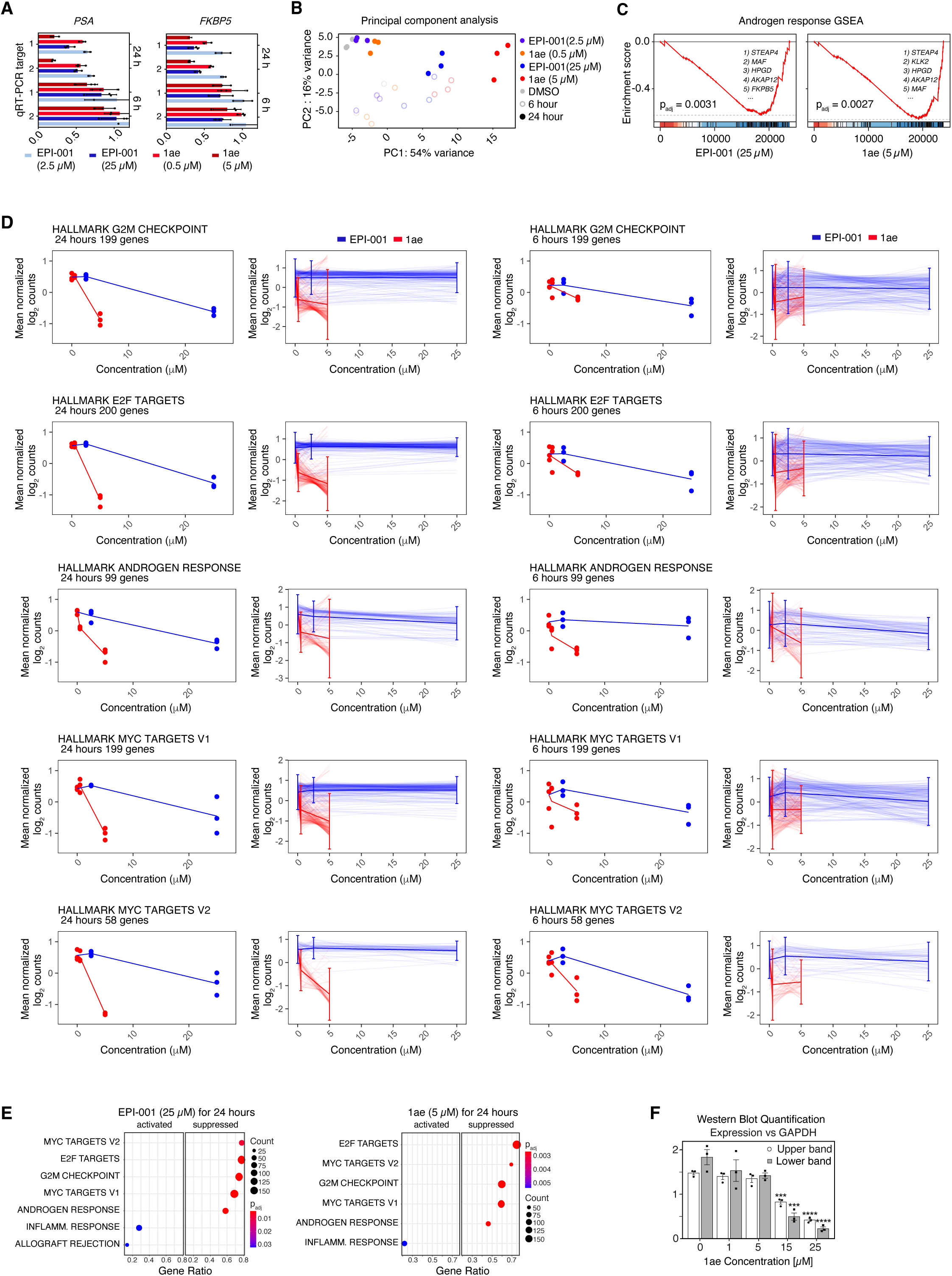
1ae inhibits AR dependent oncogenic pathways in human and mouse models of CRPC. **A)** qRT-PCR of *PSA* and *FKBP5* transcript targets using two primer pairs for each loci, with indicated compound at each time point and concentration used for RNA-sequencing. Values indicate 2^-ΔΔCt^ (Log fold change in target signal signal versus β-Glucuronidase housekeeping gene signal in treatment sample normalized to values from corresponding DMSO control sample) (N = 3). **B)** Principal component analysis of LNCaP cells treated with EPI-001 or 1ae, at indicated time points and concentrations (N = 3). **C)** Random walk of the GSEA running enrichment score of hallmark androgen response pathway genes in LNCaP cells treated with 25 µM EPI-001 or 5 µM 1ae for 24 hours versus DMSO at 24 hours. Top 5 down regulated genes for EPI-001 and 1ae treatment contributing to the leading edge indicated in top right, and adjusted p-value of GSEA statistic indicated in bottom left (N = 3). **D)** Line plots of mean normalized, log transformed read counts of significantly depleted gene sets in LNCaP cells treated with 25 µM EPI-001 or 5 µM 1ae versus DMSO at 24 hours (shown in Fig. 6E), as a function of compound concentration at 6 and 24 hours. Light lines represent individual genes, dark lines represent average of all genes, and bars represent standard deviation (N = 3). **E)** GSEA analysis of RNA-seq experiment showing most significantly activated and suppressed pathways for 25 µM EPI-001 and 5 µM 1ae treatment vs. DMSO at 24 hours, ranked by the adjusted p-value (p_adj_). Gene pathways split by ‘activated’ or ‘suppressed’ based on GSEA enrichment in the gene list ranked by Log_2_FC vs DMSO, in order of gene ratio (detected genes / all genes in pathway) of the analyzed pathway. Circles scale to the count of detected genes from the analyzed pathway, and color scales to p_adj_ from the analyzed pathway. (N = 3).

## Supplementary Tables

**Supplementary Table 1 |** BioID-MS data corresponding to Fig. 2 and Supplementary Fig. 3. The Bait tab contains the bait IDs used for SAINTq analysis and the SAINTq output tab is the analysis output. The Data summary is a pivot table generated from the output data. The “WT Top 75 GO” and “Y22S Top 75 GO” tabs are the output data from STRING analysis of the top 75 most abundant proteins with a BFDR ≤ 0.02 and a FC ≥ 3 from the t_DHT_ = 60 minute samples.

**Supplementary Table 2 |** BioID-MS data corresponding to Fig. 5 and. The Bait tab contains the bait IDs used for SAINTq analysis and the SAINTq output tab is the analysis output. The Data summary is a pivot table generated from the output data. Of fold change of change in small molecule inhibitor vs DMSO LNCaP MTID-AR-WT cells. GO tab refers to most depleted proteins vs DMSO with a p ≤ 0.05 and a FC ≥ -1.5.

**Supplementary Table 3 |** RNA-Seq DESeq2 Log_2_FC values by contrasts indicated in Fig. 6 and Supplementary Fig. 7.

**Supplementary Table 4 |** RNA-Seq DESeq2 raw count matrix and normalized count matrix used to calculate expression values plotted in Fig. 6 and Supplementary Fig. 7D.

## Supplementary Videos

**Supplementary Video 1 |** Fluorescence Time-Lapse Video of eGFP-AR condensates in PC3 cells. Cells were treated with 1 nM DHT and imaged with spinning disk microscopy. Scale bar: 10 µm.

**Supplementary Video 2 |** Fluorescence Time-Lapse Video of eGFP-AR-ΔNLS condensates in PC3 cells. Cells were treated with 1 nM DHT. Scale bar: 10 µm.

**Supplementary Video 3 |** Fluorescence Time-Lapse Video of PC3 cells expressing eGFP-AR or the indicated YtoS mutant. Cells were treated with 1 nM DHT and imaged with spinning disk microscopy. Scale bar: 10 µm.

## Supplementary Documents

**Supplementary Document 1 |** Chemical synthesis.

## Notes

### Summary of Updates

Modified title, abstract, order of presentation of the data. Removed data, added data and edited author list.

